# The integrin-mediated adhesome complex, essential to multicellularity, is present in the most recent common ancestor of animals, fungi, and amoebae

**DOI:** 10.1101/2020.04.29.069435

**Authors:** Seungho Kang, Alexander K. Tice, Courtney W. Stairs, Daniel J. G. Lahr, Robert E. Jones, Matthew W. Brown

## Abstract

Integrins are transmembrane receptor proteins that activate signal transduction pathways upon extracellular matrix binding. The Integrin Mediated Adhesion Complex (IMAC), mediates various cell physiological process. The IMAC was thought to be an animal specific machinery until over the last decade these complexes were discovered in Obazoa, the group containing animals, fungi, and several microbial eukaryote lineages. Amoebozoa is the eukaryotic supergroup sister to Obazoa. Even though Amoebozoa represents the closest outgroup to Obazoa, little genomic-level data and attention to gene inventories has been given to the supergroup. To examine the evolutionary history of the IMAC, we examine gene inventories of deeply sampled set of 100+ Amoebozoa taxa, including new data from several taxa. From these robust data sampled from the entire breadth of known amoebozoan clades, we show the presence of an ancestral complex of integrin adhesion proteins that predate the evolution of the Amoebozoa. Our results highlight that many of these proteins appear to have evolved earlier in eukaryote evolution than previously thought. Co-option of an ancient protein complex was key to the emergence of animal type multicellularity. The role of the IMAC in a unicellular context is unknown but must also play a critical role for at least some unicellular organisms.

## INTRODUCTION

Integrins are transmembrane signaling and adhesion heterodimers, made up of α-integrin (ITGA) and β-integrin (ITGB) protein subunits [1]. They are important signal transduction molecules across the cell membrane and associate with a set of intracellular proteins that act on the actin cytoskeleton, together known as the integrin mediated adhesion complex (IMAC) (figure 1A) [2]. The intracellular component of the IMAC includes adhesome proteins that are 1) integrin bound proteins that can bind to actin directly (talin, α-actinin, and filamin); 2) integrin bound proteins that can bind indirectly to the cytoskeleton (integrin linked kinase (ILK), particularly interesting new cysteine-histidine rich protein (PINCH), and Paxillin); and 3) non-integrin binding proteins such as vinculins.

**Figure 1:**
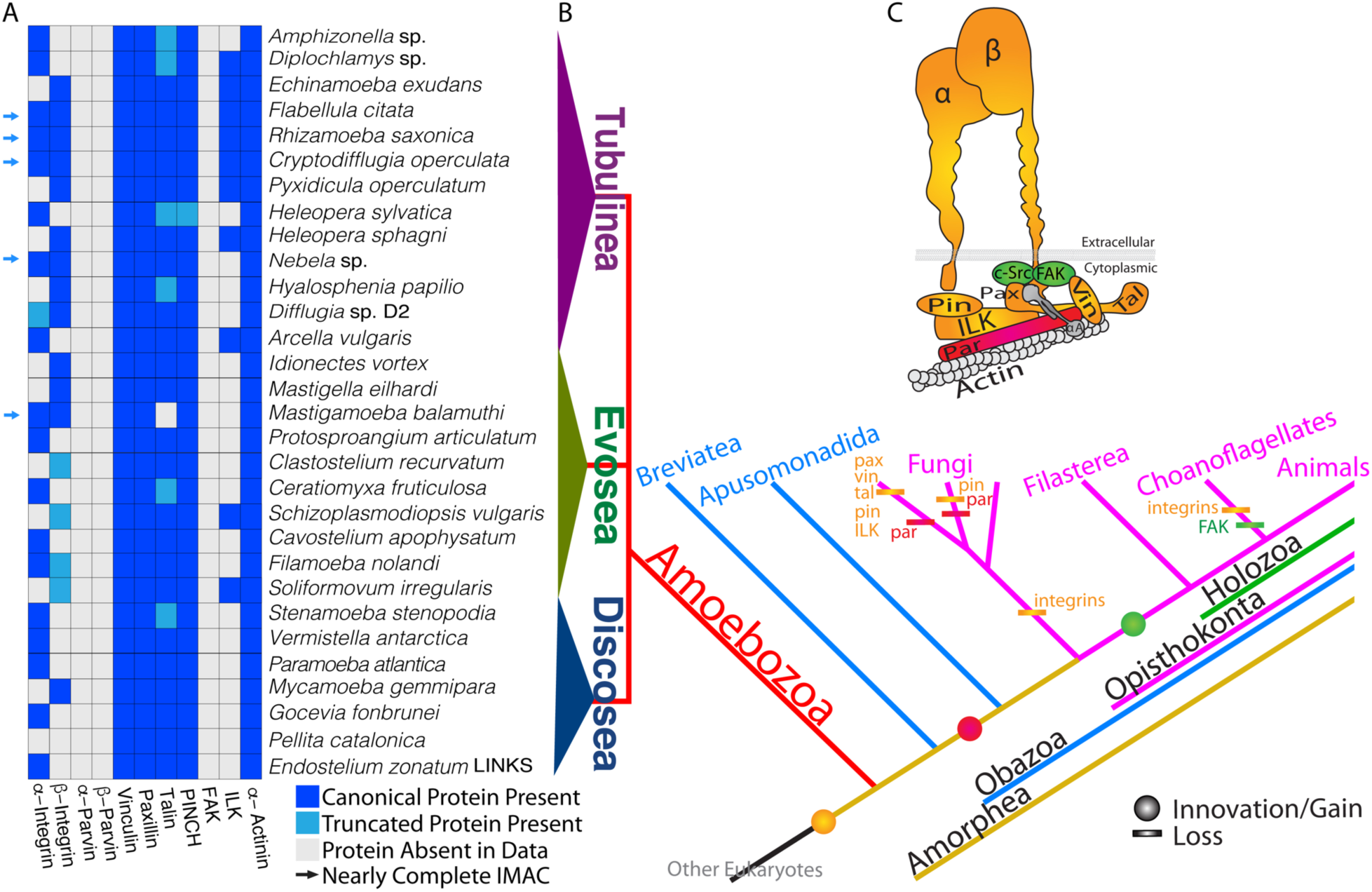
A) Repertoire of IMAC in amoebozoan species that have either integrin alpha or beta. Names of IMAC proteins are listed at the bottom and names of amoebozoan taxa are listed on side of the presence/absence heat map. Arrows indicates amoebozoan species that have nearly complete set of IMAC proteins. Dark blue indicates the presence of a canonical IMAC protein, white indicates the absence of an IMAC protein and light blue indicates a truncated form of IMAC protein. B) Phylogenetic distribution of IMAC. Circle, innovation/gain; bar, loss. Abbreviations: vin, vinculin; pax, paxillin; tal, talin; par, parvin; pin, particularly interesting new cysteine-histidine-rich protein; FAK, focal adhesion kinase; ILK, integrin-linked kinase; αA, alpha-actinin; α, α-integrin; β, β-integrin. C) Cartoon schematic of the membrane docked IMAC, which associates with intracellular actin (modified from Brown et al., 2013), color corresponds with the species tree.

The IMAC is best known for its critical role in cellular aspects of behavior including development, migration, proliferation, survival and metabolism [1,3,4]. They function by relaying messages through outside/in and inside/out between the extracellular matrix and the internal cytoskeleton[5].

Since integrin proteins have been demonstrated to be absent in other complex multicellular lineages of eukaryotes (i.e., plants and fungi) as well as in the choanoflagellates, which are the closest protistan relatives of Metazoa, integrins were initially believed to be an animal specific evolutionary innovation. However, over the last decade, investigations into the genomic content of other close protistan relatives of animals have led to the discovery of integrins and other IMAC components outside of Metazoa. Integrins and complete components of the IMAC have been reported outside Metazoa in unicellular opisthokonts, such as *Capsaspora owczarzaki*[6], *Pigoraptor* spp.[7], *Syssomonas multiformis*[7], *Corallochytrium limacisporum*[8], *Creolimax fragrantissima*[9], *Sphaeroforma arctica*[10], as well in the apusomonad *Thecamonas trahens* [6], and breviate *Pygsuia biforma*[11]. Thus, the IMAC complex and other associated proteins have a much more ancient evolutionary origin than previously expected, well outside of the animals, but IMAC remains to be only reported the in lineage called Obazoa, which is composed of Opisthokonta, Breviatea, and Apusomonadida [11]. However, are these complexes exclusive to Obazoa, or have they appeared even deeper in evolutionary history?

Amoebozoa is the eukaryotic supergroup that is sister to Obazoa, altogether grouped in the Amorphea[12]. Despite the relatively close evolutionary proximity of Amoebozoa to the animals containing lineage, genomic-level data and relevant research into the supergroup remains sparse. The predicted or inferred genomic complement present has yet to be undetermined in the last common ancestor of Amoebozoa. Recent works revealed that in Amoebozoa (*Dictyostelium discoideum* and *Acanthamoeba castellanii*) crucial components of the IMAC were missing [13], namely ITGA and ITGB. Was this result simply due to a lack of data or is it an evolutionary trend? Here, we examine the gene repertories across the Amoebozoa supergroup to investigate the evolutionary history of the adhesome associated with the integrin proteins. The absence of the integrin proteins reported previously may simply be due to the great paucity of taxonomically broad genomic data from this clade, which is a limiting factor in our knowledge of IMAC evolution.

To test if the lack of IMAC is truly absence or rather unobserved due to a lack of data from a broad taxonomic sampling, we took a comparative genomic/transcriptomic approach utilizing data recently used to resolve the tree of Amoebozoa [14]. Here, we have found a number of predicted IMAC proteins across a wide breadth of Amoebozoa and in some unexamined obazoans (*Ministeria vibrans* (Opisthokonta [12]) and *Lenisia limosa* (Breviatea[15])), including integrins, both ITGA and ITGB (figure 1B). Across the breadth of these taxa, we have identified a nearly complete IMAC. Here we demonstrate that IMAC is not exclusive to Obazoa. Rather it was in the last common ancestor of Amorphea (i.e., Obazoa + Amoebozoa), which from our interpretations of the gene distributions probably had a full set of these components (figure 1C).

## BACKGROUND

### Molecular details on Integrin proteins

The overall architecture of both integrin proteins includes a large extracellular N-terminal ligand-binding head domain and C-terminal stalk (also termed “legs”). Integrin activation is dependent on the binding of a ligand (such as extracellular matrix proteins (collagen, fibronectin, and laminin) [5]and the binding of divalent cations (primarily Ca^2+^, Mg^2+^, or Mn^2+^) on unique but well-conserved cation binding motifs[16–19]. In animals, there are universally five cation binding sites on the ITGA and three on the ITGB paralogs[16,17,19,20]. Intracellular integrin engagement with cytoskeletal proteins in IMAC can regulate cell-signaling pathways that control differentiation, survival, and motility making the integrins one of the most functionally diverse family of cell adhesion molecules [5,21].

### Integrin Alpha (ITGA)

#### Metazoa ITGA

In metazoan ITGA, which we refer herein to as “canonical ITGA proteins”, there is a β-propeller made of seven blades (IPR013519)[18], an integrin leg domain (IPR013649)[18], a transmembrane domain, and a C-terminal cytoplasmic tail amino acid motif (KXGFFXR - IPR018184) [22](numbers are InterPro ids derived from Interproscan version 5.27.66 (IPRSCAN)[23]). These proteins are typically 700-800 amino acids in animals. The β-propeller blade motif is composed of an FG-GAP domain (IPR013517), and in the final 3-4 blades there is a cation binding site motif between an “FG” and the “GAP” motifs. Of note, the consensus pattern of the FG-GAP domain is highly variable[20], and not necessarily those amino acids (i.e., FG and GAP)[20]. The cation binding blades of the β-propeller influences extracellular ligand affinity and ligand specificity[24,25]. The cation motifs follow the consensus pattern of D/E-h-D/N-x-D/N-G-h-x-D/E (h=hydrophobic residue; x=any residue)[16]. In metazoan ITGA, there are three ITGA leg domains (IPR013649) located next to the ITGA head. These ITGA leg (IPR013649) domains are structurally and functionally similar to immunoglobulin (Ig) fold-like beta sandwich[18].

The classification of an authentic ITGA protein outside Metazoa would be a membrane docked protein with an ITGA head composed of a β-propeller made of several β-propeller blades (IPR013519) (figure 2), some with FG-GAP domains (IPR013517) only, and some with a cation motif flanked by FG and GAP motifs (figure 4). However, we do not discard the possibility of truncated ITGA because they occur in metazoans [26,27] and in the opisthokont *Creolimax fragrantissima* [9].

**Figure 2:**
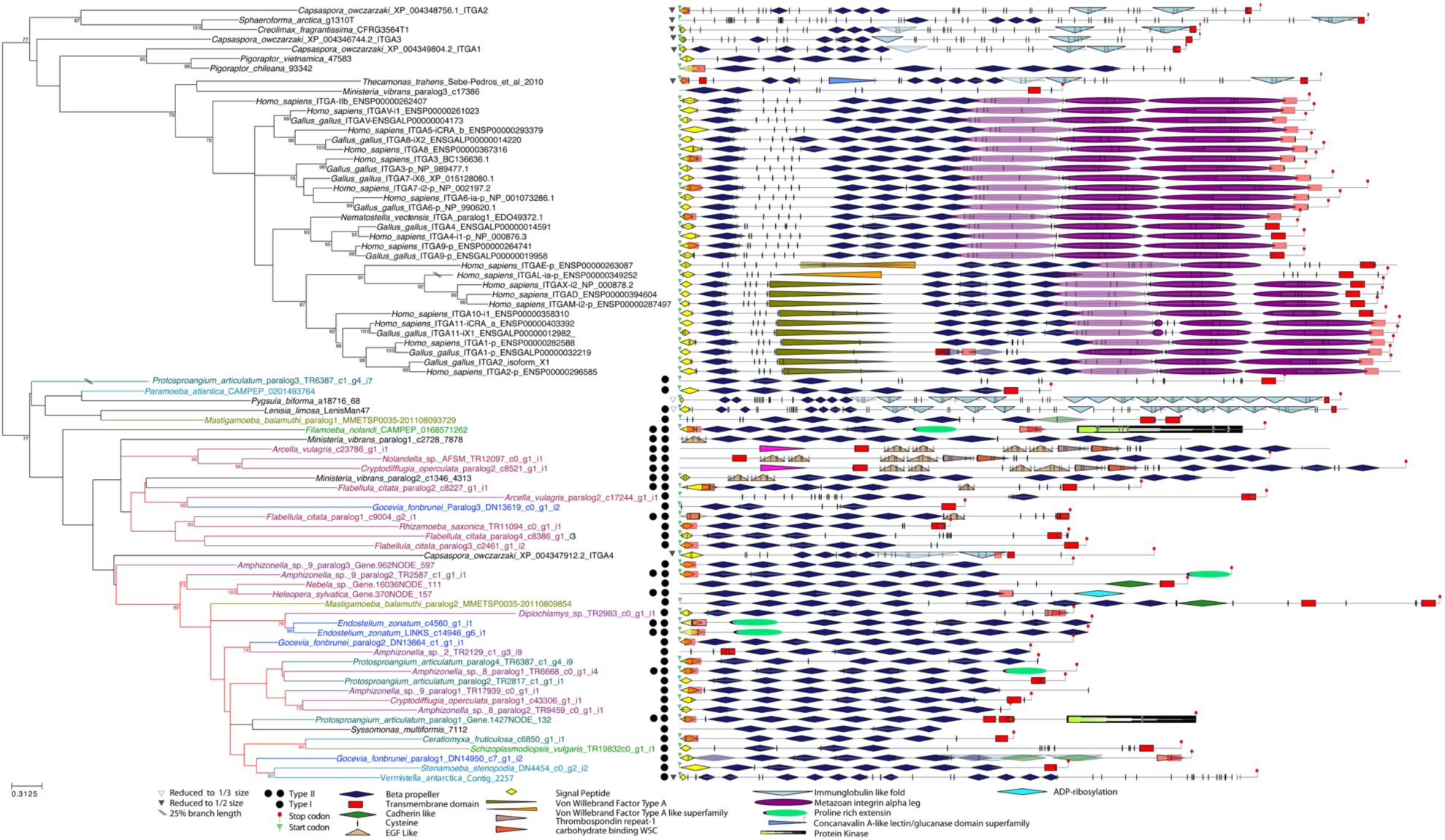
Domain architecture of ITGA from Amoebozoa. The domain architecture of ITGA for amoebozoan species are shown at the right side and phylogenetic tree of Amoebozoa is shown at the left side of figure. Keys to each domain’s color and shape are represented at the bottom of the figure. Phylogenetic tree of amoebozoan ITGA, 160 sites phylogeny of amoebozoan ITGA rooted with Obazoa ITGA. The tree was built using IQTREE v1.5.5 under the LG+C60+F+G model of protein evolution. Numbers at nodes are ML bootstrap (MLBS) values derived from 1000 ultrafast MLBS replicates. Any values that are below 50% are not shown. The color of branches amoebozoan sequences are illustrated according to Kang et al figure 3[14]. When domains are overlapped because of different software algorithms, these are represented in transparent colors.

**Figure 3:**
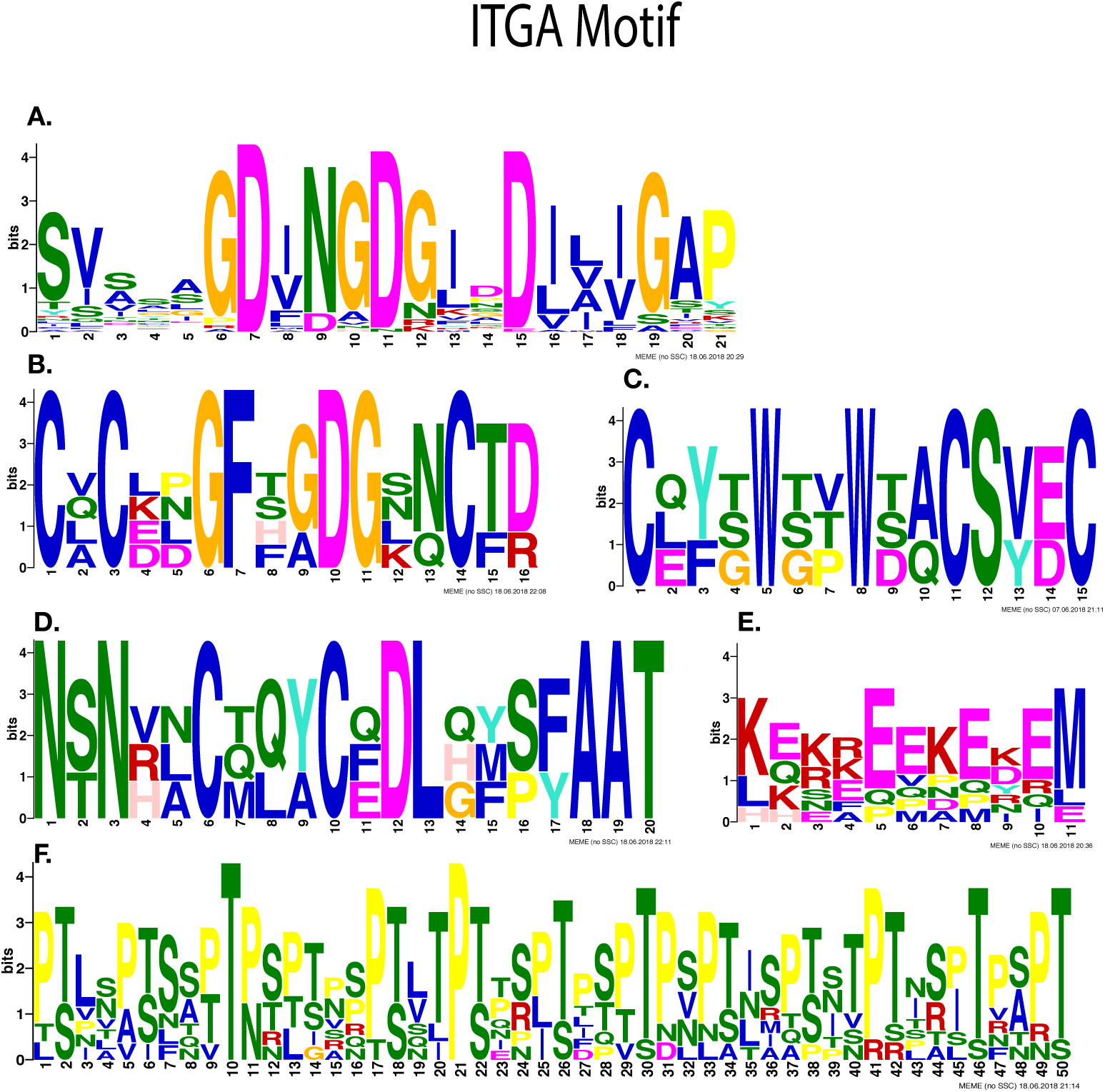
Most significantly enriched motifs in 23 amoebozoans ITGA homologs, as obtained by MEME-suite except for (**E**). (**A**) Consensus cation motif and FG-GAP motifs found in 23 amoebozoans ITGA homologs. (B-D) For figure 3B to 3D, these consensus motifs are exclusively enriched in three amoebozoan ITGA homologs out of 23 amoebozoan ITGA homologs: (**B**) EGF-like motif (**C**) Thrombospodin repeat 1 motif (**D**) Carbohydrate WSC motif. (**E**) GFFKR motif found in three ITGA homologs from *Filamoeba nolandi* and two obazoans species, *Ministeria vibrans* and *Pygsuia biforma*. (**F**) Proline-rich extensin motif found in 8 ITGA amoebozoan homologs out of 23 amoebozoan ITGA homologs.

**Figure 4:**
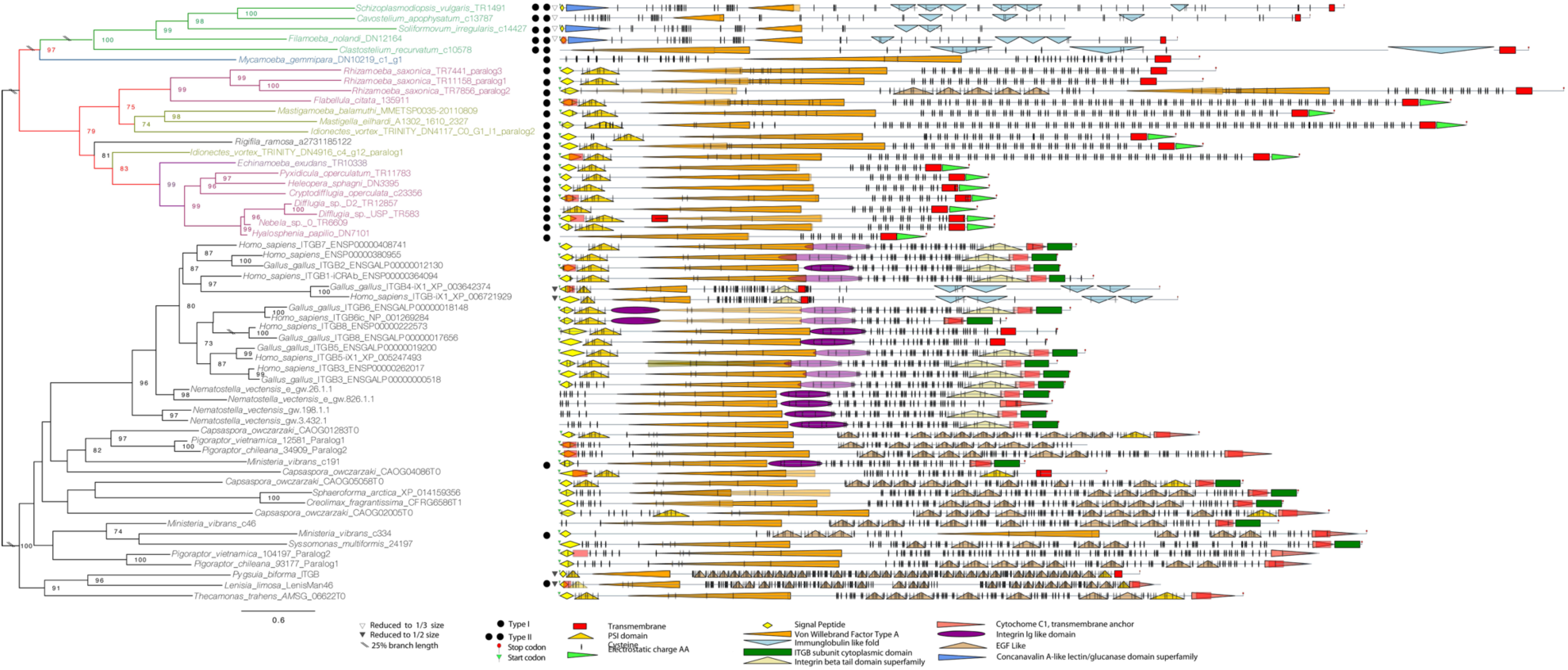
Phylogenetic tree of ITGB homologs with a focus on Amoebozoa. The domain architecture of ITGB homologs are shown to the right and the phylogenetic tree of ITGB. Keys to each of domains color and shape are represented in the box at the bottom of the figure. Tree of amoebozoan ITGB 295amino acids sites phylogeny of amoebozoan ITGB rooted with Obazoa. The tree was built using IQTREE v1.5.5 under the LG+C60+F+G model of protein evolution. Numbers at nodes on the phylogenetic tree are maximum likelihood bootstrap support values derived from 1000 ultrafast bootstrap replicates. Any values that are below 50% are not shown. The color of branches amoebozoan sequences are illustrated according to Kang et al. figure 3[14]. When domains are overlapped because of different software algorithms, these are represented in transparent colors.

### Integrin Beta (ITGB)

#### Metazoan ITGB

In metazoan ITGB, which we refer herein to as “canonical ITGB proteins”, there are seven domains; a plexin-semaphorin-integrin (PSI) (IPR002165), a hybrid domain (described below), three to four EGF modules (IPR013111), an integrin β tail subunit (IPR012896), transmembrane, and signal peptide domains[18]. The metazoan hybrid domain is composed of a unique von Willebrand factor A (known as ITGB “inserted” domain (or β-I) (IPR002369) and a Ig-like domain (IPR013783) upstream of the β-I domain [18] (figure 3). EGF domains make up a cysteine rich stalk motif (CRSM) that corresponds to a β-subunit stalk region[18]. The consensus pattern of the CRSM in Opisthokonta is CXCXXCXC[6,28], in Metazoa there are typically 3-4 CRSMs. The metazoan transmembrane domain consists of GXXXG membrane motif, important for oligomerization of integrin proteins[29].

In Metazoa, the β-I-domain is made up of three metal ion binding sites (comprising a ca. 200 amino-acid region in total)[18]. These are, a metal ion dependent adhesion site (MIDAS), site adjacent to MIDAS (called ADMIDAS), and a synergistic metal ion binding site (SyMBS)[16]. These metal ion binding sites are key regulators for all integrin-ligand interactions and require the divalent cation ions such as manganese, calcium, and magnesium ions[16]. Three metal binding sites have a consensus motif of DXSYSX^95^EX^30^D (at site 150 in Human ITGB1 (NCBI:P05556.2) (MIDAS), SXXDDX^121^DX^83^A (at site 154 in Human ITGB1 (NCBI:P05556.2) (ADMIDAS), and D/EX^55^NDXPXE (at site 189 in Human ITGB1 (NCBI:P05556.2) (SyMBS)[17]. The MIDAS and SyMBS (the positive regulatory site for integrin activation[16]) motifs are essential for integrin-ligand binding[16,18], but ADMIDAS (the negative regulatory [30] site for integrin inhibition) is not absolutely necessary for integrin activation[30]. The ITGB tail subunit (IPR012896) consists of two conserved motifs responsible for activating the integrin complex: the cytosolic tail motif (NPXY/F - to scaffold cytoskeletal proteins) located after the transmembrane region, and the ITGA interacting motif (LLXXXHDRKE – a highly electrically charged amino acid motif) downstream (C-terminus) of the NPXY/F motif, which interacts with scaffolding and signaling proteins[31].

The classification of an authentic ITGB protein outside Metazoa would be a membrane docked protein with an ITGB head composed of a von Willebrand factor A (IPR002369) with three metal ion binding sites, and cysteine rich regions at downstream (N-terminal) (figure 3). Again, we do not discard the possibility of truncated ITGB since these are present in both metazoans [26,27] and *Creolimax fragrantissima*[9].

## RESULTS

### Integrin Alpha

#### Amoebozoan ITGA: *General observations on ITGA*

Our results show ITGA homologs are found in 23 out of 113 amoebozoan transcriptomes examined (figure 2). Of those 23 amoebozoans, eight have more than one ITGA paralog (figure 2). Amoebozoan ITGAs vary greatly in size ranging from 523-1200 amino acids in length (supplemental table 1). Many of the proteins we identified are predicted to have a diverse number of β-propellers from a few (3-5 β-propeller blades), to canonical (6-7 β-propeller blades), to many (more than 8 β-propeller blades) and cation binding sites (1 to 5 sites). However, all amoebozoan ITGA proteins have at least three β-propeller blade (IPR013517) domains (figure 2), with an FG-GAP domain and FG-GAP/cation binding motif (figure 3A). The positions of the cation binding motif/FG-GAP domains are seemingly distributed randomly (supplemental table 1), and not necessarily in the last 3-4 blades as in canonical animal ITGA. We noticed the presence of a C-terminal transmembrane anchor (IPR021157) domain in three amoebozoan ITGAs. Animal-like KXGFFXKR cytoplasmic tail motif is absent in all but one amoebozoan (figure 3E; supplemental table 1).

We find amoebozoan ITGAs are present in all three major clade, but interestingly these proteins are not in eight amoebozoan taxa that have genome sequences, (*Dictyostelium* spp., *Acytostelium* spp., *Heterostelium* spp., *Speleostelium caveatum, Physarum polycephalum, Entamoeba* spp., *Acanthamoeba* spp., and *Protostelium fungiovrum*[32]). However very importantly, we were able to confidently find ITGA in *Mastigamoeba balamuthi* (figure 2), which has a complete genome available [33]. To confirm the presence of these genes in the genome, we used a BLAST-based homology search to identify 2 ITGA genes of *M. balamuthi* to the genome data (CBKX00000000) on NCBI (see Methods) [86]. For ITGA, there are no introns in either paralog 1 (CBKX010025823.1) or paralog 2 (CBKX010005624.1).

We classified ITGA into two representative types based on their predicted IPRSCAN domains, cation motifs, their similarity to metazoans ITGA, and novel domains that are previously unobserved in ITGA proteins.

#### Amoebozoa ITGA: *Type I ITGA: similar architecture to Metazoa*

For communication purposes we have classified the amoebozoan ITGA homologs as ‘Type I’ and ‘Type II’ based on their similar or divergent domain complements compared to metazoan ITGA sequences, respectively (figure 2). From the 23 amoebozoan taxa in which we identified an ITGA, 19 taxa have Type I (figure 2). One amoebozoan (*Amphizonella* sp. 9 paralog 2) has a Type I ITGA with a domain architecture and cation motifs that are virtually identical to head region of metazoan ITGA (figure 2, supplemental table 1). The rest of the amoebozoan Type I ITGAs have canonical beta propellers, but they have varying numbers of cation motifs that differs from the canonical ITGA proteins (supplemental table 1). Most of the cation motifs of all but four amoebozoan Type I ITGAs follow the consensus metazoan amino acid pattern (figure 3A, supplemental table 1). thirteen of the amoebozoan Type I ITGAs have both a predicted signal peptide and transmembrane region (figure 2). The rest of the amoebozoans Type I ITGAs have only a predicted signal peptide region or transmembrane region. Amoebozoa ITGA without transmembrane region may be the result of endogenous soluble integrin. In a few cases we did not observe the signal peptide or the transmembrane region. This may be due to truncation as we are predicting proteins from a transcriptome or the product of shedding [34]. However, in the genome of *Mastigamoeba balamuthi* all ITGA paralogs lack signal peptides and one lacks a transmembrane region (paralog 1).

Out of 19 Type I amoebozoan ITGA homologs, four ITGAs have an extracellular Ig-fold-like (IPR013783) or cadherin-like (IPR015919) domain next to the ITGA head (figure 2). A cadherin like domain (IPR015919) and an Ig-fold-like domain (IPR013783) are part of an immunoglobulin fold like beta sandwich protein class, which fold in a similar structure[35]. The three amoebozoan ITGAs that contain either an Ig-fold-like or a cadherin-like domain also have canonical β-propellers (6-7 blades) (figure 2), although two of the four Type I ITGA have an extra cation motif (supplemental table 1).

#### Amoebozoan ITGA is a Type II ITGA with novel domains

We designate any ITGA homolog with a domain flanking the ITGA head that has not previously been reported in an ITGA protein as a “Type II” ITGA (figure 2). From the 23 amoebozoan taxa in which we identified an ITGA, 12 taxa have Type II (figure 2). Of the 23 Type II amoebozoan ITGAs we discovered, two have a protein kinase domain (IPR000719) (figure 2), five amoebozoans have a proline rich extension motif called “extensins” (PR01217) (numbers are SPRINTS ids derived from Prints version 42.0 [36] and Interproscan version 5.27.66 [23](supplemental methods)) (figure 3F; supplemental table 1).

In one Type II ITGA, one amoebozoan sequence (*Flabellula citata* paralog 1) uniquely has an epidermal growth factor (EGF)-like domain (IPR000742) as its only novel domain located downstream (C-terminal) to the ITGA head (figure 2). Additionally, three amoebozoans uniquely have two epidermal growth factor (EGF) domains (figure 3B), a Thrombospondin repeat-1 (TSP1) (IPR000884) (figure 3C), and a carbohydrate binding domain called a wall stress component (WSC) (IPR002889) (figure 3D) all with cysteine rich motifs located upstream (N-terminal) to the ITGA head (figure 2). We were unable to detect any other protein in the *nr* database with significant similarity to the N-terminal domain composition of these sequences (EGF-TSP1-WSC) using an e-value cut-off < 1e-10. This suggests that this EGF-TSP1-WSC architecture may be unique to these three amoebozoans.

Seven of the amoebozoan ITGAs we designated as Type II possess canonical (6-7 blades) β-propellers (figure 2) and a different number of cation motifs (supplemental table 1). The other five amoebozoans Type II ITGAs have short (3-5 blades) β-propellers, and a wide range of number of cation motifs (supplemental table 1). Four amoebozoan ITGA homologs of this type contained both a signal peptide and a transmembrane domain (figure 2). Eight others had either a signal peptide or a transmembrane region.

#### Obazoan ITGA: *General observations on ITGA*

We also found ITGA homologs in several underexplored obazoan transcriptomes: the opisthokont *Ministeria vibrans* and the breviate *Lenisia limosa* (figure 2). Obazoan ITGA proteins have at least three β-propeller blade (IPR013517) domains, with an FG-GAP domain (figure 2, supplemental figure 5A) and FG-GAP/cation binding motif domains are seemingly distributed randomly (supplemental table 1). Animal-like KXGFFXKR cytoplasmic tail motifs were observed in most obazoan transcriptomes that were examined in this study (figure 4E, supplemental figure 5C). Among the non-animal obazoans, the seven novel obazoans ITGA have an Ig-fold-like domain leg next to ITGA head (figure 2; supplemental figure 5D).

*Ministeria vibrans* has both Type I and Type II paralogs (figure 2). In Type II, paralog 1 and 2 have a predicted long (10) ITGA β-propeller, but they do not contain transmembrane domains. The other paralog has a few (3) β-propeller blades (one with FG-GAP/cation binding motif), a cytosolic tail motif (figure 3E), and a transmembrane domain (figure 2). In both paralog 1 and 3, a signal peptide region was not predicted. *Ministeria vibrans* Type II ITGA uniquely has an *epidermal growth factor* (EGF) like domain (IPR000742) attached upstream (N-terminal) to the ITGA head (figure 2). All obazoans in our study have a signal peptide or a transmembrane region except for *Pigoraptor* spp., *Sphaeroforma arctica*, and *Syssomonas multiformis* (figure 2), which lack both.

#### Phylogenetic analysis of ITGA

The unrooted tree of ITGA recovered all of Amoebozoa ITGA as a clade (figure 2), which is generally sister to most of Obazoa. However, a few opisthokont sequences are nested in this clade. These include a paralog of *Capsaspora owczarzaki* (ITGA4: XP_004347912), both *M. vibrans* paralogs, Breviates and the lone *Syssomonas multiformis* ITGA. The major lineages or sublineages of Amoebozoa were not recovered. Instead amoebozoan ITGA are divided into two clades. One clade consists primarily of tubulinid amoebozoans (+ *Ministeria*) and the other contains the rest of Amoebozoa.

It should be noted that in addition to these taxa, the genome sequence of the very evolutionary distant Stramenopile brown alga (*Ectocarpus siliculosus*) revealed the presence of three ITGA homologs[37] (supplemental figure 8). However, no other available Stramenopile genome data or transcriptomic data searched thus far (for a list of taxa searched, please see supplemental text) contains integrin homologs in our examinations. Therefore, we inferred an additional ITGA tree with this species (supplemental figure 3). Through BlastP to NCBI-NR we identified several predicted beta propeller proteins in Archaea that have a similar architecture to the canonical ITGA. An unrooted tree of beta propeller homologs recovered Archaea as a paraphyletic across our phylogeny (supplemental figure 8), this is further discussed below.

We observed more FG-GAP proteins in other taxa such as cyanobacteria (e.g. *Gloeobacter violaceus*; WP_011143221.1, *Nostoc punctiforme*; ACC84844.1 and *Crocosphaera watsonii*; WP_007307935.1), Haptophyta (*Emiliania huxleyi*; XP_005778208.1), Crytophyta (*Guillardia theta*; XP_005842481.1), and Stramenopiles (e.g. *Nannochloropsis* spp. CCMP1776: TFJ84058.1, EWM21519 and *Cafeteria roenbergensis*; KAA0148838.1) on NCBI. These FG-GAP proteins either do not possess transmembrane and signal peptide regions or possess 7 transmembrane or more. Thus, in our estimation these FG-GAP proteins fail to pass our minimum criteria of being an ITGA.

### Integrin Beta

#### Amoebozoan ITGB: General observations on non-metazoan ITGB

We find that ITGB homologs are found in 19 out of 113 amoebozoan transcriptomes examined with three ITGB paralogs in *Rhizamoeba saxonica* and two ITGB paralogs in *Idionectes vortex* (figure 4). We find amoebozoan ITGBs are present in all three major clades of Amoebozoa. However like in ITGA, these proteins are not in four amoebozoans groups that have genome sequences, (the dictyostelids, entamoebids, acanthamoebids, and *Protostelium fungiovrum*[32]). We were able to find ITGB gene in the genome of the amoebozoan *Mastigamoeba balamuthi* ATCC 30984 (figure 6). For ITGB, there were five introns in the genomic scaffold CBKX010020319.1 and eight introns in CBKX010020318.1. These introns are canonical *M. balamuthi* introns and the other genes on the scaffold are mastigamoebid/amoebozoan in nature. We detected a tenascin ORF on ITGB genomic scaffold of CBKX010020318.1 and PA14 ORF on CBKX010020318.1. We inferred a phylogenetic tree for tenascin and PA14 by searching for homologs in our extensive datasets, both of which clearly show amoebozoan signal (shown as a subset in figure 6). The presence of introns within the genomic scaffold for ITGB clearly demonstrates that this protein is indeed part of the genome. Additionally, the phylogenetic signal of other ORFs on these genomic scaffolds show amoebozoan signal and are not from contamination in the genome project.

**Figure 5:**
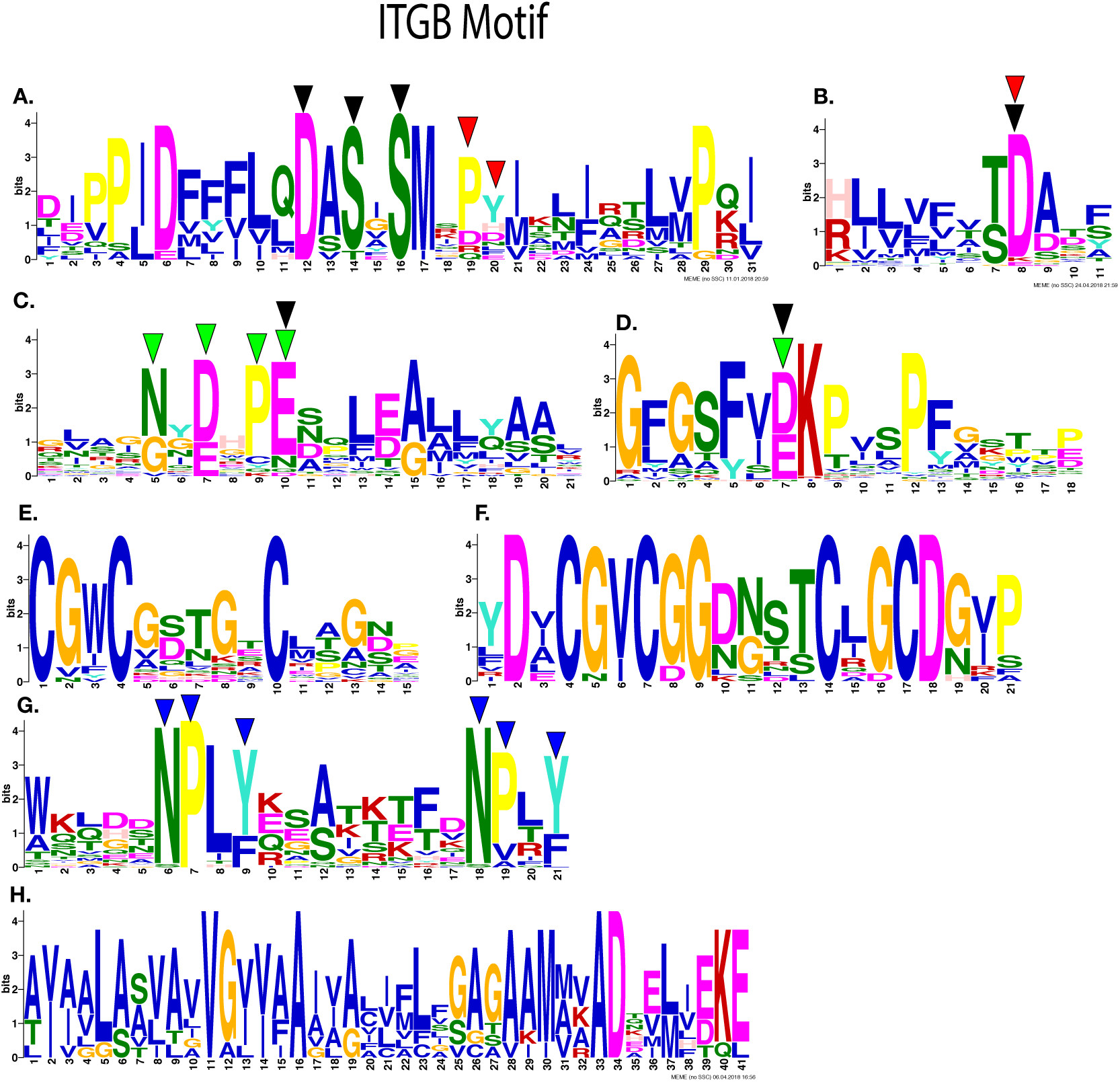
Most significantly enriched motifs in 18 amoebozoans ITGB homologs, as obtained by MEME-suite. (**A**)-(**F**) Three divalent cation binding motifs: MIDAS, ADMIDAS and SyMBS are shown. (**A**) MIDAS (black arrow) and ADMIDAS (red arrow) motifs are shown. (**B**) A shared amino acid (indicted by arrow) in MIDAS and ADMIDAS motifs are illustrated. (**C**) SyMBS motif represented by green arrow and a shared amino acid of MIDAS motif indicated by black arrow are shown. These consensus motifs are obtained from type I amoebozoan ITGB homologs sequences only. (D) SyMBS motif indicted by a green arrow is shown. (**E**-**F**) Cysteine rich motifs are shown. (E) PSI domain and (F) Cysteine-rich motif stalk is shown. These are recovered from type I amoebozoan ITGB sequences. (**G**-**H**) Motifs found at the C-terminal are shown. (G) NPXY motif shown in blue arrow. found in most amoebozoans ITGB except three amoebozoan ITGB homolog. (H) An electrostatic charge motif is shown, this motif is found exclusively in type I ITGB homologs.

**Figure 6:**
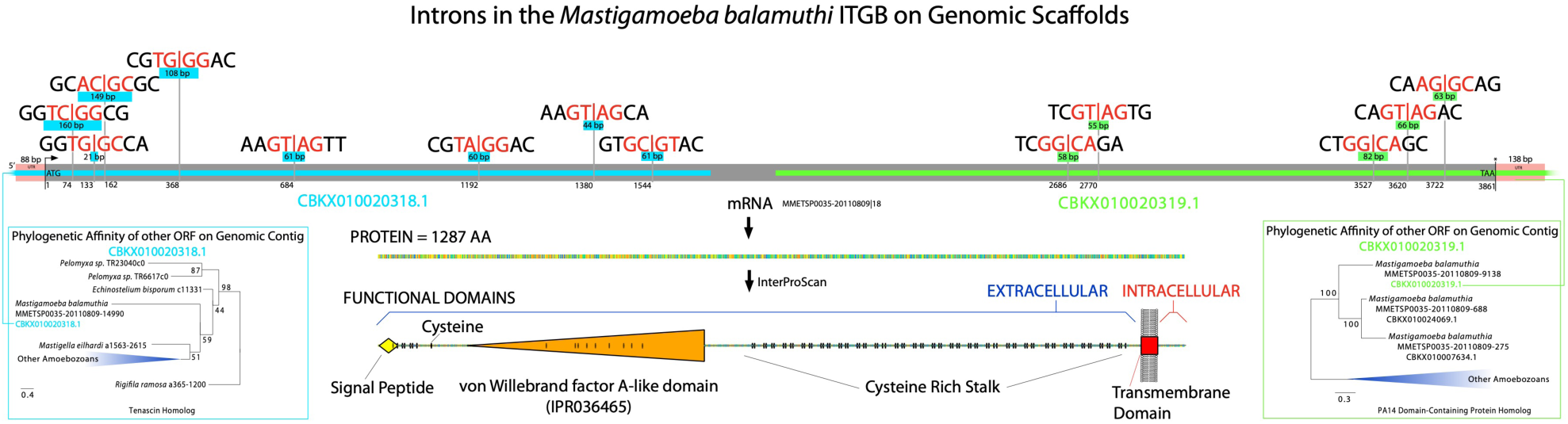
The presence of ITGB gene in a *Mastigamoeba balamuthi* genome sequence. Introns/exons boundaries sequences are shown along with corresponding intron size. The genome contig that possess ITGB gene are colored blue and green. The protein domain architecture of ITGB gene are shown along with protein size. The adjacent tenascin gene on the ITGB genome contig are shown left and right along with the inferred phylogenetic tree.

The size range of ITGB paralogs in Amoebozoa vary widely from 595 to 3296 amino acids (supplemental table 2). In addition to the novel ITGB proteins in Amoebozoa, we also found a previously unreported ITGB homologs in two obazoan transcriptomes, as well in CRuMs, which is a recently described clade closely related to Amorphea[38]. Based on domain and motif architecture from IPRSCAN outputs, we classify non-metazoan ITGBs into two representatives; ITGB that are similar to a canonical ITGB, and unique domain architecture proteins that are ITGB-like.

#### Type I ITGB: similar architecture to Metazoa

Our first representative type of ITGB has a similar domain architecture to metazoan ITGB (figure 4). From the 19 amoebozoan taxa in which we identified an ITGB, 14 taxa have Type I (figure 4). Most amoebozoans in this class appear to have the vWA domain, β-subunit stalk region and the PSI domains except two amoebozoans that lacks PSI domains and one amoebozoans which possess the duplicated vWA domain (figure 4). Amoebozoa have the CRSM pattern in the β-subunit stalk region of DXCGXCGGX^(3-6)^CXGC (figure 5F) ranging from three to seven times (Supplemental table 2). In of β-tail motifs (IPR012896), the cytosolic tail motif (NPXY/F) is present across Amoebozoa (figure 5G) except one amoebozoan (supplemental table 2) whereas an ITGA interacting motif (LLXXXHDRKE) is absent in Amoebozoa. Twelve amoebozoans in this class possess amino acids with a predicted electrostatic charge where the ITGA interacting motif (LLXXXHDRKE) would be present in metazoan ITGB (figure 4, figure 5H). MIDAS are well conserved especially in amoebozoan β-I domains (figure 5A, B) and SyMBS are well conserved across the amoebozoans (figure 5C, D). However, ADMIDAS motifs are quite variable in Amoebozoa compared to the metazoan consensus pattern (figure 5B, Supplemental table 2). A signal peptide and a transmembrane domain were predicted in most amoebozoan ITGB in this class and three amoebozoans lacking signal peptides (figure 4).

The ITGB proteins of non-metazoan obazoans have domain/motif architectures very similar to a canonical metazoan ITGB (figure 4, supplemental 6). At the B-tail motifs (IPR012896) of Obazoa, the cytosolic tail motif (NPXY/F) is present in all obazoans ITGB homologs (supplemental figure 6G), but we did not observe the ITGA interacting motif (LLXXXHDRKE) in ITGB of Breviatea. We found a metazoan GXXXG motif in three amoebozoan ITGB (supplemental table 2), suggesting that this motif is not animal specific. There are multiple tandem repeats of the opisthokont CRSM pattern in Obazoa ITGB and both breviate taxa have a clear expansion of β-subunit stalk region as previously reported [6–11] (figure 3). All of the obazoan ITGB MIDAS, ADMIDAS, SyMBS motifs were well-conserved (supplemental figure 6A-D). We also investigated a possible “choanoflagellate ITGB gene” that was recently reported to be present in the transcriptome of *Didymoeca costata*[39]. The protein domain architecture has a serine protease flanking vWA, PSI domain. However, it is missing the CRSM, NPXY/F (IPR021157), LLXXXHDRKE (IPR014836) and GXXXG motifs. To our surprise *Rigifila ramosa* (in CRuMs), has a protein that contains all the canonical domains of ITGB such as vWA, amoebozoan CRSM pattern and c-terminal motifs (figure 3) and all typical cation motifs (supplemental table 2). We were unable to find an ITGA homolog in either *Rigifila ramosa* or *Didymoeca costata*.

#### Type two ITGB: unique architecture compared to Metazoa and solely found in the amoebozoan clade Variosea

Our second representative class of ITGB, which we refer to as Type II, are large proteins (1405 aa to 3641 aa) that have novel domain architectures unobserved before in ITGB, which we refer to as Type II. Of the total 19 amoebozoan taxa that have ITGB five amoebozoans have an ITGB assigned to our Type II class (figure 4). Type II amoebozoan ITGB domain architectures include Laminin G3 (LamG3) (IPR013320), Von Willebrand Type D (vWD) (IPR001846), Ig-like fold (IPR013783), EGF-like (IPR013032) and a cyclic nucleotide binding (IPR000595) domain (figure 4. A canonical human ITGB has four EGF domains [16,18] that coincide with the location of Ig regions in amoebozoan ITGB. Three amoebozoan ITGB in this class have an extracellular EGF– like domain at the N-terminus, at the location of the PSI domain (figure 3). LamG3 (CSM) and EGF-like domain are recovered in two amoebozoan ITGB (figure 3). The cation binding motifs MIDAS, ADMIDAS motifs are hypervariable in Type II representatives compared to the metazoan consensus pattern, but SyMBS is well conserved except in *Filamoeba nolandi* (supplemental table 2).

#### Phylogenetic analysis of ITGB

We selected *Homo sapiens, Gallus gallus*, and *Nematostella vectensis* as a selective reference group because integrin proteins are highly conserved among Metazoa. The tree of ITGB reveals a three major clade when we rooted on Opisthokonta as an outgroup (figure 3). We observed three clades of type II amoebozoan ITGB, breviates with apusomonads ITGB, and type I amoebozoan with the CRuMs taxon *Rigifila ramosa* ITGB. Therefore, we do not recover Amoebozoa as a monophyletic group.

Additionally, we investigated the evolutionary history of Sib (*Similar to Integrin Beta*) proteins. Sib proteins is a cell adhesion molecule that are similar to ITGB, originally discovered in *Dictyostelium discoideum* [40]. However, they do not appear to have a canonical ITGB (Supplemental result). Additionally, an ITGB homolog was shown in cyanobacteria, but it contained only one of the ITGB domains[6]. Therefore, we inferred an additional ITGB tree with these SIBA proteins from these species (*Didymoeca costata, Dictyostelium discoideum* and cyanobacteria) (supplemental figure 2). In this tree, a clade of Type II ITGB is placed between the cyanobacteria *Trichodesmium* and the choanoflagellate *Didymoeca*. However, these apparent ITGB homologs do not have obvious ITGB-stalk region or NPXY motifs. While they are recovered well with other ITGB, given the important missing motifs and domains the placement of these in a phylogenetic context alone does not clearly define these as ITGB.

### Scaffold Cytoplasmic proteins

In our comparative transcriptome and genome analysis, we observed the following intracellular scaffold cytoskeletal adhesomes proteins: talin, α-actinin, vinculin, PINCH, paxillin, and ILK throughout Amoebozoa. These six adhesome proteins were have been reported to be present in Amoebozoa[6]. Since then the number of the core consensus adhesomes were expanded[21]. Filamin, kindlin, and tensin are three consensus adhesome proteins and cross-actin linkers that interact with ITGB[21]. We have observed two novel adhesome proteins (filamin, tensin) that were not known in Amoebozoa (Supplemental figure 1). We suspect these cytoskeleton proteins may be critical, and the absence of these genes in some amoebozoan taxa is probably a result low gene coverage in some transcriptomes. Talin, PINCH, filamin, and Paxillin key players in the integrin adhesome, were not present in seven amoebozoan transcriptomes (figure 1B), however their absence is also suspected to be a artefact of low gene coverage in these data. Across the amoebozoan clade, we also noticed a large number of what appear to be truncated talin compared to the rest of adhesome proteins (figure 1B).

ILK is scattered across the Amoebozoa unlike in Apusomonadida and Opisthokonta[6–10]. α-actinin is the only signaling protein that is present in all of the amoebozoan taxa. Consistent with previous reports, we did not detect FAK-cSRC or Parvin in any of Amoebozoa or Breviatea[11]. Since the gene coverage for *Pygsuia* (Breviatea) is high and genomes are available for *Lenisia limosa* (Breviatea) and *Entamoeba* spp. (Amoebozoa), we conclude that breviates and *Entamoeba* spp. likely do not possess vinculin proteins (supplemental figure 1). For *Ectocarpus siliculosus*, its genome has talin, and α-actinin homologs[37], but it does not contain any other integrin-signaling adhesome protein homologs.

## DISCUSSION

Our findings show that the IMAC is not exclusive to obazoans as previously proposed[6,8–11]. We report a nearly full set of IMAC proteins is present throughout Amorphea including four amoebozoan species (*Flabellula citata, Rhizamoeba saxonica, Cryptodifflugia operculatum, Mastigamoeba balamuthi*, and *Nebela* sp.), and two previously unsurveyed obazoan taxa *Lenisia limosa* (Breviatea) and in *Ministeria vibrans* (Opisthokonta). Our increased taxon sampling that spans the breadth of the known diversity within Amoebozoa permitted discovery of these proteins. While the integrin proteins undoubtedly played a role in the origins of animal multicellularity[6,41,42], we do not find any evidence of integrins in the few multicellular lineages of Amoebozoa, those which socially aggregate to form emergent fruiting body structures (*Copromyxa protea* and the model dictyostelids). Interestingly, secondary loss of these proteins seems to be rampant, as the majority of amoebozoan taxa appear to lack integrins or some of the IMAC components, but the caveat is that some of these data are based on incomplete transcriptomic data. Therefore, their absence may be due to the available data. None-the-less, the most parsimonious explanation of the IMAC evolution is that the ancestor of Amoebozoa must have contained them. An analogous scenario of secondary loss of the IMAC plays out in obazoan evolution, in which we see that the closest relative of animals, the Choanoflagellata[39,43,44], as well as the Nucletmycea (also referred to as the Holomycota, the fungal lineage) are devoid of key IMAC components (namely the integrins)[6,11].

The functional domains of ITGA in amoebozoans are similar to the canonical ITGA found in animals and their close relatives like the opisthokont filastereans *Capsaspora owczarzaki*[6], *Ministeria vibrans*, and *Pigoraptor* spp.[7]. Given the homologous nature of the functional domain architecture in amoebozoan ITGA, it is likely that they function at the cellular-level in a similar manner to those in animals. Therefore, we suspect the last common ancestor of Amorphea (LCAA) probably already had ITGA within its genome. Of note, there are potential ITGA homologs in Archaea, therefore it is likely ITGA homologs predate the eukaryotic last common ancestor, but have been retained mostly in Amorphea. We also suspect that the ITGA Ig-like domains possessed by some amoebozoans and obazoans are equivalent to metazoan ITGA legs originally thought to be specific to metazoans[6,11]. An ITGA leg domain was probably in the ITGA of the LCAA, and secondarily lost in some taxa.

Some novel functional domains in Type II representatives are integrin binding domains that are present in ECM protein such as LamG[45], TSP-1[46], and EGF-like domains[47]. Could the last common ancestor of Type II representatives or LCAA contain these integrin binding domains? Other functional domains in amoebozoan ITGA such as protein kinase, TSP-1 together with WSC and intramolecular chaperone are not found in Obazoa. Domain shuffling is an important mechanism for creating a novel genes in evolution[48]. It has been suggested that many domains in the ECM are found in unrelated proteins[49]. Whether these proteins were parts of the domain shuffling in ITGA of the last common ancestor of Amoebozoa, which resulted in ECM evolution remains to be seen and their function remains elusive. However, based on our results, it is possible that we have predicted novel integrin proteins with catalytic as well as adhesive properties.

The domains of amoebozoan ITGB are very similar to the canonical metazoan ITGB and we suspect that the last common ancestor of Amoebozoa retained these canonical ITGB structures including the CRSM motif and three metal binding sites. The cytosolic tail motif “GFFKR” is probably specific to Obazoa and there may be no interaction in the c-terminal intracellular region of integrins in Amoebozoa. The presence of transmembrane and the signal peptide regions are sporadic in Amorphea including metazoan, a previous study showed shedding (evolution from membrane docked to a secretory function) of integrins occurs[27]. The amoebozoan ITGB cysteine pattern of CGXCGGXXXXCXGC possesses more amino acids, including glycine, compared to Obazoa (figure 5F). Glycine is a small non-polar amino acid[50] and due to its small size, glycine positions are often highly conserved within proteins[51,52]; because, glycine residues maintain the overall architecture[53]. It may be interpreted that increased glycine residues could maintain the overall protein structure in amoebozoan ITGB.

The type II ITGB form an independent clade, and it is a sister to the other amoebozoans and *R. ramosa* in figure 4. LamG3 and vWD in ITGB have never been observed in Obazoa and the functions of these type II ITGB proteins are unknown. Some unicellular protists such as breviates, *Capsaspora owczarzaki*, and apusomonads have a long expansion of the CRSM for ITGB [6,11]. We suspect that Breviatea with a long integrin legs form a long integrin heterodimerization complex at the extracellular area of the cell, as both ITGA and ITGB has long legs of nearly equal proportions in *Pygsuia biforma* (figures 2 and 3)[11].

Our prediction is that most of Discosea, which includes *Acanthamoeba* (again with several genomes available) and many other sampled transcriptomes in our study, have lost these proteins. However, one representative discosean (*Mycamoeba gemmipara*) has canonical ITGB homolog (figure 4). Additionally, four other amoebozoan taxa with genome sequences available also appear to have lost ITGB in their evolution. There are amoebozoans in which we have observed only ITGA or ITGB receptors, without their counterpart integrin protein. A study by Li et al. and Schneider et al. showed that homodimerization of ITGA or ITGB might occur by clustering of integrin in a lipid raft[54,55]. Our results show that many of the amoebozoan species singly possess either ITGA or ITGB and yet they possess all the components of signal components of IMAC. These amoebozoan with singular integrins may be capable of initiating the integrin signaling pathway by a similar process.

Also, the question remains that the complexity of extra domains of integrin are not species related and their compositions are a single multidomain protein that is present only in Amoebozoa. Phylogenetically these orthologs (i.e., amoebozoan ITGA and ITGB) form novel clades independent to known organisms and most likely are ancestral to the Amoebozoa as a whole. In order to further examine the overall distributions and protein architectures, genomic sequencing will be necessary, and the combination of localization and gene knockout strategies are necessary to understand the function of these ‘multicellularity’ proteins in unicellular taxa.

## Conclusion

Our most comprehensive comparative genomic and transcriptomic of Amoebozoa has revealed the presence of a full set of IMAC across the Amoebozoa. To date, this complex was believed to be Obazoa specific, our data shows that these IMAC proteins predates Obazoa. Therefore, the last common ancestor of Amorphea contains the necessary machinery of IMAC, yet it remains unknown if and what functional role they play in these unicellular protists. In future work, localization and functionality studies of these IMAC proteins will be investigated.

## Acknowledgements

This project was supported in part by the United States National Science Foundation (NSF) Division of Environmental Biology (DEB) grant 1456054 (http://www.nsf.gov), awarded to MWB. We thank Dr. Andrew J. Roger (Dalhousie University) for advanced access to *Mastigamoeba balamuthi* transcriptome, which was supported by grant MOP-142349 from the Canadian Institutes of Health Research awarded to A.J. Roger.

## Supplemental Materials

### Supplemental Results

Details on the *Sib (similar to integrin beta) proteins found in dictyostelids*. Supplemental Figure 1: Repertoire of IMAC including filamin and tensin in Amoebozoa. Supplemental Figure 2: Repertoire of IMAC in Amoebozoa species. Supplemental Figure 3: Maximum likelihood tree of ITA homolog and other cell adhesion molecules that contain beta-propellers. Supplemental Figure 4: Maximum likelihood tree of ITB homolog and Sib proteins. Supplemental Figure 6: Most significantly enriched motifs in Obazoan ITA, obtained by MEME-suite. Supplemental Figure 7: Most significantly enriched motifs in Obazoan ITB, obtained by MEME-suite. Supplemental Figure 8: Most significantly enriched motifs in Sib protein, obtained by MEME.

## METHODS

### Method Details

We have strategically sampled 118 Amorphea species and their transcriptomes and genomes to cover all known Amorphea subgroups. Additionally, we also searched for IMAC proteins in any of the eukaryotic data in NCBI.

### Culturing and Isolation of Mycamoeba gemmipara

*Mycamoeba gemmipara* was obtained from Q. Blandenier and cultured as in Blandenier et al. 2017[90].

### Ultra-low input/ Single-Cell cDNA library construction based on Smart-Seq2

The cultured amoeba from *Mycamoeba gemmipara* were subjected to Smart-Seq2 [91] for cDNA library construction as in Onsbring *et al*. 2019[92]. About 50 cells scraped off of a agar plate using a 30-gauge platinum wire placed directly into a 200 µL thin-walled PCR tube containing the cell lysis mixture [91]. Each reaction was then subjected to six freeze-thaw cycles in -80 °C isopropanol and ∼25 °C DI H_2_O respectively, before following the remainder of the protocol. A modified version of Smart-Seq2 [91] was used for RNA isolation and dscDNA preparation. cDNA libraries were prepared using a Nextera XT DNA Library Prep Kit (Illumina, CA) following the manufacturer’s protocol with dual index primers. The *Mycamoeba gemmipara* library was sequenced using an Illumina HiSeq 4000 at Genome Quebec.

### Transcriptomic Sequencing Assembly

Raw sequence data from the Illumina sequencing platform or from Bioproject accession number PRJNA380424 [14] were subjected to Trimmomatic v 0.35 [78], for cleaning and trimming of poorly called reads or specific to the adaptor will be used in sequencing with the paired-end data. Quality filtered and cleaned high quality reads were assembled through Trinity [73] for *de novo* assembly. Nucleotide sequences were translated with Transdecoder v 5.5.0 (https://github.com/TransDecoder/TransDecoder/) which also predicts open reading frames, and Pfam domains [88] as well. Contigs of transcriptomes were blasted against the non-redundant database of NCBI using the BlastX program of Blast+ suite [93]. Each predicted ORFs were assigned an ortholog number through the OrthoMCL v5 [77] database. For taxa with clearly truncated predicted proteins, the computationally laborious transcriptome assembly pipeline, Oyster River Protocol[84], was performed in an attempt to retrieve predicted a full-length integrin proteins.

### Integrin Protein analysis

We selected a model organism IMAC proteins from NCBI: ITGA5:P08648, ITGB1:NP_002202, Talin:AAF27330, Parvin:AAH16713, PINCH:NP_060450.2, vinculin:AAH39174, FAK:AAA35819 Paxillin:AAC50104 ILK:NP_001014794, α-actinin:AAC17470, Filamin:AAF72339 and Tensin:AAG33700. InterProScan 5.27-66.0[23] was used to determine these IMAC protein domain architecture along with SignalIP v 5.0 [83] and TmHmm v 2.0 [94]. Meme-suite 5.0.4 [85]was used to examine integrin motifs. DeepLoc v 1.0[86] was used to determine the subcellular localization of integrin proteins. We set the minimum criteria of canonical IMAC proteins based on their architecture and motifs. We used OrthoMCL to assign IMAC proteins their own ortholog numbers.

Since OrthoMCL database is heavily metazoan biased, we created an ortholog database with eukaryotes listed in resources table. A modified version of OrthoMCL-DB [95] was used to create a novel ortholog database using the above listed transcriptomes and genomes as well as the whole OrthoMCL DB v5. To do this, an all-against-all Blast using Diamond-BlastP[76] was conducted using each protein from the above data as queries. The Diamond-BlastP collected up to 1,000 hits with e-value of 1e-5 as a cutoff for putative homology. Blast results were clustered using the OrthoMCL pipeline methodology [95].

BlastP was used to search for novel amoebozoan IMAC proteins through the new ortholog database and using the previous amoebozoan ortholog protein as a query. Using a modified custom pipeline was used to collect a novel IMAC orthologs. These methods generated various orthologs with a similar protein architecture. We created a FASTA file of all novel eukaryote integrin proteins based on their protein architecture. Clustering of these integrin proteins was performed by CD-Hit [71], which used the condition of 0.95 global sequence identity. From CD-Hit output, we used Diamond [76] to perform All-vs-ALL blast with database size of 50 million sequences. Consequently, a custom script of Markov cluster (MCL) algorithm[89] was used to cluster the output of Diamond. A custom Python script was used to eliminate any MCL clustered integrin proteins that were less than 500 AA. We used a custom Python pipeline to collects ortholog sequences that clustered to model metazoan IMAC orthologs listed above.

Sometimes, we fail to capture integrin proteins even after all these strenuous processes. Therefore, Pfam ID of Integrins was used to grab any putative integrins from Transdecoder output. For ITGA, we used FG-GAP (PF01839), and for ITGB we used PSI_integrin (PF17205), integrin_beta (PF00362), integrin_b_cyt (PF08725) and integrin B tail (PF07965). We disregarded any protein sequences with PFam ID lower than e-value of 1e-10. The identity of the putative integrins were confirmed by InterProScan, SignalIP, TmHmm, DeepLoc, and Meme-suite.

For each putative integrin protein, we examined proteins motifs with Meme-Suite, and always blasted our proteins of interest using BlastP against NCBI’s GenBank NR database to check for possible contaminations. For integrin proteins with a novel domains, the read depth of these transcripts were examined with Rsem. The final amoebozoan integrin architectures were compared with metazoan integrin proteins. A custom Python script was used to create a presence/absence binary (1,0) matrix of IMAC proteins across Amoebozoa. A custom R script were used to create a heatmap.

In our current study these data are transcriptomic in nature and by this virtue some of the predicted transcript may be truncated or missing. However, identifying full-length transcripts from short reads from Illumina technology can be difficult because assembled transcript can be fragmented or incomplete due to alternative splice sites and untranslated regions [96]. Therefore, it is possible that some transcripts are not fully assembled, and therefore their corresponding predicted protein is truncated, thus the transmembrane region and signal peptide region may be missing. The same can be said for the absence of these proteins within our data, because transcriptomes contain only genes being expressed at the time mRNA was harvested some genes within an organism’s genome may be missed. None-the-less, our methodology permitted the unparalleled discovery of these proteins within a whole supergroup, highlighting the utility of the method.

### Manually assembled contigs in Sequencher

Where necessary due to fragmentation, contigs of our automated de novo transcriptomic assemblies as above from each gene of interest were blasted (BlastN) back to the raw nucleotide data and to the assembly for each transcriptome. Hits were collected and assembled using Sequencher v 5.4.6 (GeneCodes, Madison, WI, USA). The taxa which we had to take this approach were *Amphizonella* sp. 2, *Centropyxis aerophyla, Goncevia foncevia* and *Hyalosphenia papilio, Pellita catalonica* for ITGA, and *Nebela* sp. for both integrin proteins.

### *Presence of integrin in the* Mastigamoeba *genome*

To confirm the presence of these genes in the genome, we blasted the integrin transcripts of *M. balamuthi* to the whole shotgun genome data (CBKX00000000; https://www.ncbi.nlm.nih.gov/nuccore/CBKX000000000.1) on NCBI [33]. The common splicing sites were for searched in integrins intron and exon boundaries by assembling the integrin genomic contig and integrin transcript in Sequencher v 5.4.6. The genome contigs of which contained integrin gene were searched for additional ORFs to examine the phylogenetic affinity of other ORFs on the genomic contigs. We inferred maximum likelihood phylogenetic tree of tenascin (adjacent to the ITGB gene on CBKX010020318.1) and PA14 (adjacent to the ITGB gene on CBKX010020319.1), we used same conditions as the integrin phylogenetic trees (see below).

### Phylogenetic trees

To look for evolutionary relationships of ITGA and ITGB within Amorphea, protein sequences were aligned by Mafft-Linsi with the parameters “--maxiterate 1000” and “--local pair”. Ambiguous sites were trimmed from the alignments using Bmge [80] by gap penalty of 0.8 settings. Maximum likelihood (ML) trees were inferred from these trimmed alignments in IQtree v 1.5.5 [81]under the LG model with the C60 series model of site heterogeneity. Each tree is ML bootstrapped (MLBS) by 1,000 pseudoreplicates. From the resultant tree we used a custom Python script implementing ETE-Toolkit (www.etetoolkit.org) to map protein domain architecture (IPRSCAN domain IDs) onto the tree.

### Data and software availability

The accession number for the *Mycamoeba gemmipara* and *Mastigamoeba balamuthi* transcriptome raw reads is NCBI-SRA: XXXXX and XXXXX also shown in Key resources table. All protein and nucleotide sequences of each gene, all protein alignments (trimmed and untrimmed) for phylogenetic trees, and phylogenetic trees are available on the Dryad repository (https://datadryad.org/ | Accession: XXXXX).

## Supplemental Text

### SUPPLEMENTAL METHODS, NOTES, AND RESULTS

#### Key resources Table

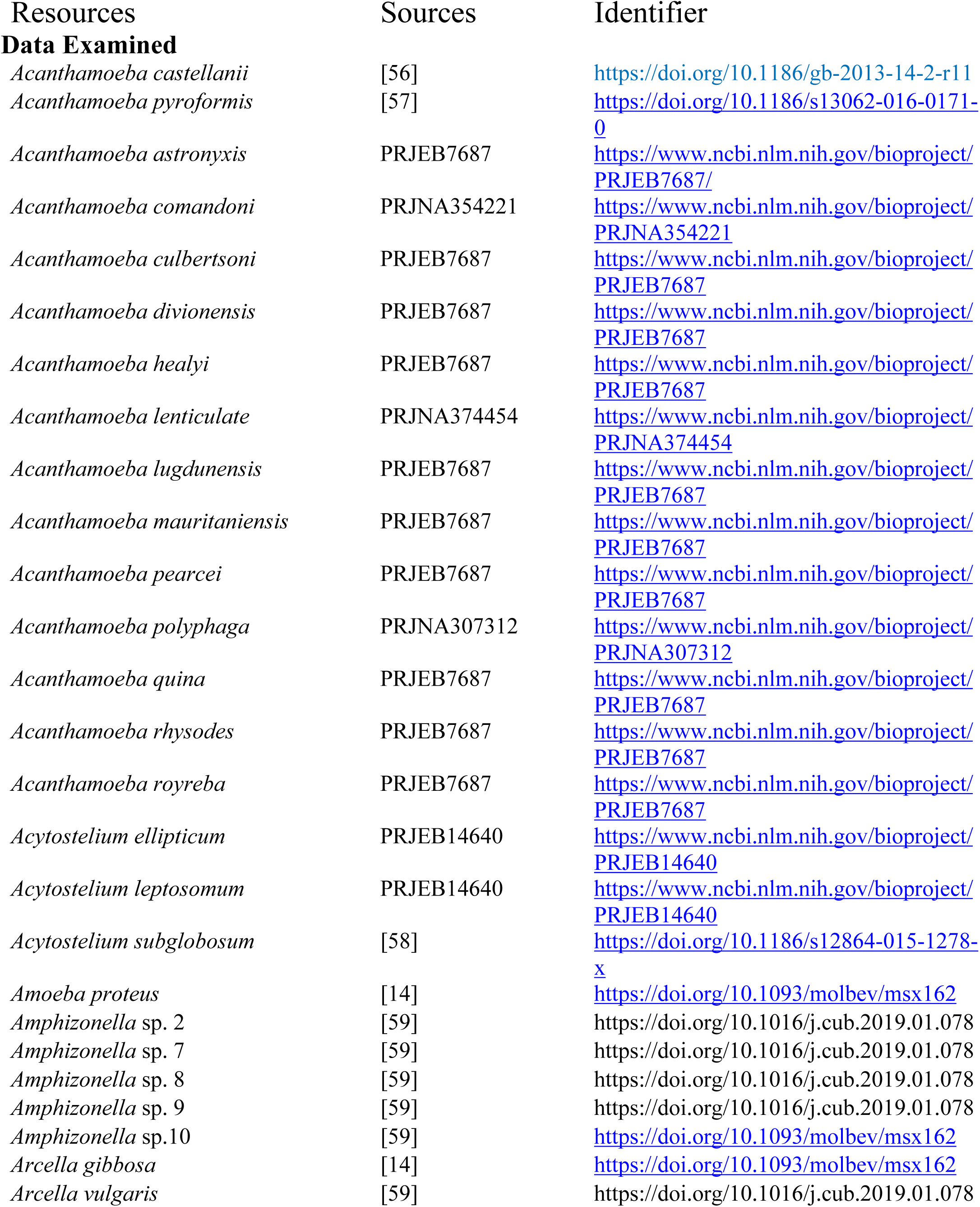
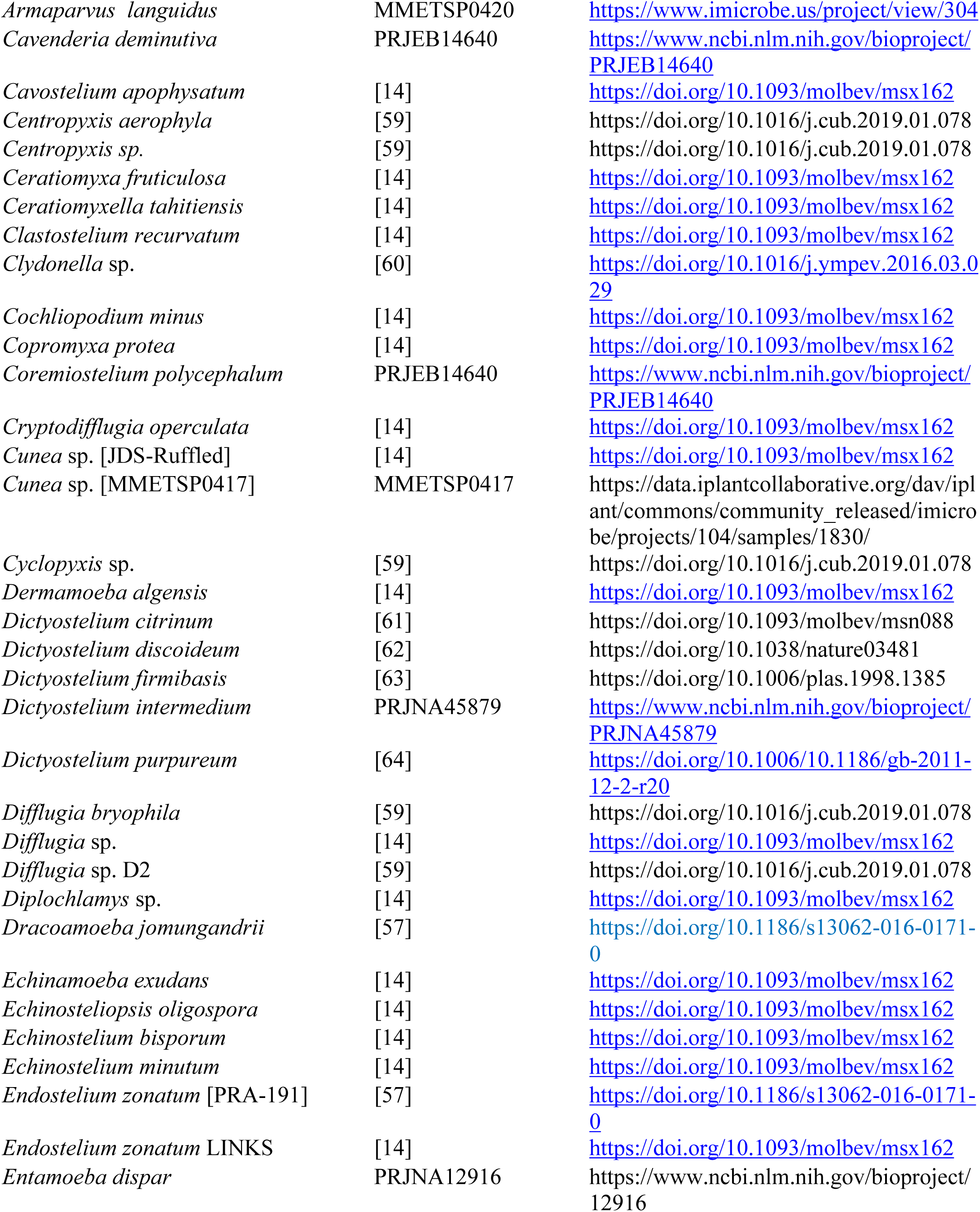
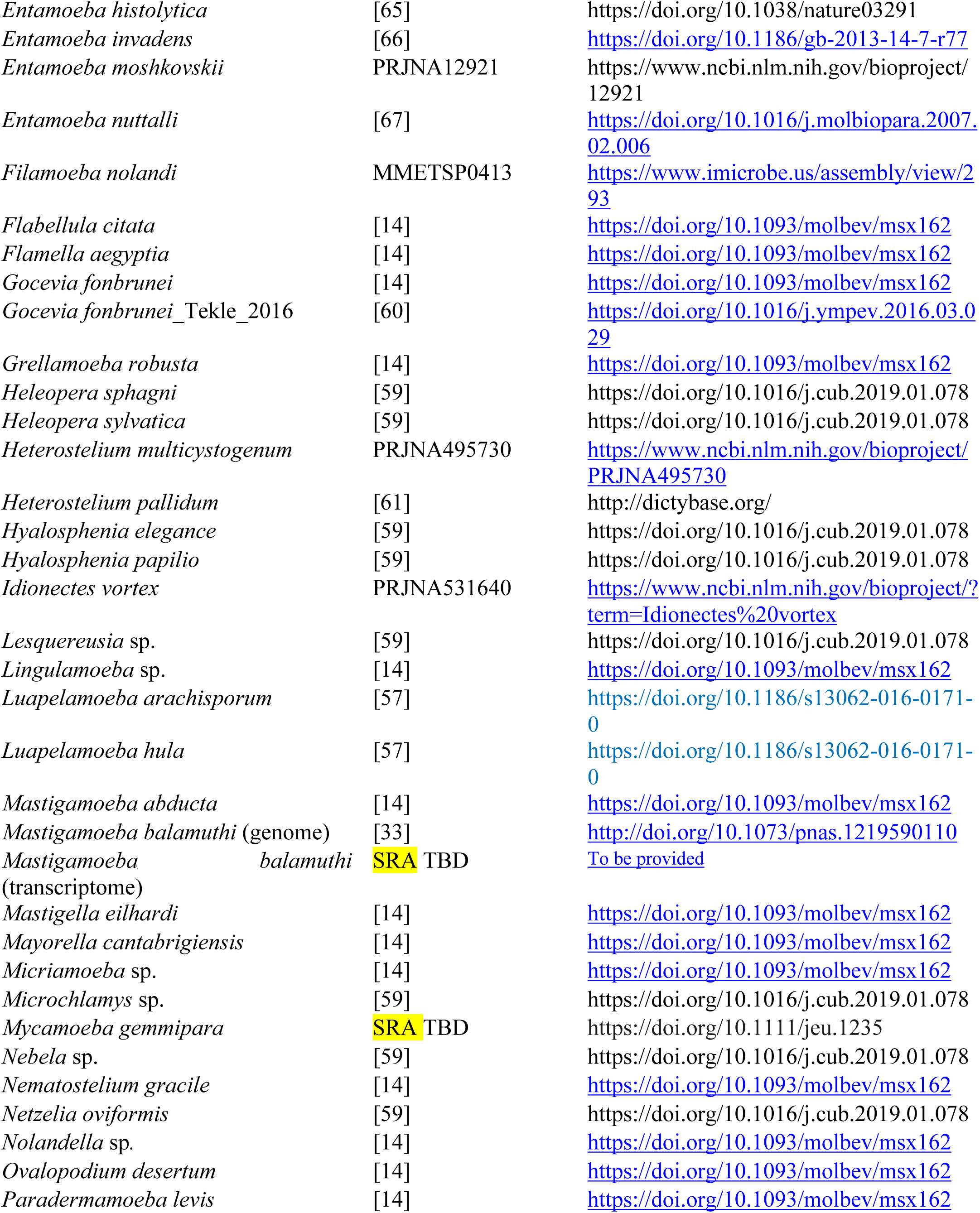
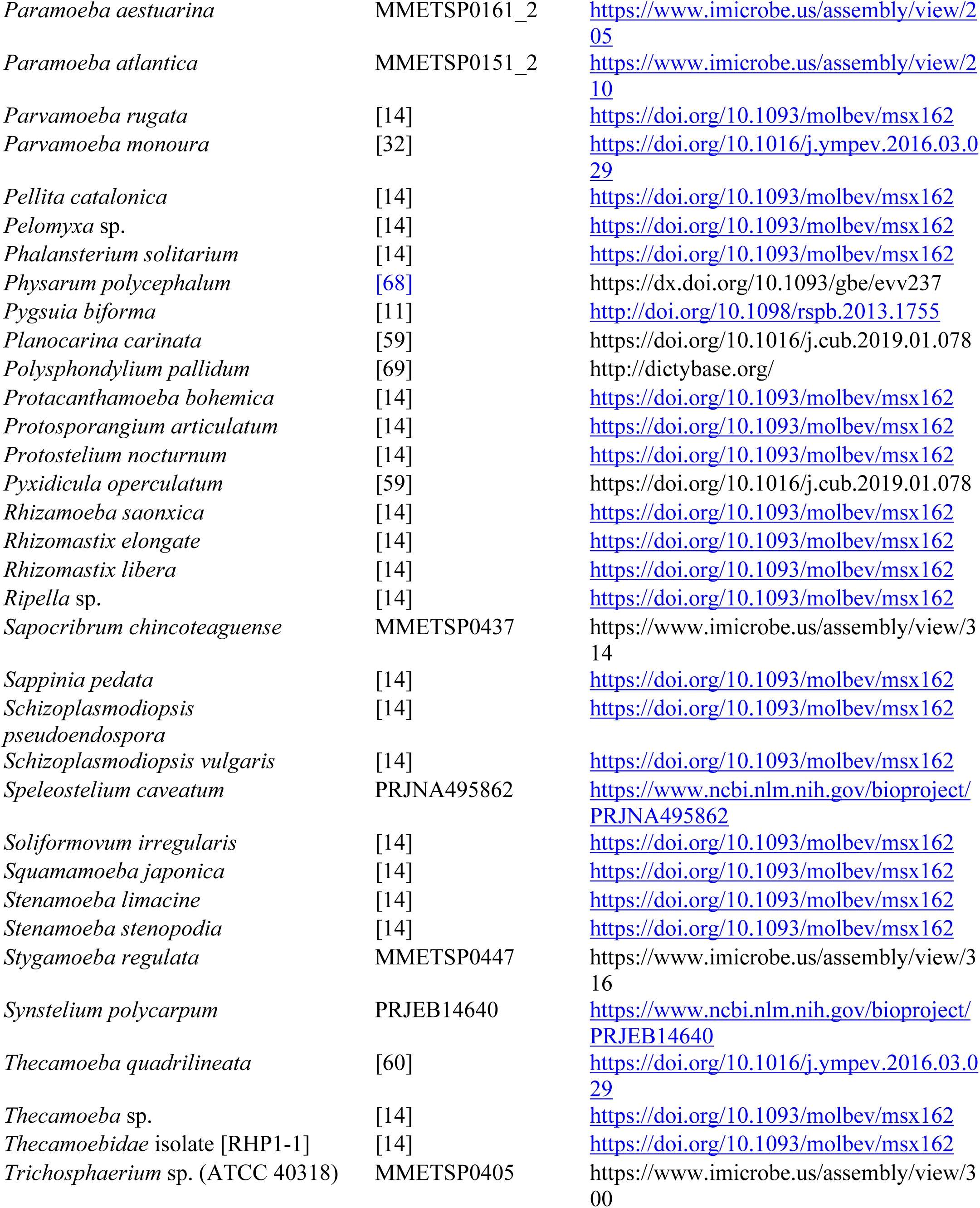
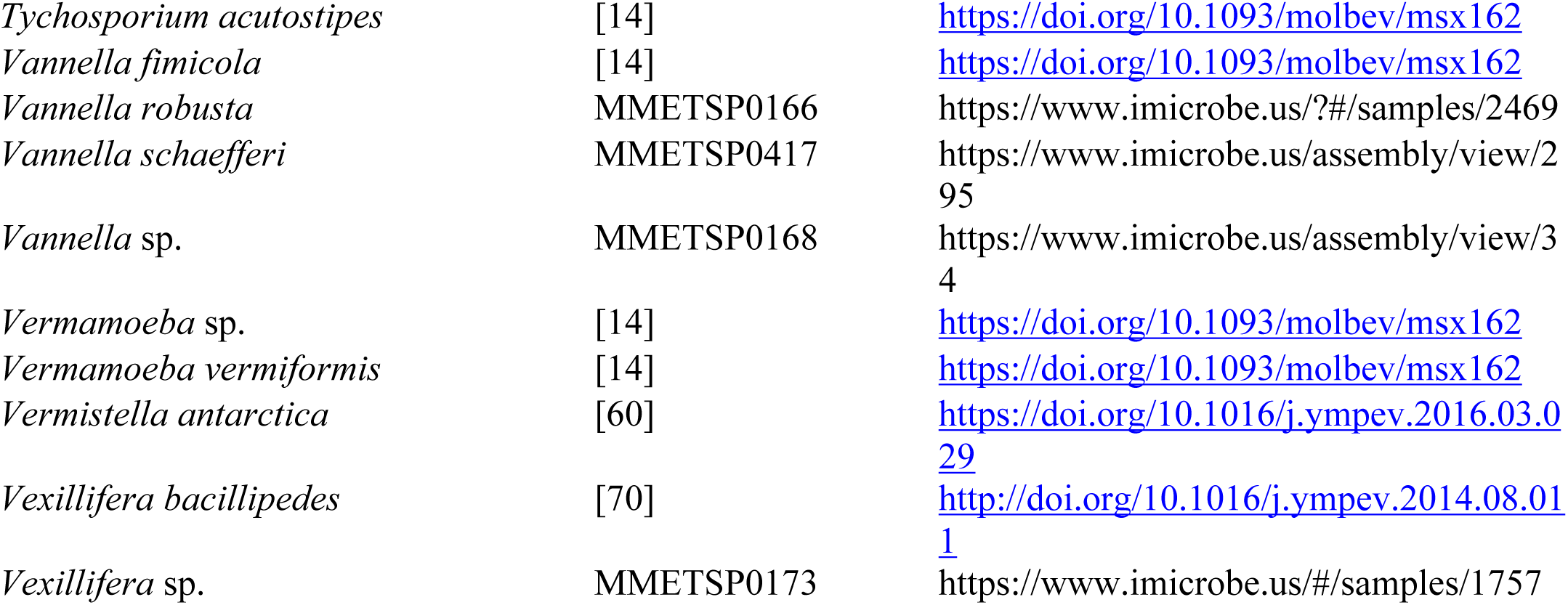

#### List of Stramenophile TAXA from MMETSP

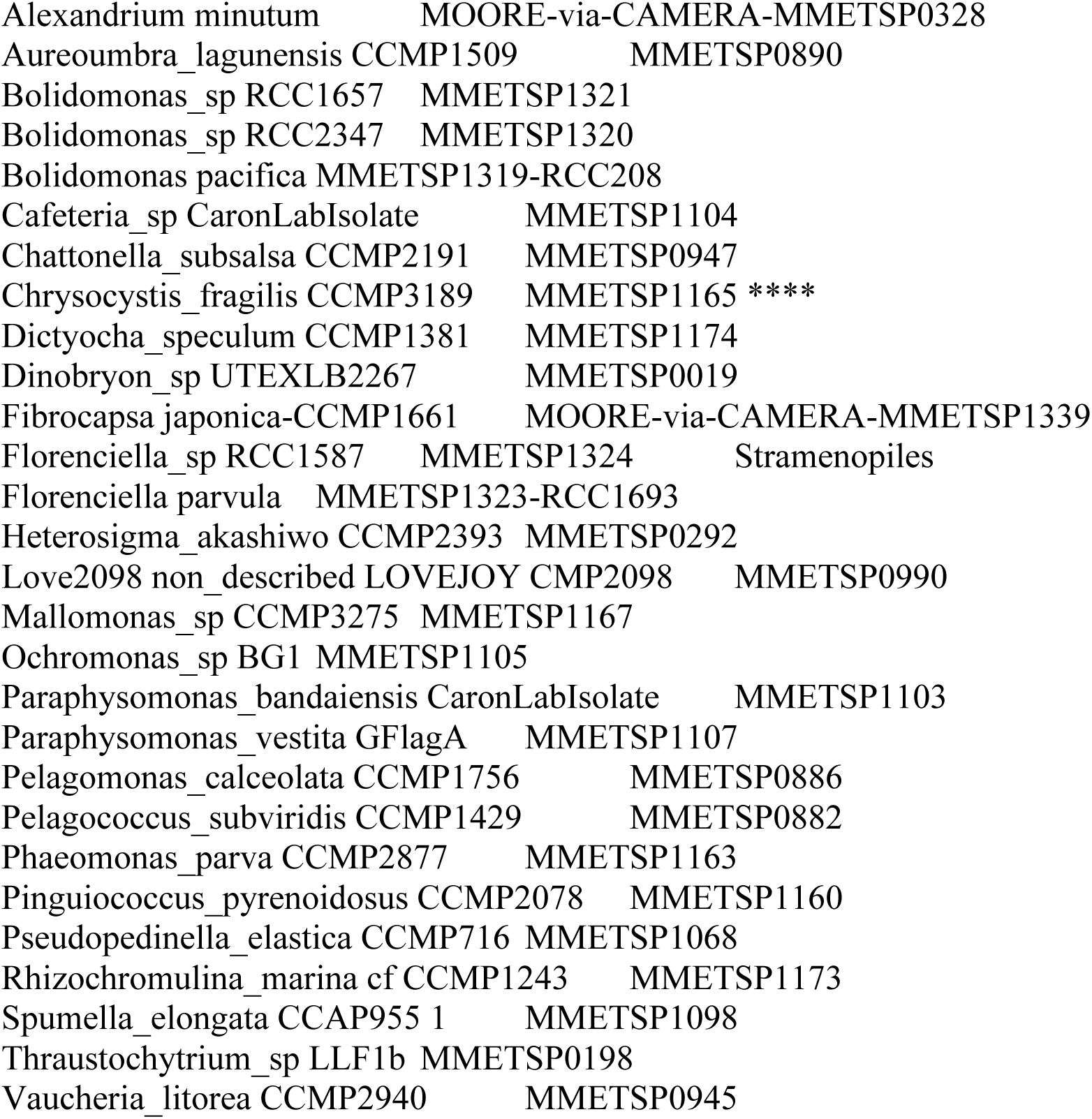

#### Reagents

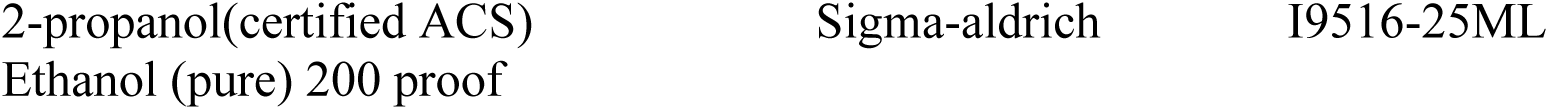

|  |  |  |
| --- | --- | --- |
| 2-propanol(certified ACS) | Sigma-aldrich | I9516-25ML |
| Ethanol (pure) 200 proof |  |  |

#### Critical Commercial Assay Kits

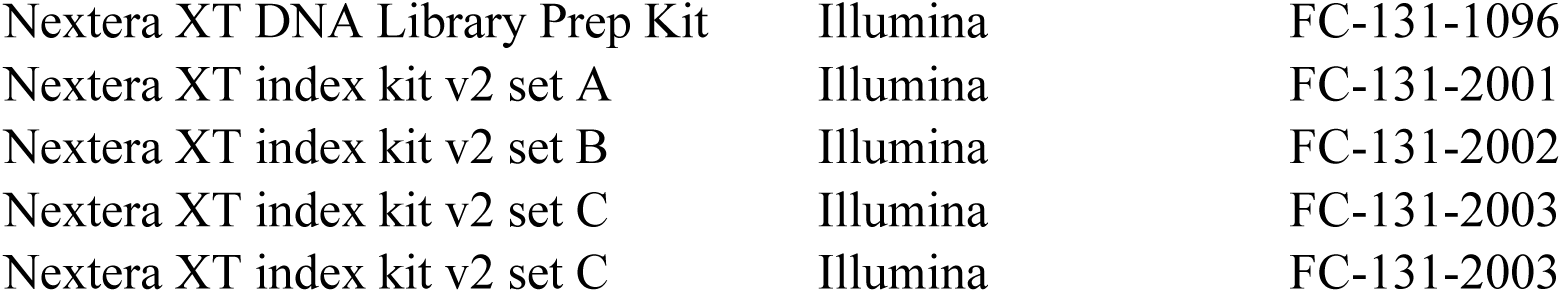

#### Software and Algorithms

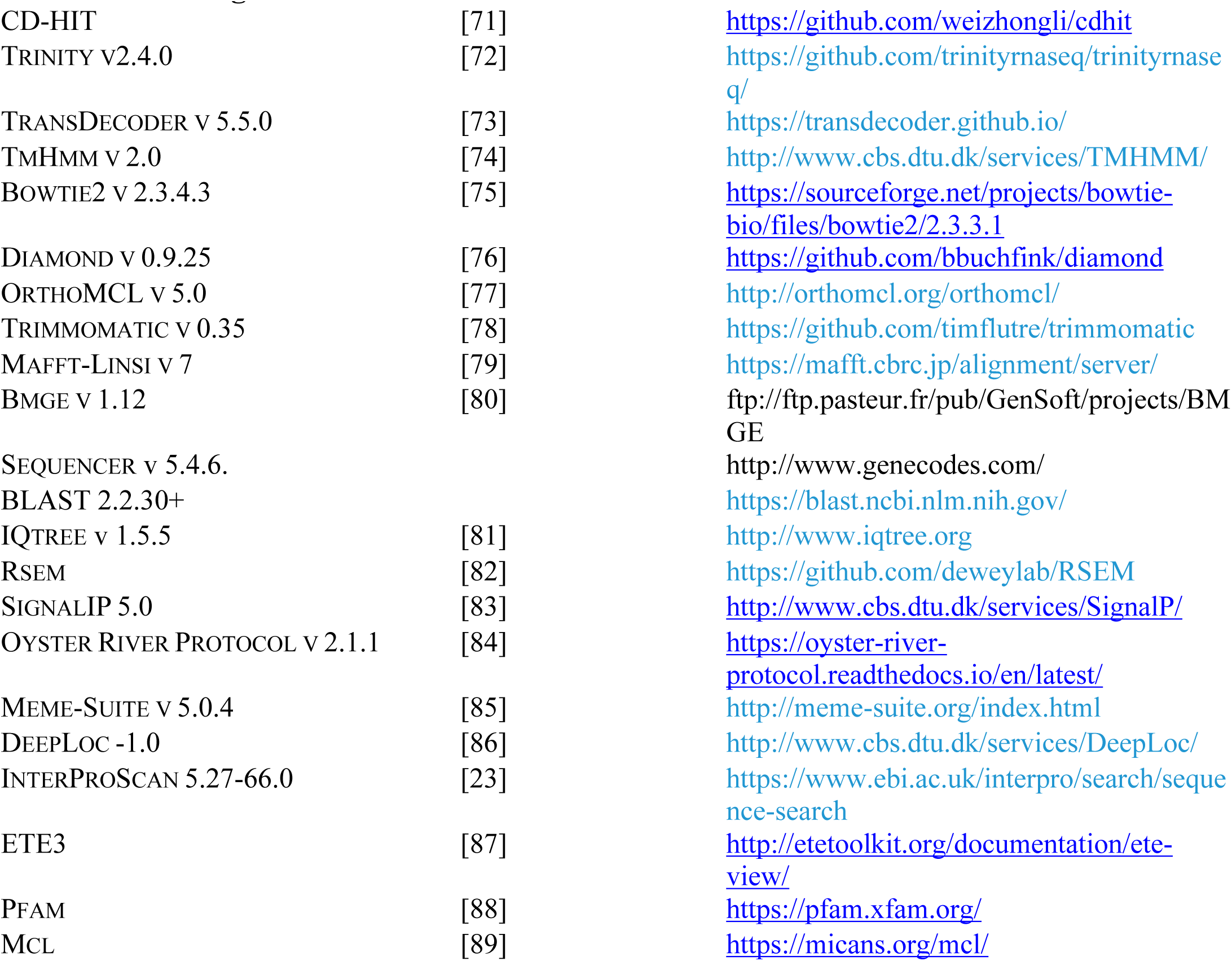

#### Contact for Reagent and resources sharing

Further information and request for resources should be directed to, and will be fulfilled by, the corresponding author Matthew W. Brown.

### *Sib (similar to integrin beta)* proteins found in dictyostelids | Sib Proteins comparison with ITB-like

SibA, a cell adhesion molecule that binds to phagocytic particles similar to ITB, was discovered in *Dictyostelium discoideum* [45]. Sib proteins (A to E) are capable of binding to talin and SibC regulates cell adhesion [46]. However their protein structure is different from the canonical ITB in a lack of three cation binding motifs as well as a cysteine rich region at the C-terminus [26]. Although the varioseans, described above, are phylogenetically close to dictyostelids their type two ITB-like proteins with three cation motifs are distinct from Sib proteins, though with a great variation in amino acid sequences. Interestingly, we did not observe Sib proteins outside of the Dictyostelia clade, suggesting that they evolved recently in the dictyostelids. The protein architecture of SibA has a similar composition to metazoan ITB solely with a GXXXG motif in the transmembrane region, but it is observed in type II ITB (see supplemental table 2). In addition, LamG3, vWD domains in type II ITB have never been observed in Sib.

---

**Supplemental Figure 1:**
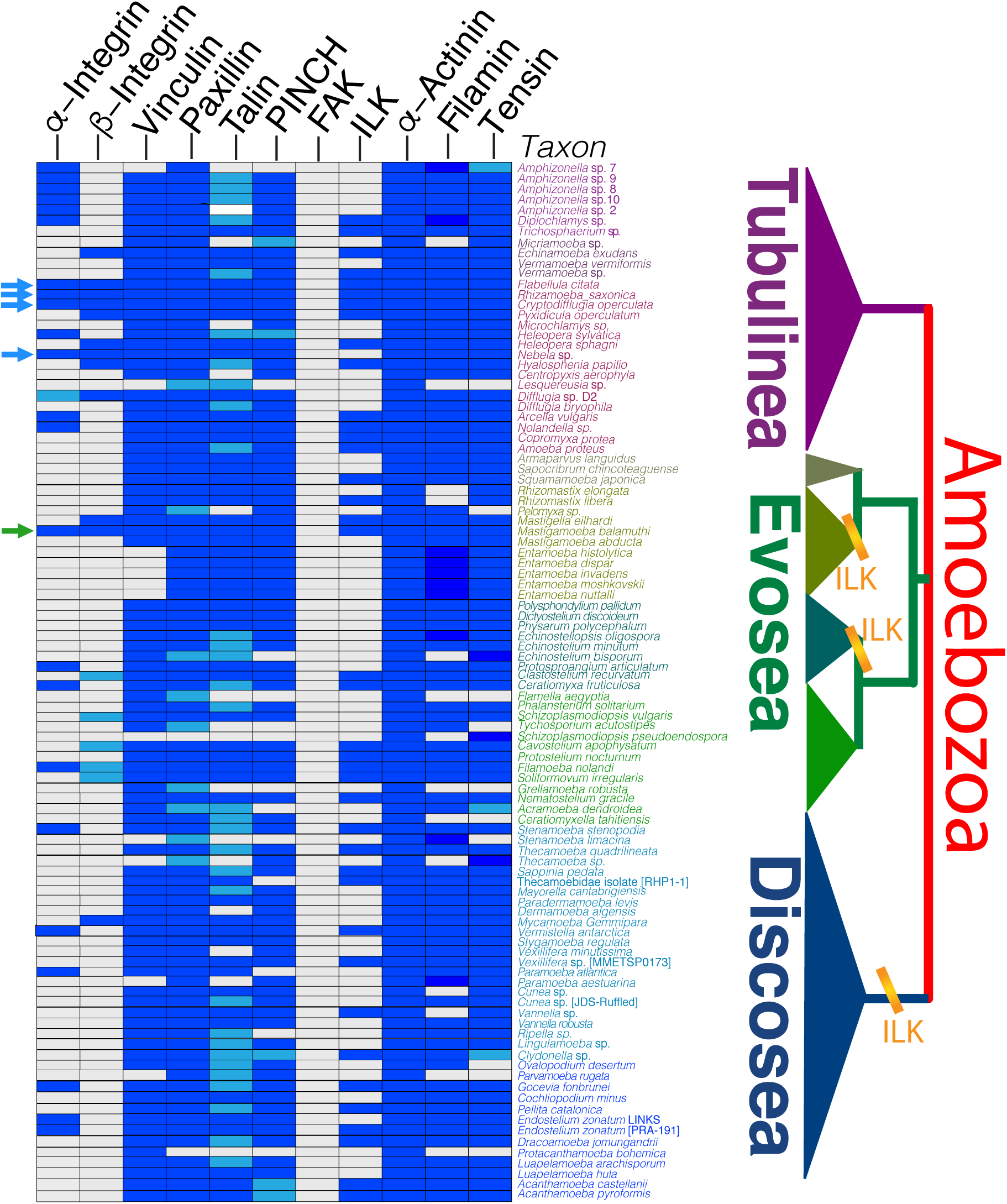
Repertoire of IMAC including filamin and tensin in Amoebozoa: names of IMAC proteins are listed at the top and names of amoebozoan taxa are listed on side of the map. Arrows indicates amoebozoan species that have nearly complete set of IMAC proteins. Dark blue indicates the presence of a canonical IMAC protein, white indicates the absence of an IMAC protein and light blue indicates a truncated form of IMAC protein.

**Supplemental Figure 2:**
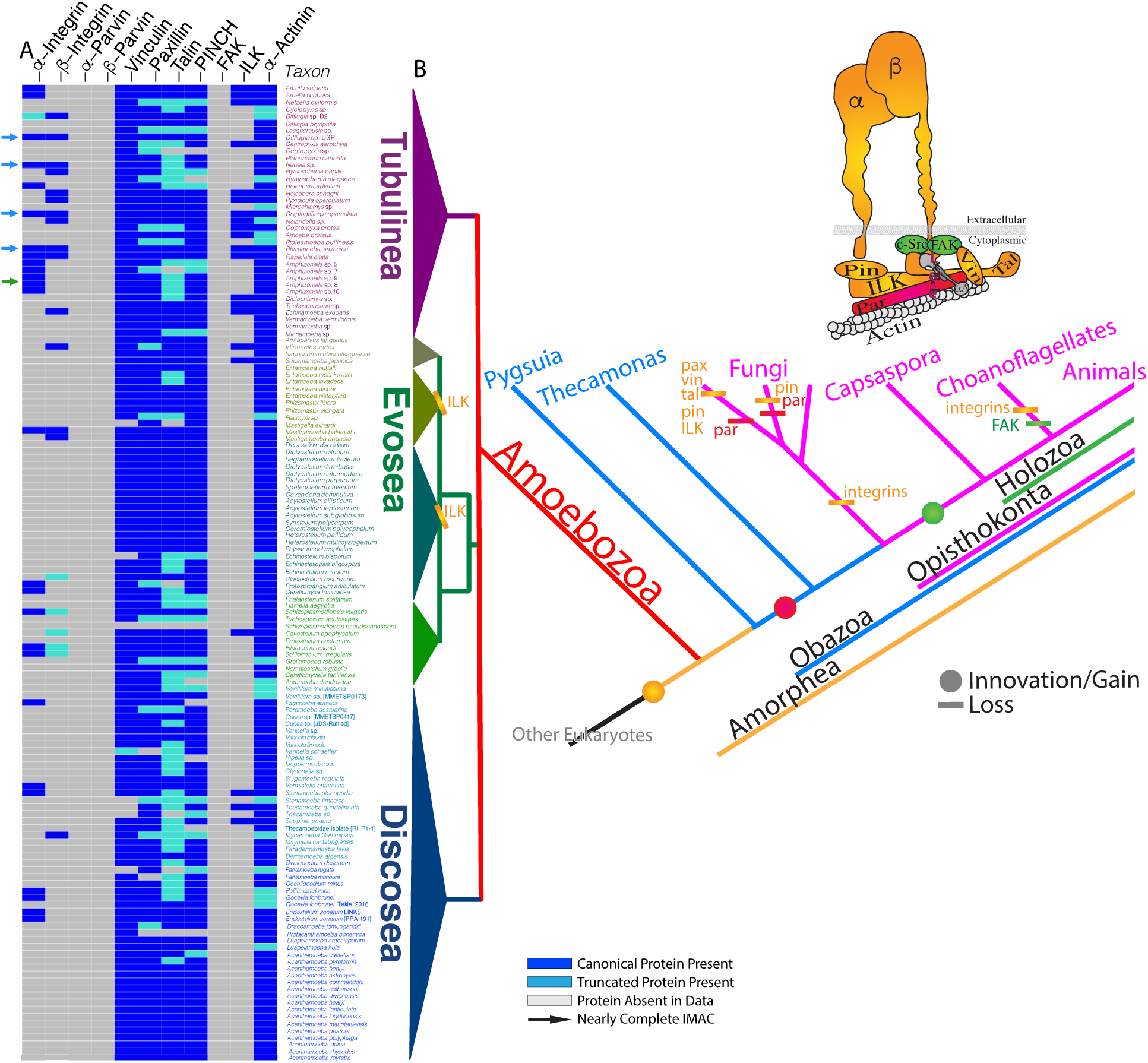
Repertoire of IMAC in only amoebozoan species that have either integrin alpha or beta. Names of IMAC proteins are listed at the top and names of amoebozoan taxa are listed on side of the map. Arrows indicates amoebozoan species that have nearly complete set of IMAC proteins. Dark blue indicates the presence of a canonical IMAC protein, white indicates the absence of an IMAC protein and light blue indicates a truncated form of IMAC protein

**Supplemental Figure 3:**
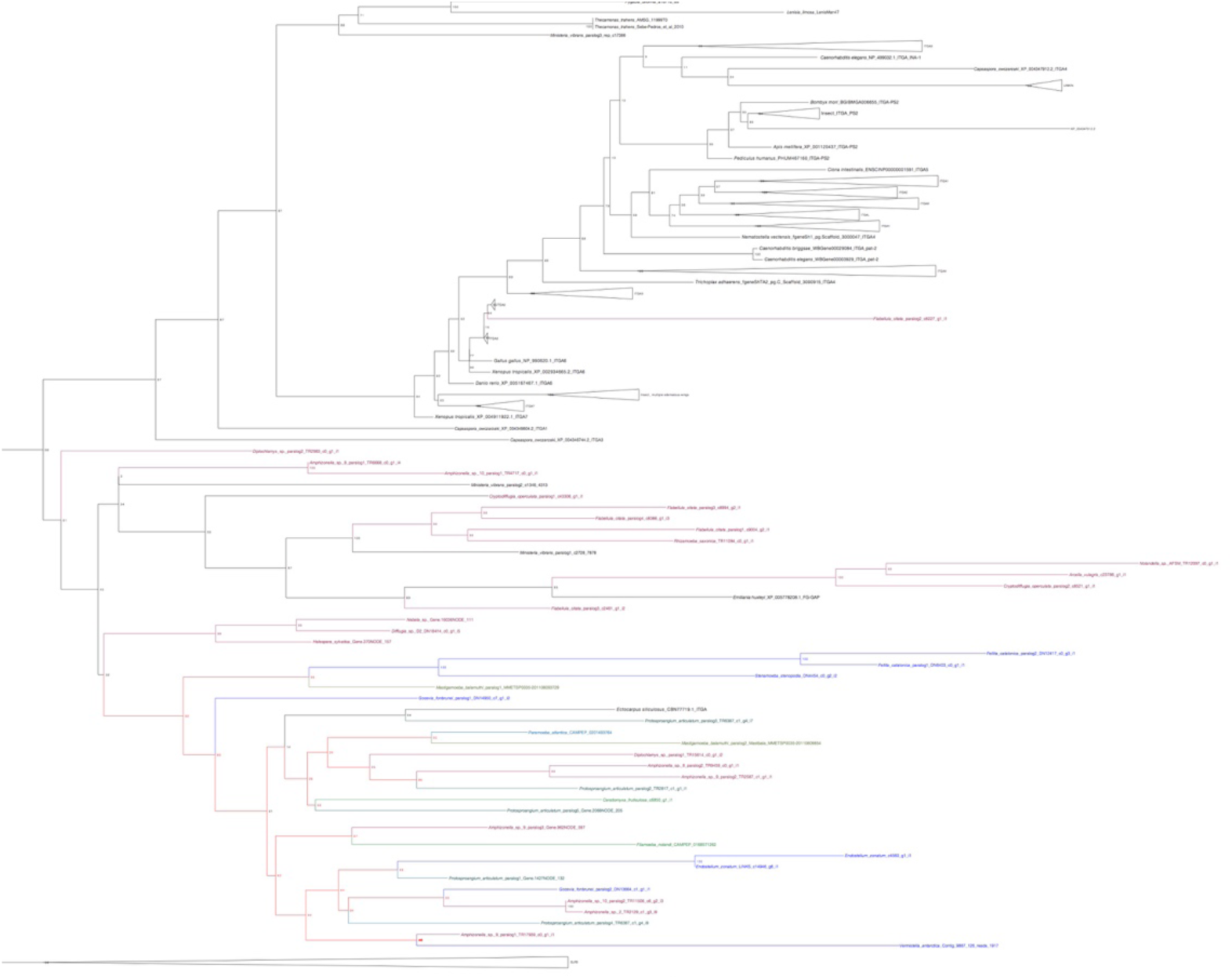
Maximum likelihood tree of ITA homolog and other cell adhesion molecules that contain beta-propellers. Bootstrap values ≥50 are shown on the branch points. Phylogenetic tree was built with IQtree v1.5.5 under under the LG+C60+F+G model of protein evolution ML bootstrap (MLBS) (1000 ultrafast BS reps) values respectively. The protein sequences were aligned by Mafft with the parameter linsi, maxiterate 1000 and local pair. BMGE was used to mask the alignment with the parameter of gap penalty of 0.8. ITA homolog phylogenetic tree are rooted with Linkin protein. Amoebozoan ITA genes are mostly concentrated on a single clade (colored) except for the *Flabellula citata* paralog.

**Supplemental Figure 4:**
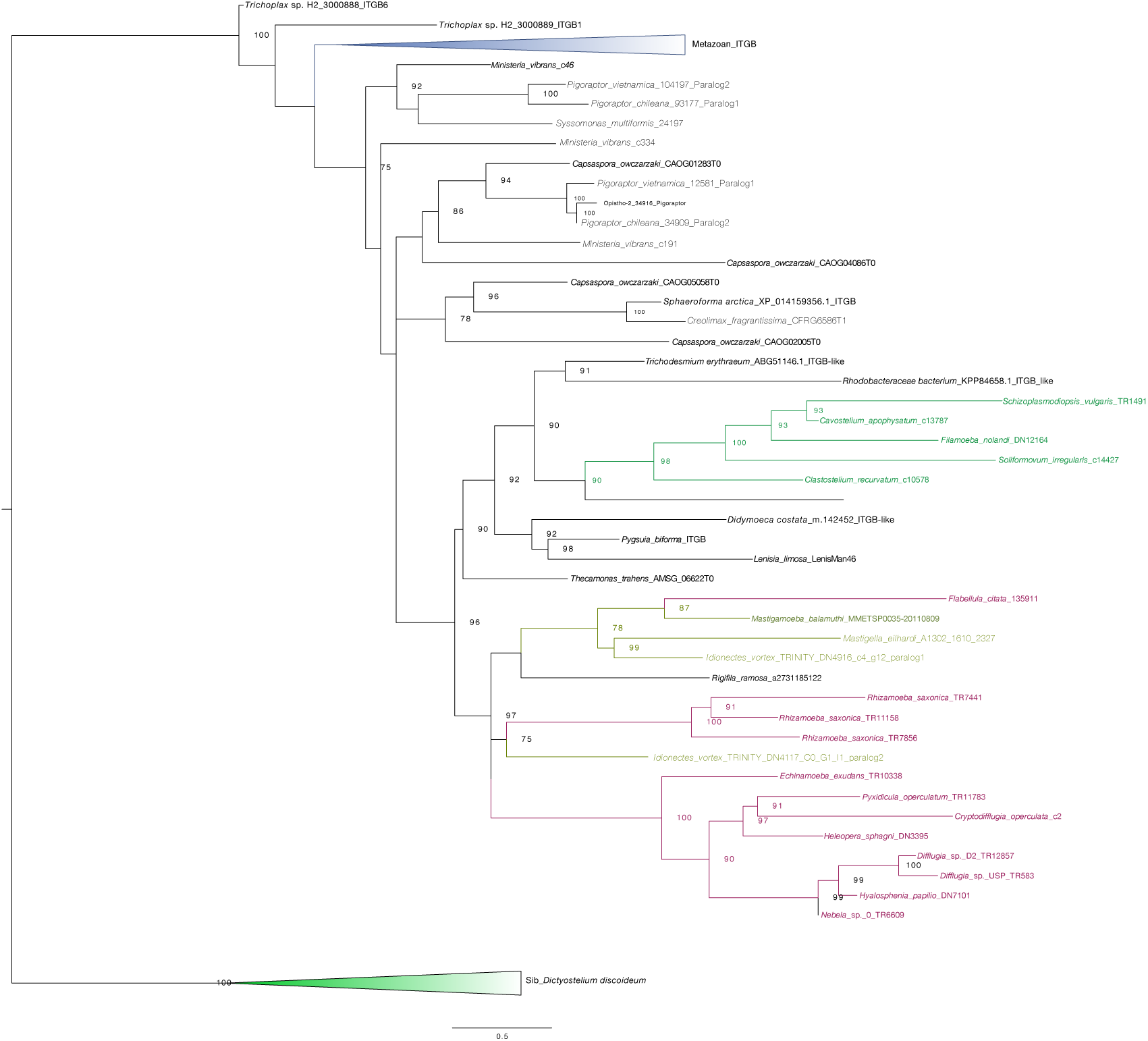
Maximum likelihood tree of ITB homolog and Sib protein. Bootstrap values ≥50 are shown on the branch points. Phylogenetic tree was built with IQtree v1.5.0 under the LG+C60+F+G model of protein evolution ML bootstrap (MLBS) (1000 ultrafast BS reps) values respectively. The protein sequences were aligned by Mafft with the parameter linsi, maxiterate 1000 and local pair. BMGE was used to mask the alignment with the parameter of gap penalty of 0.6. ITB homolog phylogenetic tree are rooted with Sib protein. Amoebozoan ITB genes are mostly concentrated on a single clade (colored).

**Supplemental Figure 6:**
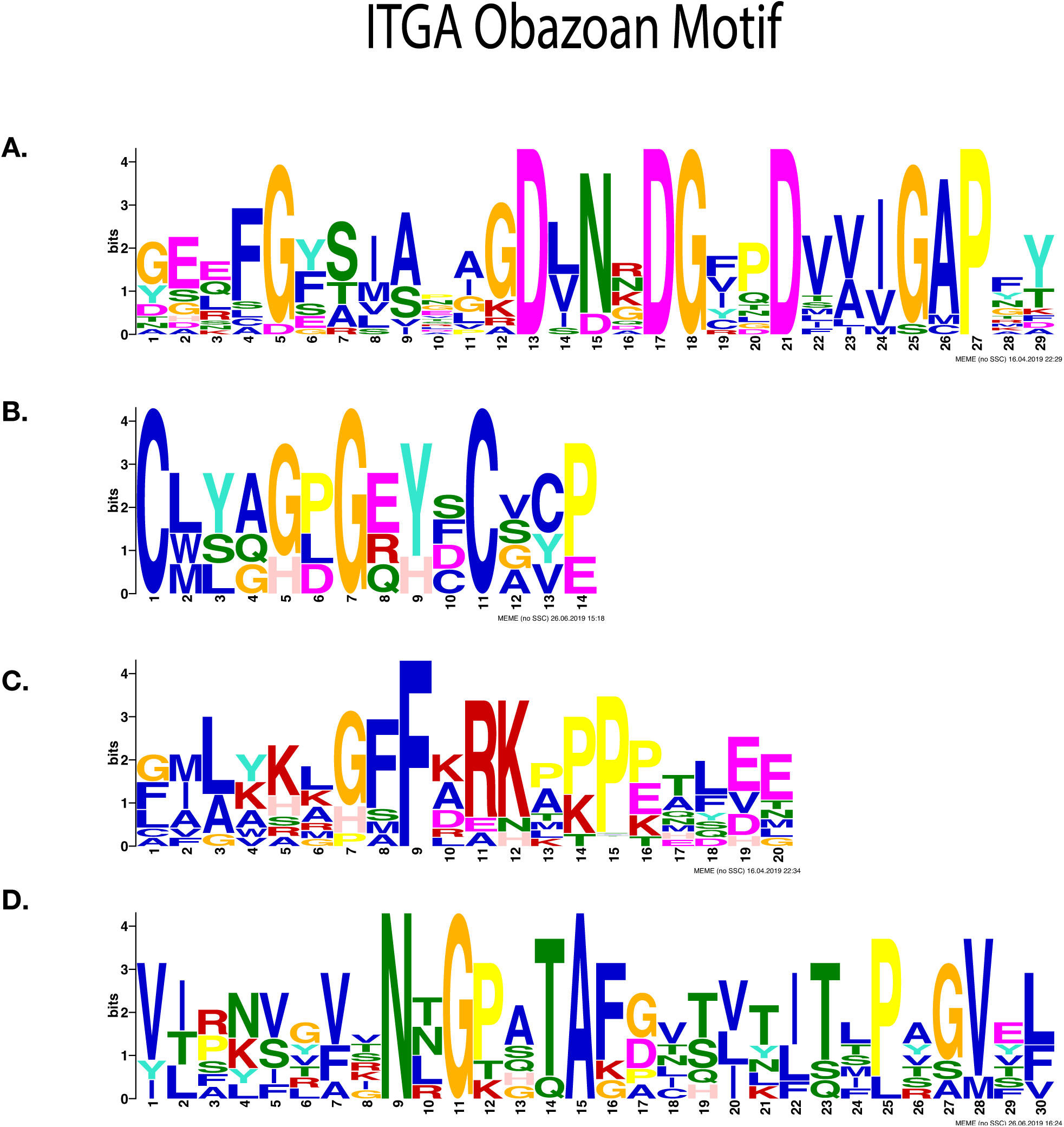
Most significantly enriched motifs in Obazoan ITA, obtained by MEME-suite. (**A**) Consensus cation motif and FG-GAP obtained by RNA-seq for an Obazoan ITA, (**B**) EGF motif found exclusively in *Ministeria vibrans* (**C**) GFFKR motif found in Obazoa, (**D**) immunoglobulin like motif found in Obazoa.

**Supplemental Figure 7:**
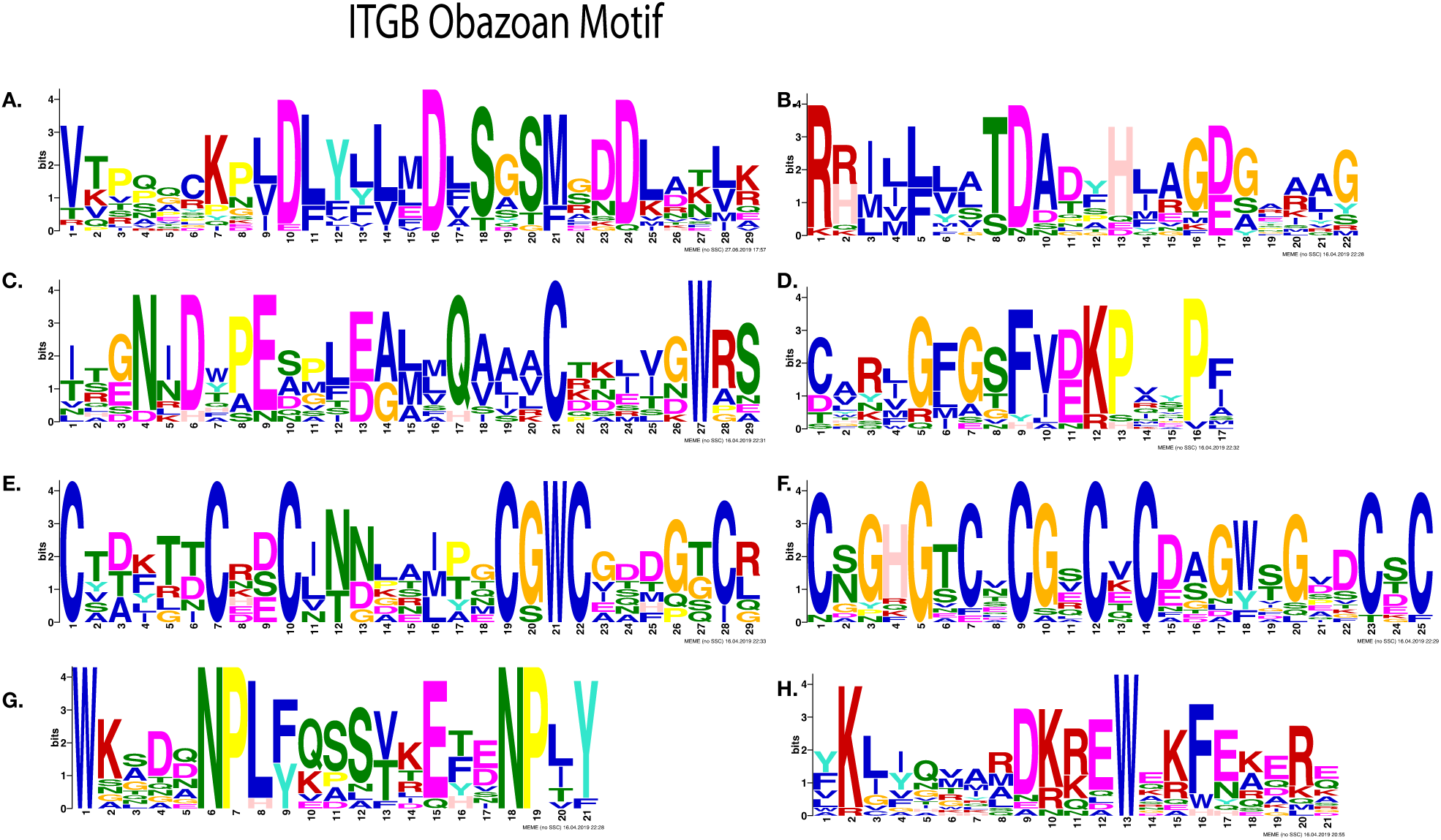
Most significantly enriched motifs in Obazoan ITB, obtained by MEME. (**A**-**D**) Cation binding motifs (**A**) MIDAS binding motif DXSXS (**B**) Possible fifth position of MIDAS and ADMIDAS, (**C**) ITB SyMBS (green) and MIDAS (Yellow), (**D**) Possible E of SyMBS (first amino acid of SyMBS) (**E**-**F**) Cysteine rich motif, (**E**) PSI domain, (**F**) Cysteine-rich motif stalk, (**G**-**H**) C-terminal motifs (**G**) NPXY motif, (**H**) Possible KLXXXD motif

**Supplemental Figure 8:**
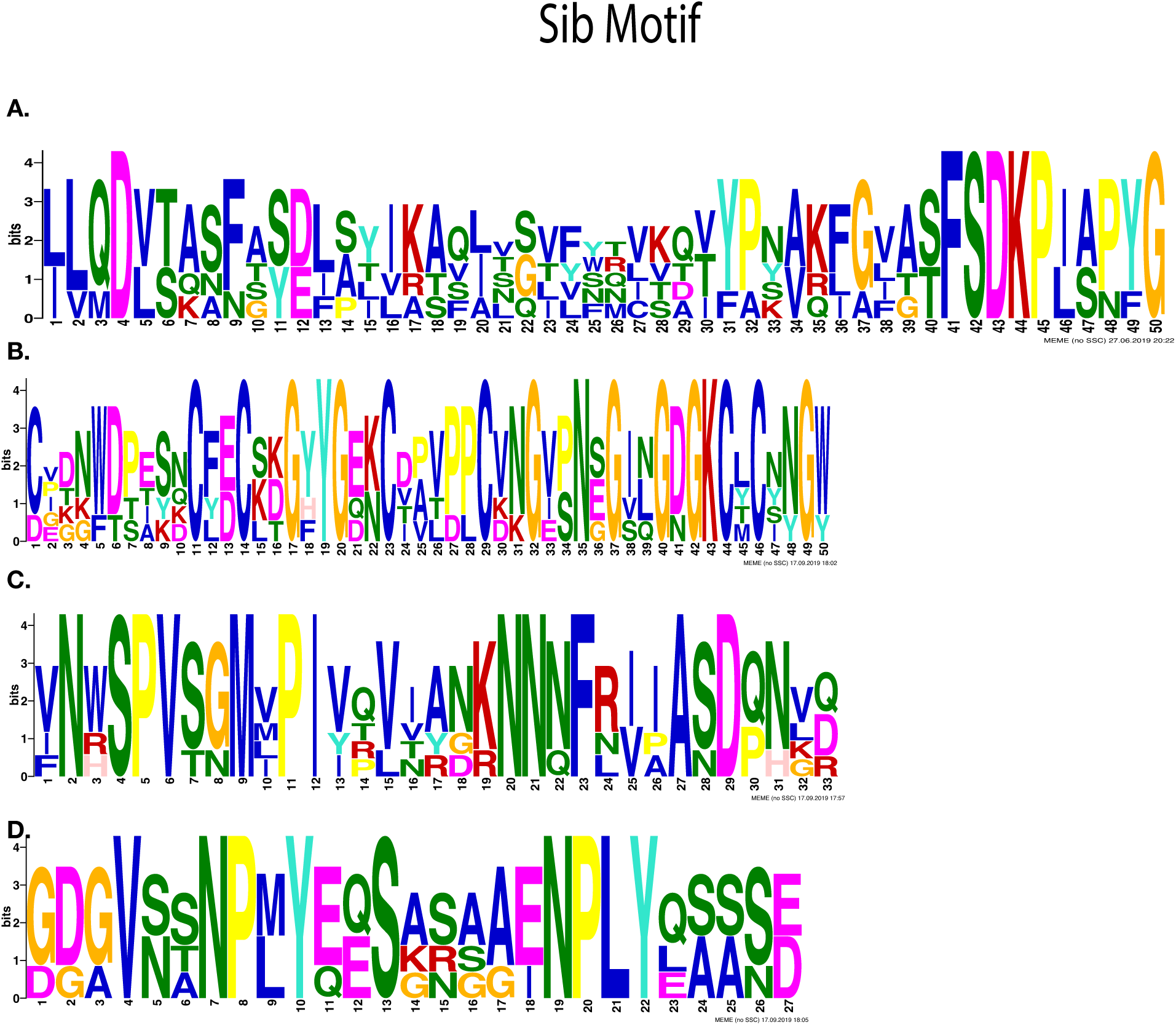
Most significantly enriched motifs in Sib protein, obtained by MEME. (**A**)-(**D**) Cation binding motifs (A) Possible binding motif of MIDAS and ADMIDAS, (**B**) Possible fifth position of MIDAS and ADMIDAS (C) Possible binding motif of SyMBS (**D**) NPXY motif

**Supplemental Table 1:**
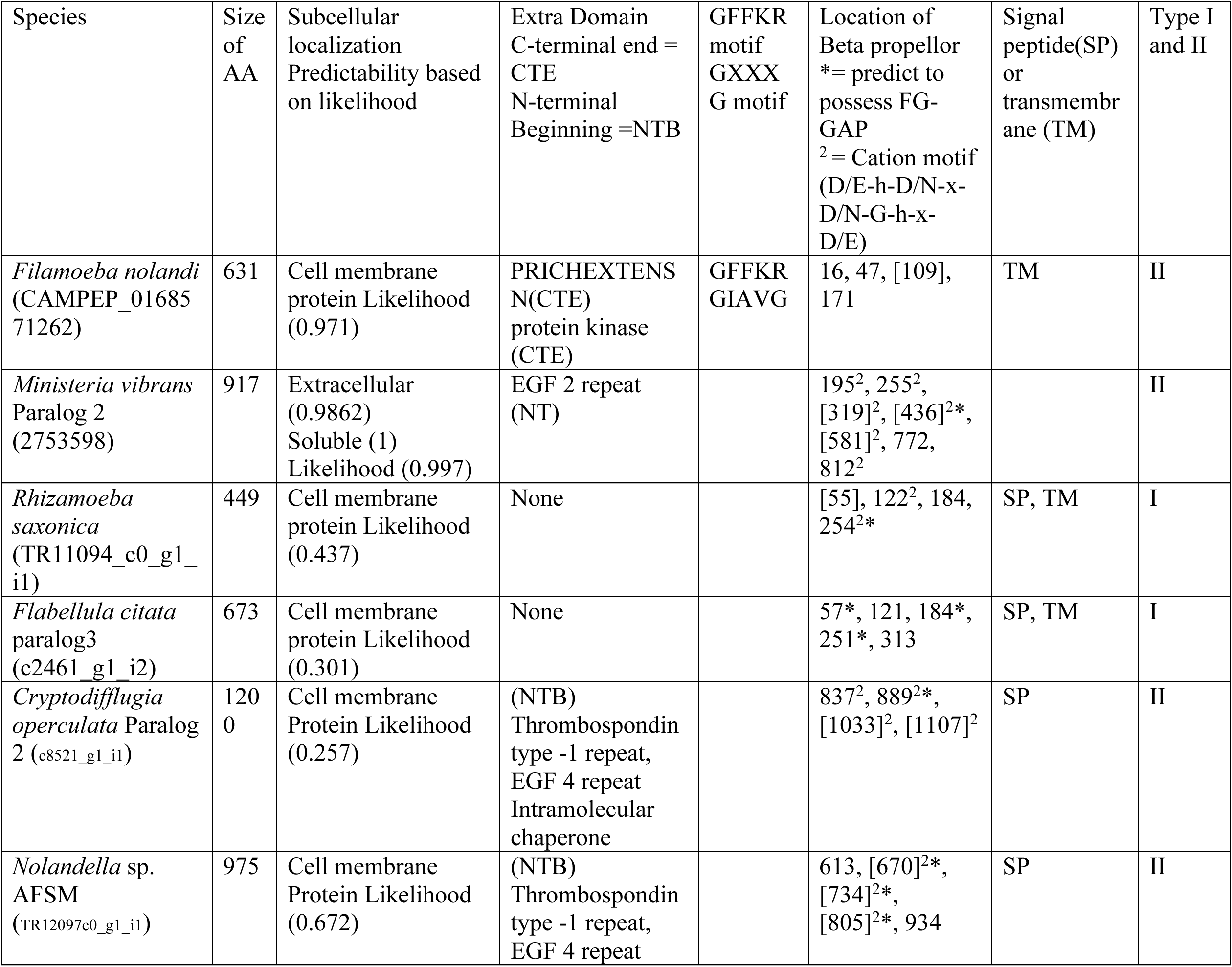
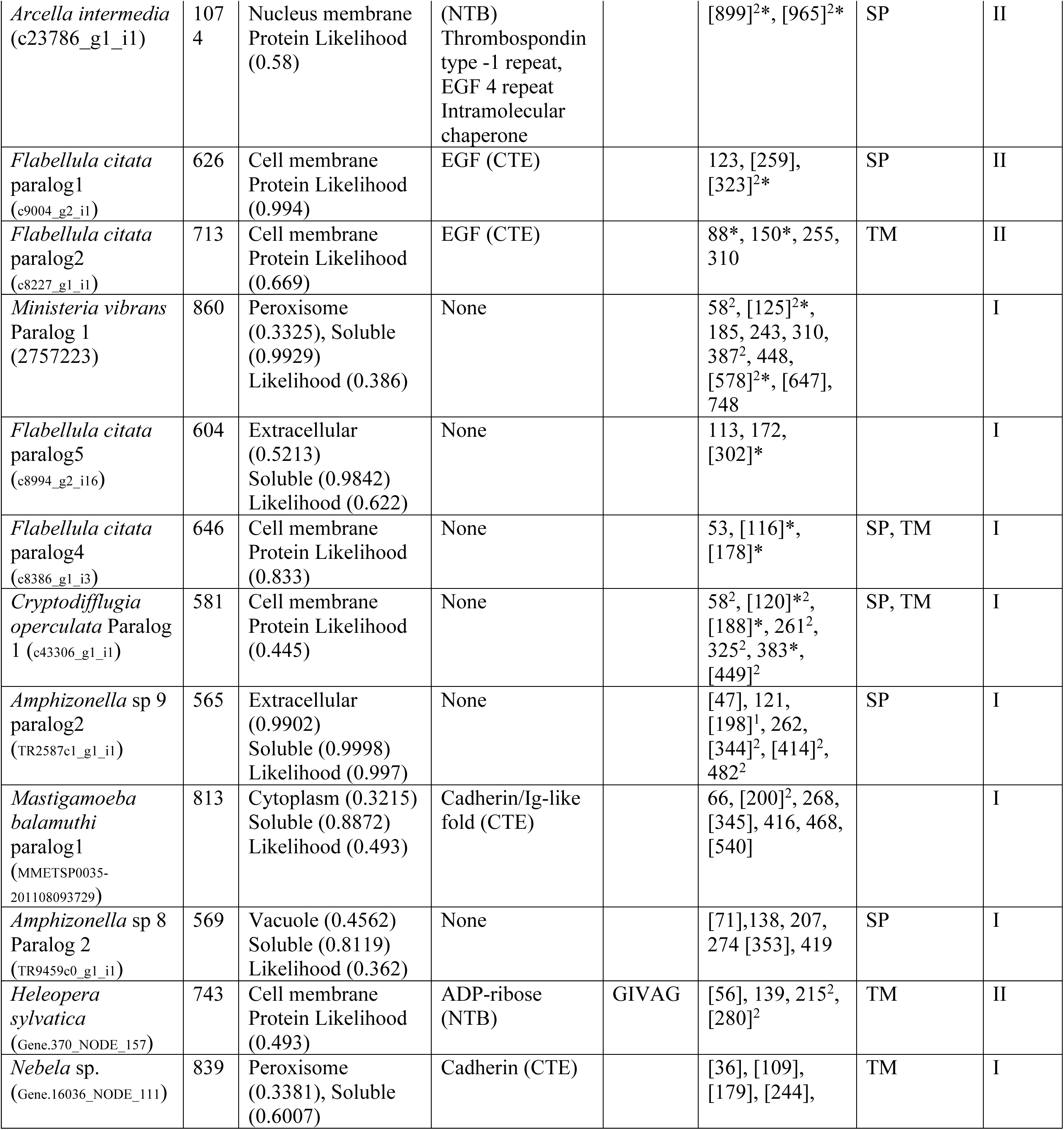
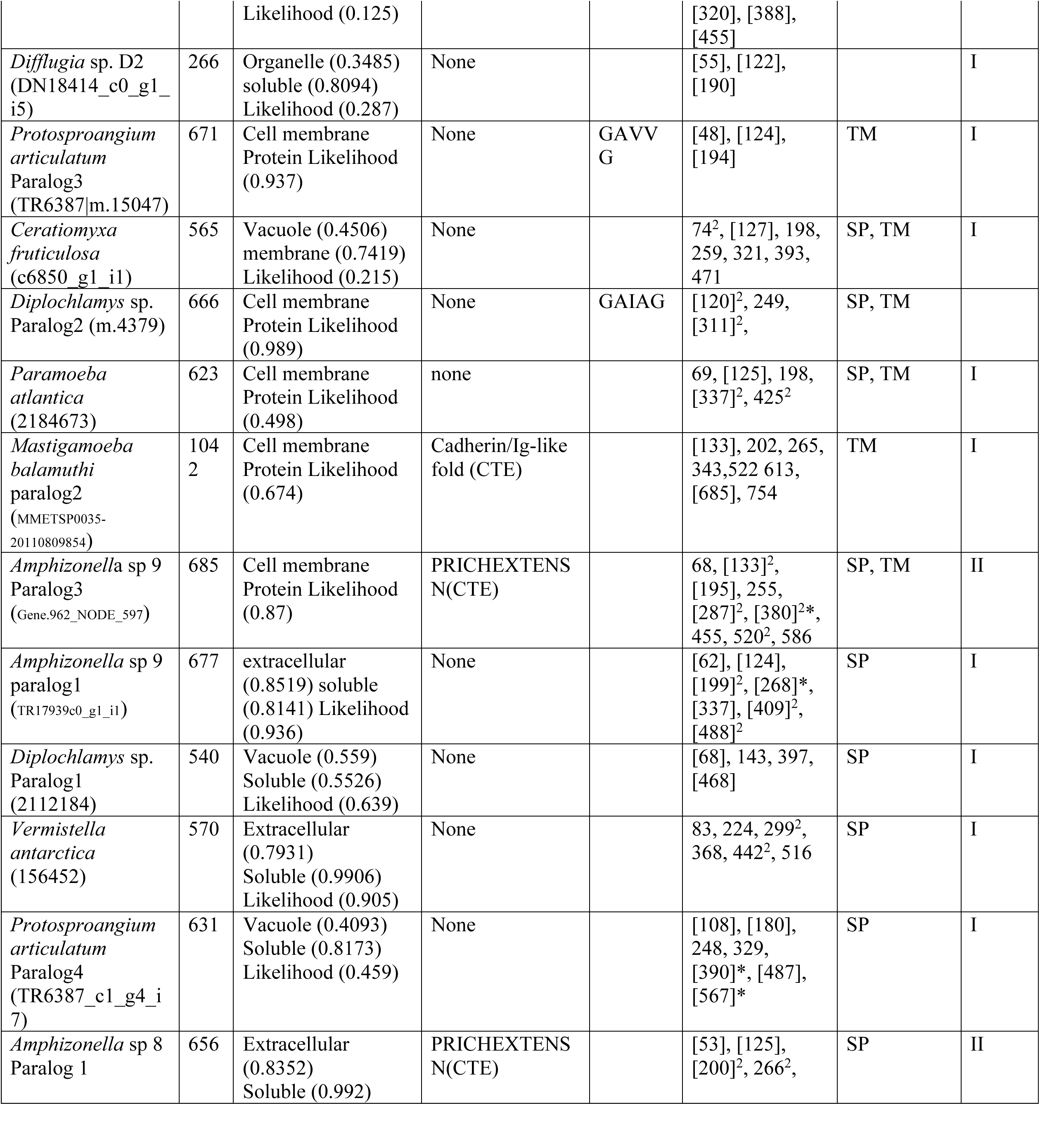
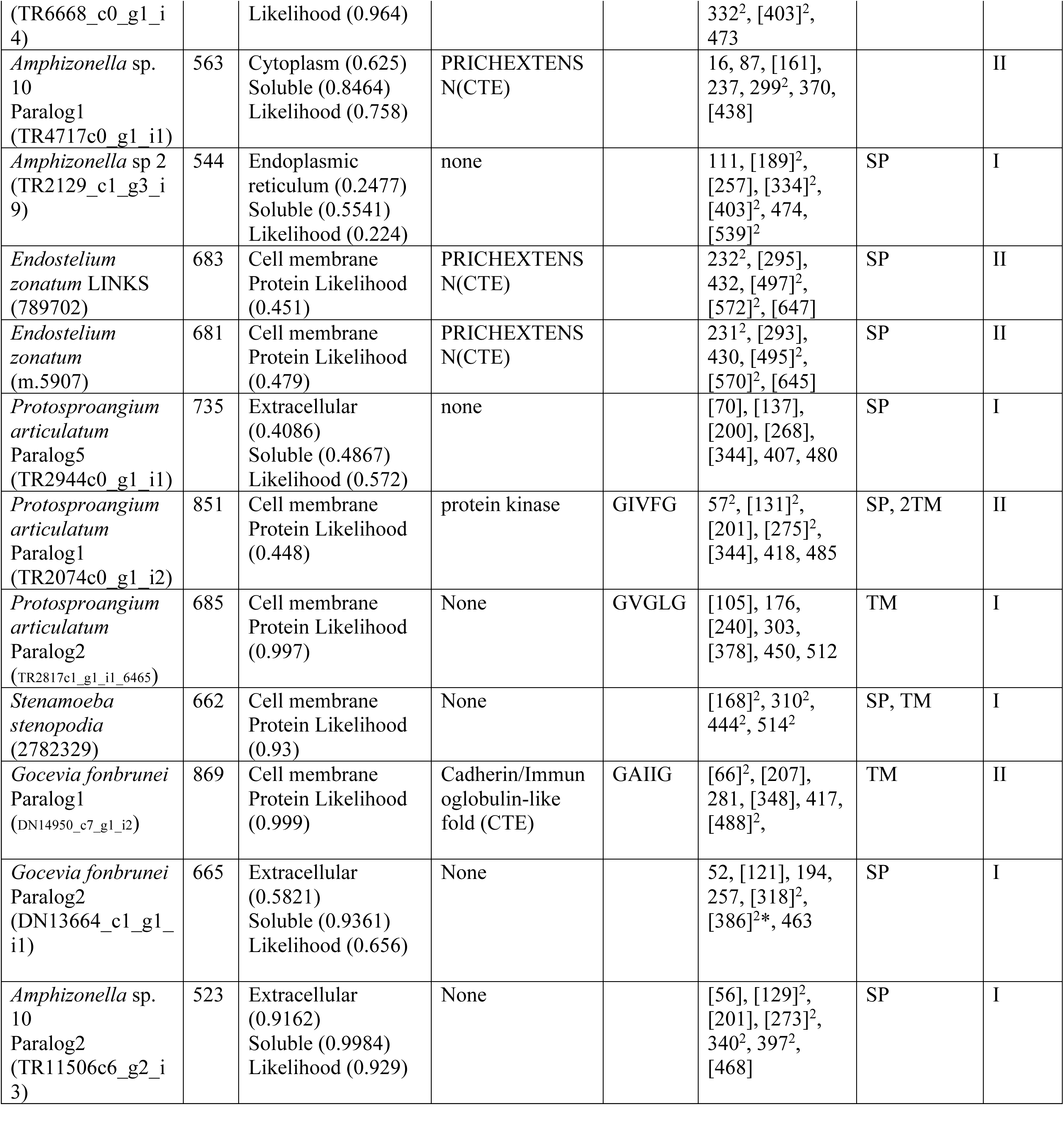
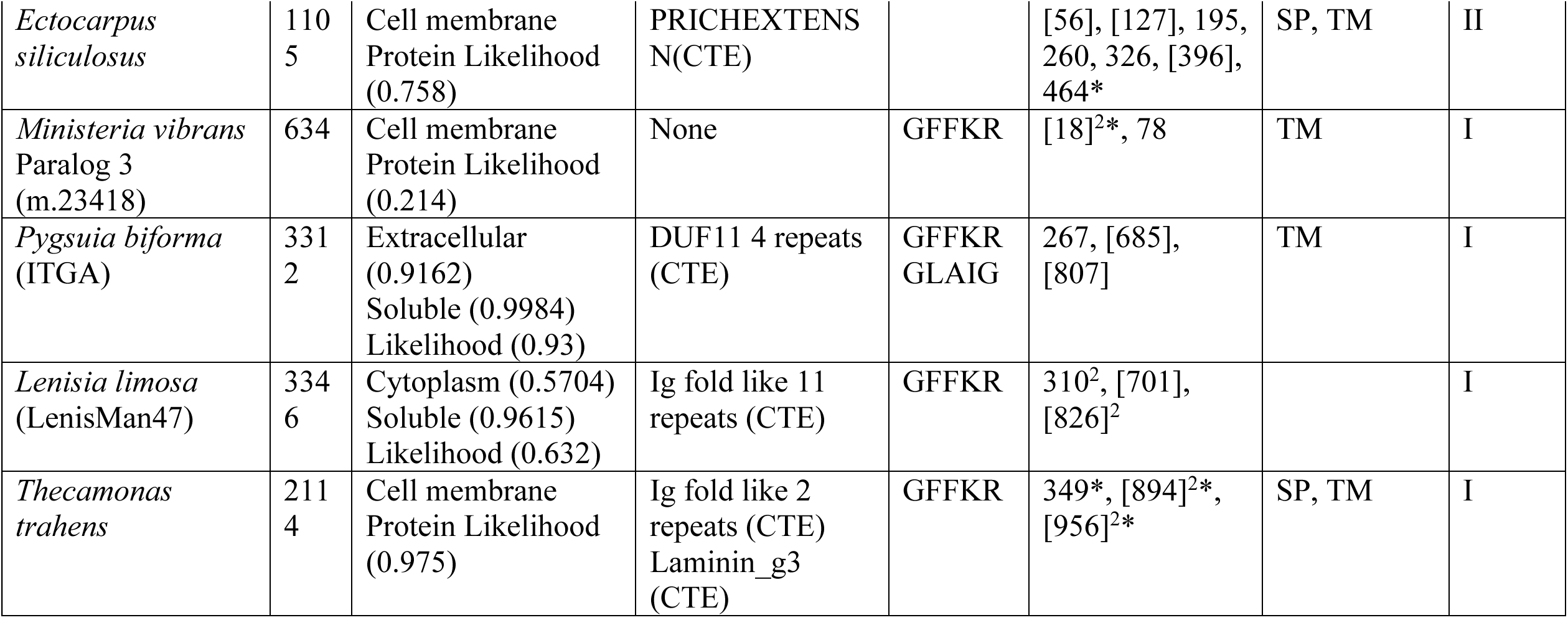
The table shows the list of Amorphea ITGA proteins characteristic including: length, beta propeller motifs, subcellular location, any extra domains attached to ITGA, c-terminal motif signal peptide, transmembrane regions and type of ITGA. The order of taxons are listed in accordance to the Figure 2. The subcellular localization is displayed with likelihood value in parentheses. The location of extra domains attached to ITGA are shown as the C-terminal end (CTE) or at the start of N-terminal (NTB). The presence of ITGA interacting motif with ITGB is shown as GFFKR and GXXXG. The location of beta propeller motifs performed by MEME-suite is displayed. The presence of FG-GAP motifs are shown in asterixis. The location of beta motifs that corresponds to IPRSCAN output are highlighted by parentheses. The presence of consensus cation motifs are displayed in number 2 superscripts.

**Supplemental Table 2:**
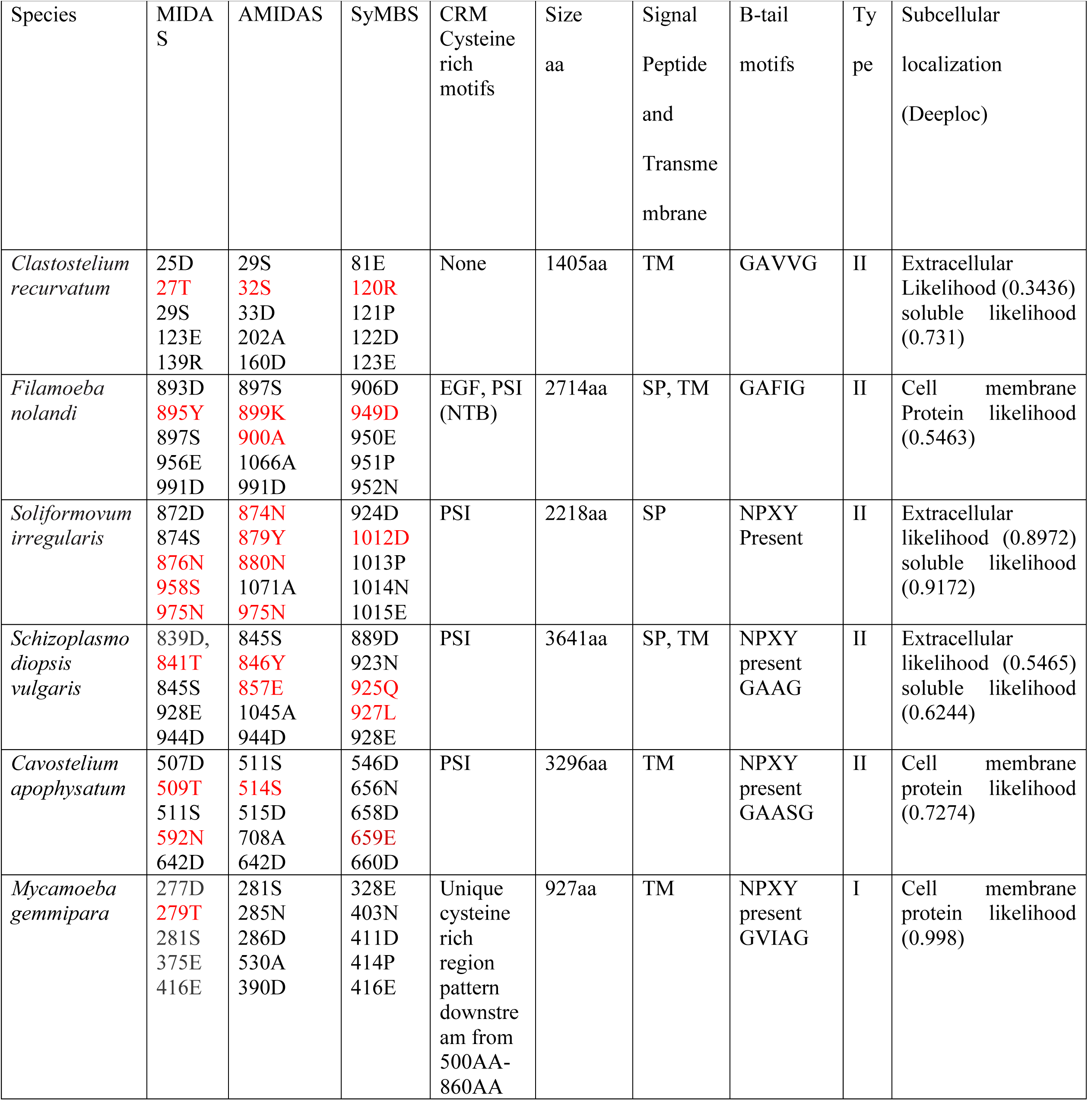
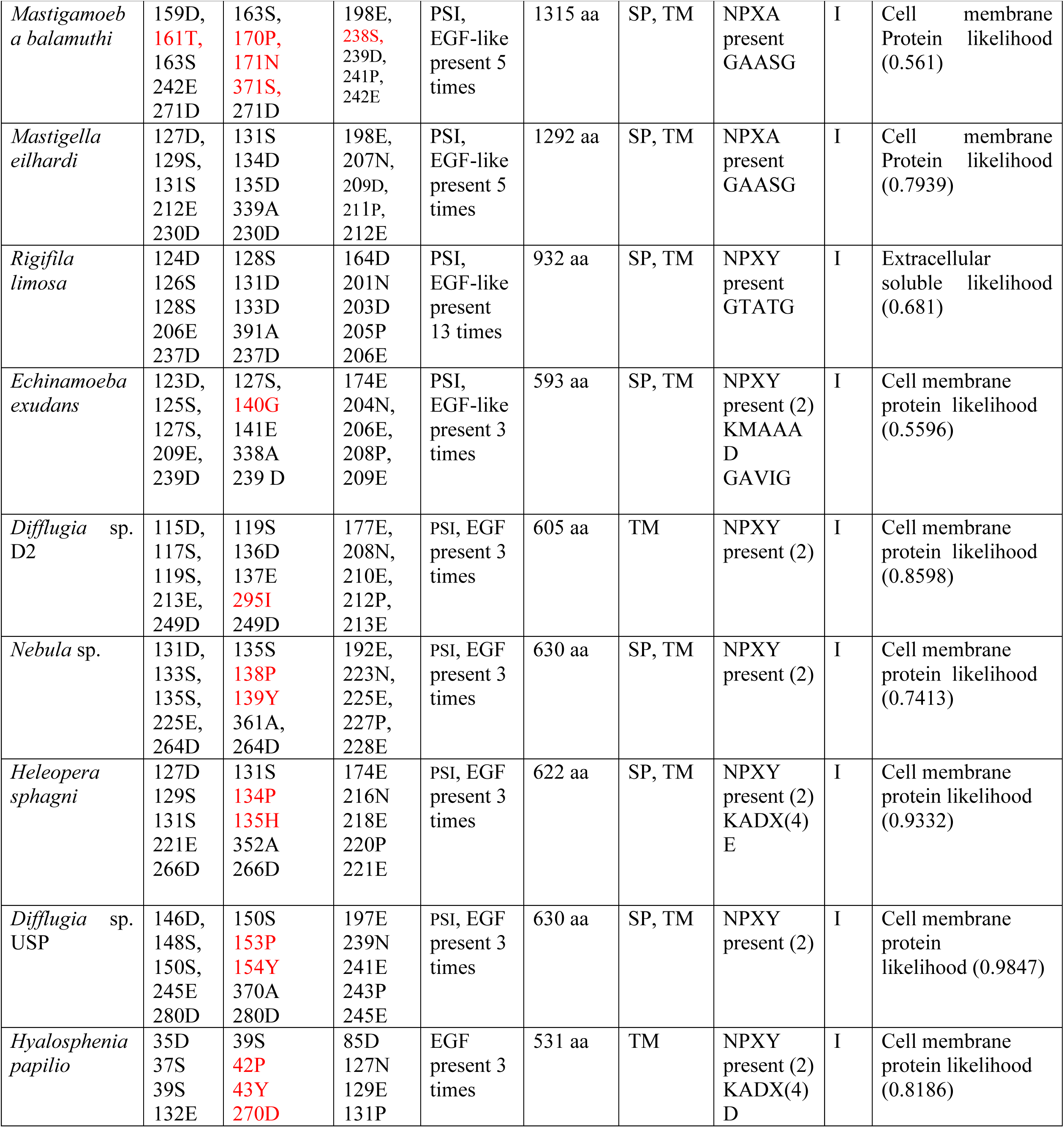
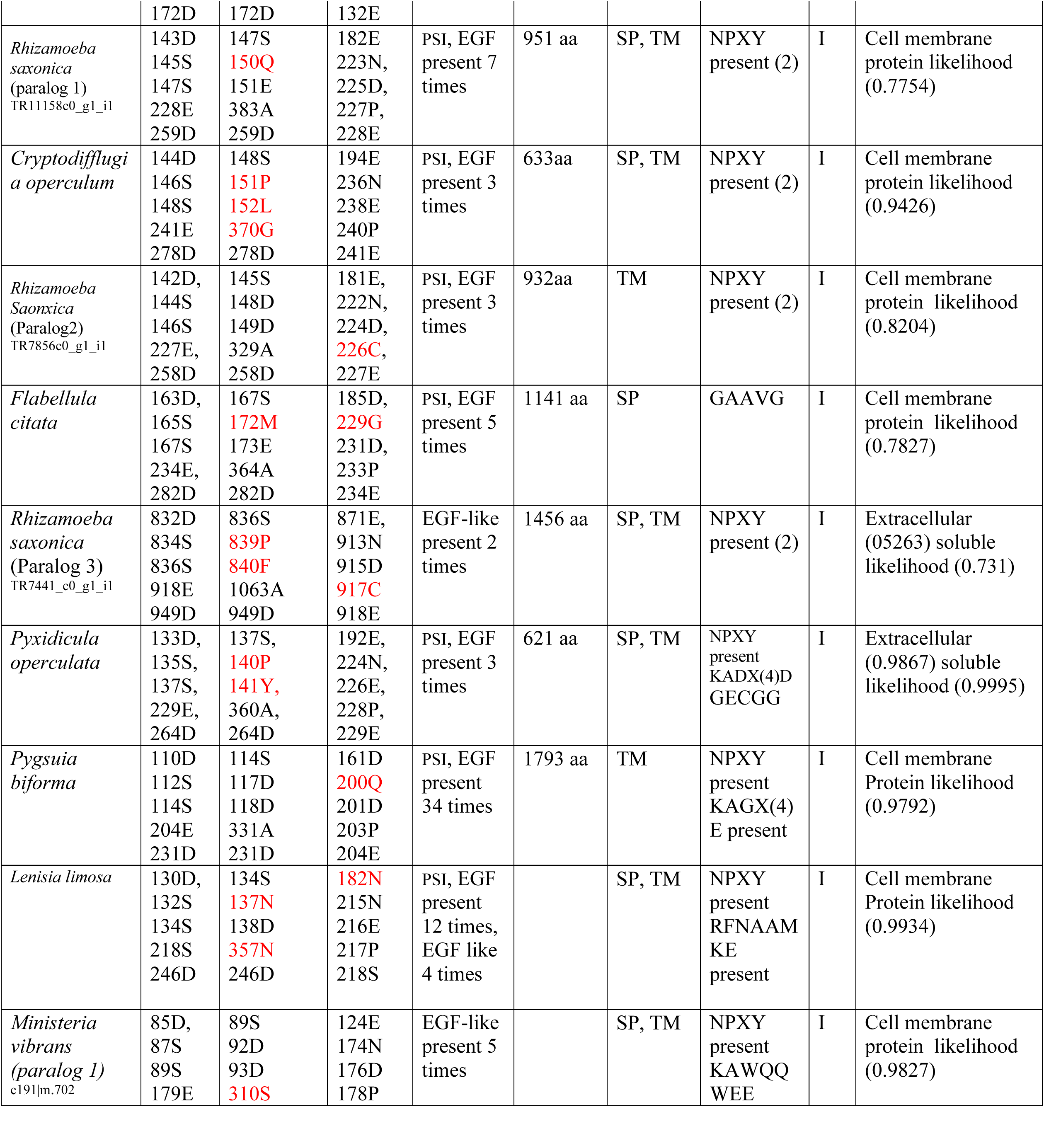
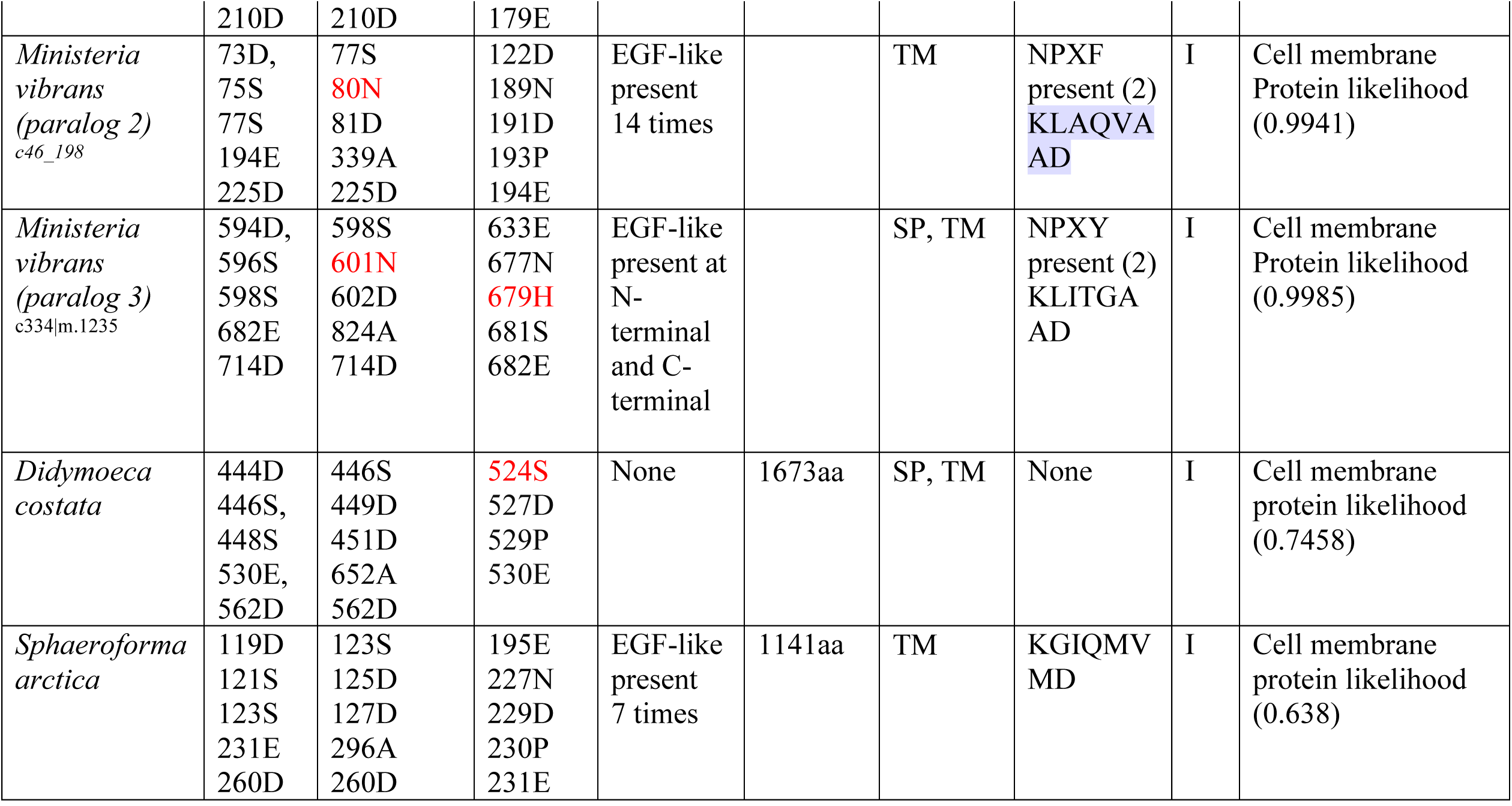
The table shows ITGB motifs, subcellular localizations, signal peptide, transmembrane and extra domains attached to ITGB protein. The order of taxons are listed in accordance to the Figure 3. For ITGB motifs of MIDAS, AMIDAS and SyMBS, any non-canonical amino acids were highlighted in red colour, cysteine rich motifs are either shown as EGF or PSI domains. The presence of C-terminal motifs are shown in KLXXXD or NPXY/F and GXXXG. The parentheses indicate number of motifs present in ITGB.

**Supplemental Table 3:**
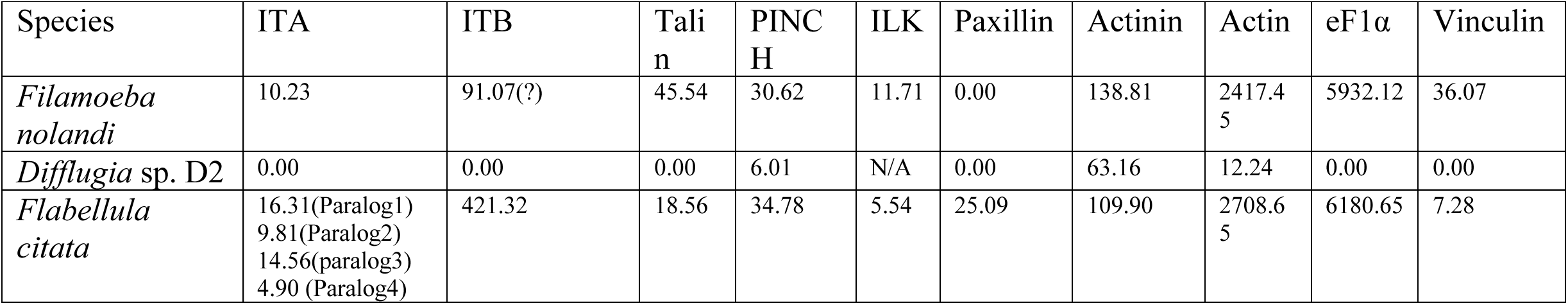
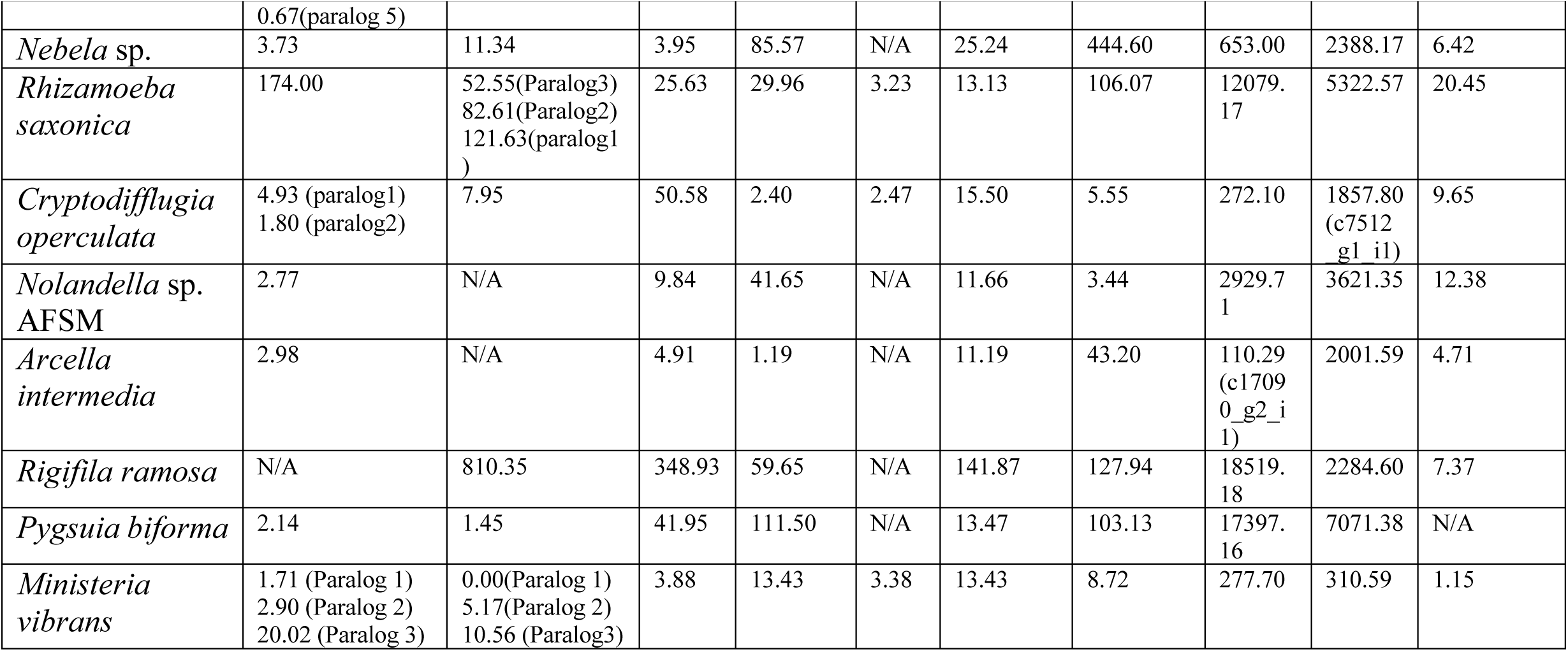
This table shows Rsem value of protists that have either a complete set of IMAC, integrins with extra domains or unexpected discovery of an integrin protein. The values are shown in transcripts per million (TPM).

**Supplemental Table 4:** A). Summary of amoebozoan ITA type I table listing: number of paralogs, Beta propeller, FG-GAP and canonical motifs. The table also shows presence of signal peptide and transmembrane region, c-terminal interacting motif, GXXXG motif and Ig-fold like/Cadherin domain. B). Summary of amoebozoan ITA type II table listing: number of paralogs, Beta propeller, FG-GAP and canonical motifs. The table also shows presence of signal peptide and transmembrane region, c-terminal interacting motif, GXXXG motif and extra domain attached to ITA.

## Notes

### Competing Interest Statement

The authors have declared no competing interest.

