## Supplemental Text for "The integrin-mediated adhesome complex, essential to multicellularity, is present in the most recent common ancestor of animals, fungi, and amoebae"

SUPPLEMENTAL METHODS, NOTES, AND RESULTS

**Key resources Table**

| Resources | Sources | Identifier |
| --- | --- | --- |

**Data Examined**

| *Acanthamoeba castellanii* | [56] | https://doi.org/10.1186/gb-2013-14-2-r11 |
| --- | --- | --- |
| *Acanthamoeba pyroformis* | [57] | <https://doi.org/10.1186/s13062-016-0171-0> |
| *Acanthamoeba astronyxis* | PRJEB7687 | <https://www.ncbi.nlm.nih.gov/bioproject/PRJEB7687/> |
| *Acanthamoeba comandoni* | PRJNA354221 | <https://www.ncbi.nlm.nih.gov/bioproject/PRJNA354221> |
| *Acanthamoeba culbertsoni* | PRJEB7687 | <https://www.ncbi.nlm.nih.gov/bioproject/PRJEB7687> |
| *Acanthamoeba divionensis* | PRJEB7687 | <https://www.ncbi.nlm.nih.gov/bioproject/PRJEB7687> |
| *Acanthamoeba healyi* | PRJEB7687 | <https://www.ncbi.nlm.nih.gov/bioproject/PRJEB7687> |
| *Acanthamoeba lenticulate* | PRJNA374454 | <https://www.ncbi.nlm.nih.gov/bioproject/PRJNA374454> |
| *Acanthamoeba lugdunensis* | PRJEB7687 | <https://www.ncbi.nlm.nih.gov/bioproject/PRJEB7687> |
| *Acanthamoeba mauritaniensis* | PRJEB7687 | <https://www.ncbi.nlm.nih.gov/bioproject/PRJEB7687> |
| *Acanthamoeba pearcei* | PRJEB7687 | <https://www.ncbi.nlm.nih.gov/bioproject/PRJEB7687> |
| *Acanthamoeba polyphaga* | PRJNA307312 | <https://www.ncbi.nlm.nih.gov/bioproject/PRJNA307312> |
| *Acanthamoeba quina* | PRJEB7687 | <https://www.ncbi.nlm.nih.gov/bioproject/PRJEB7687> |
| *Acanthamoeba rhysodes* | PRJEB7687 | <https://www.ncbi.nlm.nih.gov/bioproject/PRJEB7687> |
| *Acanthamoeba royreba* | PRJEB7687 | <https://www.ncbi.nlm.nih.gov/bioproject/PRJEB7687> |
| *Acytostelium ellipticum* | PRJEB14640 | <https://www.ncbi.nlm.nih.gov/bioproject/PRJEB14640> |
| *Acytostelium leptosomum* | PRJEB14640 | <https://www.ncbi.nlm.nih.gov/bioproject/PRJEB14640> |
| *Acytostelium subglobosum* | [58] | [https://doi.org/10.1186/s12864-015-1278-x](https://doi.org/10.1093/molbev/msx162) |
| *Amoeba proteus* | [14] | <https://doi.org/10.1093/molbev/msx162> |
| *Amphizonella* sp. 2 | [59] | https://doi.org/10.1016/j.cub.2019.01.078 |
| *Amphizonella* sp. 7 | [59] | https://doi.org/10.1016/j.cub.2019.01.078 |
| *Amphizonella* sp. 8 | [59] | https://doi.org/10.1016/j.cub.2019.01.078 |
| *Amphizonella* sp. 9 | [59] | https://doi.org/10.1016/j.cub.2019.01.078 |
| *Amphizonella* sp.10 | [59] | <https://doi.org/10.1093/molbev/msx162> |
| *Arcella gibbosa* | [14] | <https://doi.org/10.1093/molbev/msx162> |
| *Arcella vulgaris* | [59] | https://doi.org/10.1016/j.cub.2019.01.078 |
| *Armaparvus languidus* | MMETSP0420 | <https://www.imicrobe.us/project/view/304> |
| *Cavenderia deminutiva* | PRJEB14640 | <https://www.ncbi.nlm.nih.gov/bioproject/PRJEB14640> |
| *Cavostelium apophysatum* | [14] | <https://doi.org/10.1093/molbev/msx162> |
| *Centropyxis aerophyla* | [59] | https://doi.org/10.1016/j.cub.2019.01.078 |
| *Centropyxis sp.* | [59] | https://doi.org/10.1016/j.cub.2019.01.078 |
| *Ceratiomyxa fruticulosa* | [14] | <https://doi.org/10.1093/molbev/msx162> |
| *Ceratiomyxella tahitiensis* | [14] | <https://doi.org/10.1093/molbev/msx162> |
| *Clastostelium recurvatum* | [14] | <https://doi.org/10.1093/molbev/msx162> |
| *Clydonella* sp. | [60] | <https://doi.org/10.1016/j.ympev.2016.03.029> |
| *Cochliopodium minus* | [14] | <https://doi.org/10.1093/molbev/msx162> |
| *Copromyxa protea* | [14] | <https://doi.org/10.1093/molbev/msx162> |
| *Coremiostelium polycephalum* | PRJEB14640 | <https://www.ncbi.nlm.nih.gov/bioproject/PRJEB14640> |
| *Cryptodifflugia operculata* | [14] | <https://doi.org/10.1093/molbev/msx162> |
| *Cunea* sp. [JDS-Ruffled] | [14] | <https://doi.org/10.1093/molbev/msx162> |
| *Cunea* sp. [MMETSP0417] | MMETSP0417 | https://data.iplantcollaborative.org/dav/iplant/commons/community_released/imicrobe/projects/104/samples/1830/ |
| *Cyclopyxis* sp. | [59] | https://doi.org/10.1016/j.cub.2019.01.078 |
| *Dermamoeba algensis* | [14] | <https://doi.org/10.1093/molbev/msx162> |
| *Dictyostelium citrinum* | [61] | https://doi.org/10.1093/molbev/msn088 |
| *Dictyostelium discoideum* | [62] | https://doi.org/10.1038/nature03481 |
| *Dictyostelium firmibasis* | [63] | https://doi.org/10.1006/plas.1998.1385 |
| *Dictyostelium intermedium* | PRJNA45879 | <https://www.ncbi.nlm.nih.gov/bioproject/PRJNA45879> |
| *Dictyostelium purpureum* | [64] | <https://doi.org/10.1006/10.1186/gb-2011-12-2-r20> |
| *Difflugia bryophila* | [59] | https://doi.org/10.1016/j.cub.2019.01.078 |
| *Difflugia* sp. | [14] | <https://doi.org/10.1093/molbev/msx162> |
| *Difflugia* sp. D2 | [59] | https://doi.org/10.1016/j.cub.2019.01.078 |
| *Diplochlamys* sp. | [14] | <https://doi.org/10.1093/molbev/msx162> |
| *Dracoamoeba jomungandrii* | [57] | https://doi.org/10.1186/s13062-016-0171-0 |
| *Echinamoeba exudans* | [14] | <https://doi.org/10.1093/molbev/msx162> |
| *Echinosteliopsis oligospora* | [14] | <https://doi.org/10.1093/molbev/msx162> |
| *Echinostelium bisporum* | [14] | <https://doi.org/10.1093/molbev/msx162> |
| *Echinostelium minutum* | [14] | <https://doi.org/10.1093/molbev/msx162> |
| *Endostelium zonatum* [PRA-191] | [57] | <https://doi.org/10.1186/s13062-016-0171-0> |
| *Endostelium zonatum* LINKS | [14] | <https://doi.org/10.1093/molbev/msx162> |
| *Entamoeba dispar* | PRJNA12916 | https://www.ncbi.nlm.nih.gov/bioproject/12916 |
| *Entamoeba histolytica* | [65] | https://doi.org/10.1038/nature03291 |
| *Entamoeba invadens* | [66] | https://doi.org/10.1186/gb-2013-14-7-r77 |
| *Entamoeba moshkovskii* | PRJNA12921 | https://www.ncbi.nlm.nih.gov/bioproject/12921 |
| *Entamoeba nuttalli* | [67] | <https://doi.org/10.1016/j.molbiopara.2007.02.006> |
| *Filamoeba nolandi* | [MMETSP0413](https://www.imicrobe.us/sample/view/1828) | https://www.imicrobe.us/assembly/view/293 |
| *Flabellula citata* | [14] | <https://doi.org/10.1093/molbev/msx162> |
| *Flamella aegyptia* | [14] | <https://doi.org/10.1093/molbev/msx162> |
| *Gocevia fonbrunei* | [14] | <https://doi.org/10.1093/molbev/msx162> |
| *Gocevia fonbrunei*_Tekle_2016 | [60] | <https://doi.org/10.1016/j.ympev.2016.03.029> |
| *Grellamoeba robusta* | [14] | <https://doi.org/10.1093/molbev/msx162> |
| *Heleopera sphagni* | [59] | https://doi.org/10.1016/j.cub.2019.01.078 |
| *Heleopera sylvatica* | [59] | https://doi.org/10.1016/j.cub.2019.01.078 |
| *Heterostelium multicystogenum* | PRJNA495730 | <https://www.ncbi.nlm.nih.gov/bioproject/PRJNA495730> |
| *Heterostelium pallidum* | [61] | http://dictybase.org/ |
| *Hyalosphenia elegance* | [59] | https://doi.org/10.1016/j.cub.2019.01.078 |
| *Hyalosphenia papilio* | [59] | https://doi.org/10.1016/j.cub.2019.01.078 |
| *Idionectes vortex* | PRJNA531640 | <https://www.ncbi.nlm.nih.gov/bioproject/?term=Idionectes%20vortex> |
| *Lesquereusia* sp. | [59] | https://doi.org/10.1016/j.cub.2019.01.078 |
| *Lingulamoeba* sp. | [14] | <https://doi.org/10.1093/molbev/msx162> |
| *Luapelamoeba arachisporum* | [57] | https://doi.org/10.1186/s13062-016-0171-0 |
| *Luapelamoeba hula* | [57] | https://doi.org/10.1186/s13062-016-0171-0 |
| *Mastigamoeba abducta* | [14] | <https://doi.org/10.1093/molbev/msx162> |
| *Mastigamoeba balamuthi* (genome) | [33] | <http://doi.org/10.1073/pnas.1219590110> |
| *Mastigamoeba balamuthi* (transcriptome) | SRA TBD | To be provided |
| *Mastigella eilhardi* | [14] | <https://doi.org/10.1093/molbev/msx162> |
| *Mayorella cantabrigiensis* | [14] | <https://doi.org/10.1093/molbev/msx162> |
| *Micriamoeba* sp. | [14] | <https://doi.org/10.1093/molbev/msx162> |
| *Microchlamys* sp. | [59] | https://doi.org/10.1016/j.cub.2019.01.078 |
| *Mycamoeba gemmipara* | SRA TBD | https://doi.org/10.1111/jeu.1235 |
| *Nebela* sp. | [59] | https://doi.org/10.1016/j.cub.2019.01.078 |
| *Nematostelium gracile* | [14] | <https://doi.org/10.1093/molbev/msx162> |
| *Netzelia oviformis* | [59] | https://doi.org/10.1016/j.cub.2019.01.078 |
| *Nolandella* sp*.* | [14] | <https://doi.org/10.1093/molbev/msx162> |
| *Ovalopodium desertum* | [14] | <https://doi.org/10.1093/molbev/msx162> |
| *Paradermamoeba levis* | [14] | <https://doi.org/10.1093/molbev/msx162> |
| *Paramoeba aestuarina* | MMETSP0161_2 | https://www.imicrobe.us/assembly/view/205 |
| *Paramoeba atlantica* | MMETSP0151_2 | https://www.imicrobe.us/assembly/view/210 |
| *Parvamoeba rugata* | [14] | <https://doi.org/10.1093/molbev/msx162> |
| *Parvamoeba monoura* | [32] | <https://doi.org/10.1016/j.ympev.2016.03.029> |
| *Pellita catalonica* | [14] | <https://doi.org/10.1093/molbev/msx162> |
| *Pelomyxa* sp. | [14] | <https://doi.org/10.1093/molbev/msx162> |
| *Phalansterium solitarium* | [14] | <https://doi.org/10.1093/molbev/msx162> |
| *Physarum polycephalum* | [68] | https://dx.doi.org/10.1093/gbe/evv237 |
| *Pygsuia biforma* | [11] | <http://doi.org/10.1098/rspb.2013.1755> |
| *Planocarina carinata* | [59] | https://doi.org/10.1016/j.cub.2019.01.078 |
| *Polysphondylium pallidum* | [69] | http://dictybase.org/ |
| *Protacanthamoeba bohemica* | [14] | <https://doi.org/10.1093/molbev/msx162> |
| *Protosporangium articulatum* | [14] | <https://doi.org/10.1093/molbev/msx162> |
| *Protostelium nocturnum* | [14] | <https://doi.org/10.1093/molbev/msx162> |
| *Pyxidicula operculatum* | [59] | https://doi.org/10.1016/j.cub.2019.01.078 |
| *Rhizamoeba saonxica* | [14] | <https://doi.org/10.1093/molbev/msx162> |
| *Rhizomastix elongate* | [14] | <https://doi.org/10.1093/molbev/msx162> |
| *Rhizomastix libera* | [14] | <https://doi.org/10.1093/molbev/msx162> |
| *Ripella* sp. | [14] | <https://doi.org/10.1093/molbev/msx162> |
| *Sapocribrum chincoteaguense* | MMETSP0437 | https://www.imicrobe.us/assembly/view/314 |
| *Sappinia pedata* | [14] | <https://doi.org/10.1093/molbev/msx162> |
| *Schizoplasmodiopsis pseudoendospora* | [14] | <https://doi.org/10.1093/molbev/msx162> |
| *Schizoplasmodiopsis vulgaris* | [14] | <https://doi.org/10.1093/molbev/msx162> |
| *Speleostelium caveatum* | PRJNA495862 | <https://www.ncbi.nlm.nih.gov/bioproject/PRJNA495862> |
| *Soliformovum irregularis* | [14] | <https://doi.org/10.1093/molbev/msx162> |
| *Squamamoeba japonica* | [14] | <https://doi.org/10.1093/molbev/msx162> |
| *Stenamoeba limacine* | [14] | <https://doi.org/10.1093/molbev/msx162> |
| *Stenamoeba stenopodia* | [14] | <https://doi.org/10.1093/molbev/msx162> |
| *Stygamoeba regulata* | MMETSP0447 | https://www.imicrobe.us/assembly/view/316 |
| *Synstelium polycarpum* | PRJEB14640 | <https://www.ncbi.nlm.nih.gov/bioproject/PRJEB14640> |
| *Thecamoeba quadrilineata* | [60] | <https://doi.org/10.1016/j.ympev.2016.03.029> |
| *Thecamoeba* sp. | [14] | <https://doi.org/10.1093/molbev/msx162> |
| *Thecamoebidae* isolate [RHP1-1] | [14] | <https://doi.org/10.1093/molbev/msx162> |
| *Trichosphaerium* sp. (ATCC 40318) | MMETSP0405 | https://www.imicrobe.us/assembly/view/300 |
| *Tychosporium acutostipes* | [14] | <https://doi.org/10.1093/molbev/msx162> |
| *Vannella fimicola* | [14] | <https://doi.org/10.1093/molbev/msx162> |
| *Vannella robusta* | MMETSP0166 | https://www.imicrobe.us/?#/samples/2469 |
| *Vannella schaefferi* | MMETSP0417 | https://www.imicrobe.us/assembly/view/295 |
| *Vannella* sp. | MMETSP0168 | https://www.imicrobe.us/assembly/view/34 |
| *Vermamoeba* sp. | [14] | <https://doi.org/10.1093/molbev/msx162> |
| *Vermamoeba vermiformis* | [14] | <https://doi.org/10.1093/molbev/msx162> |
| *Vermistella antarctica* | [60] | <https://doi.org/10.1016/j.ympev.2016.03.029> |
| *Vexillifera bacillipedes* | [70] | <http://doi.org/10.1016/j.ympev.2014.08.011> |
| *Vexillifera* sp. | MMETSP0173 | https://www.imicrobe.us/#/samples/1757 |

**List of Stramenophile TAXA from MMETSP**

Alexandrium minutum	MOORE-via-CAMERA-MMETSP0328

Aureoumbra_lagunensis CCMP1509	MMETSP0890

Bolidomonas_sp RCC1657	MMETSP1321

Bolidomonas_sp RCC2347	MMETSP1320

Bolidomonas pacifica	MMETSP1319-RCC208

Cafeteria_sp CaronLabIsolate	MMETSP1104

Chattonella_subsalsa CCMP2191	MMETSP0947

Chrysocystis_fragilis CCMP3189	MMETSP1165 ****

Dictyocha_speculum CCMP1381	MMETSP1174

Dinobryon_sp UTEXLB2267	MMETSP0019

Fibrocapsa japonica-CCMP1661	MOORE-via-CAMERA-MMETSP1339

Florenciella_sp RCC1587	MMETSP1324	Stramenopiles

Florenciella parvula	MMETSP1323-RCC1693

Heterosigma_akashiwo CCMP2393	MMETSP0292

Love2098 non_described LOVEJOY CMP2098	MMETSP0990

Mallomonas_sp CCMP3275	MMETSP1167

Ochromonas_sp BG1	MMETSP1105

Paraphysomonas_bandaiensis CaronLabIsolate	MMETSP1103

Paraphysomonas_vestita GFlagA	MMETSP1107

Pelagomonas_calceolata CCMP1756	MMETSP0886

Pelagococcus_subviridis CCMP1429	MMETSP0882

Phaeomonas_parva CCMP2877	MMETSP1163

Pinguiococcus_pyrenoidosus CCMP2078	MMETSP1160

Pseudopedinella_elastica CCMP716	MMETSP1068

Rhizochromulina_marina cf CCMP1243	MMETSP1173

Spumella_elongata CCAP955 1	MMETSP1098

Thraustochytrium_sp LLF1b	MMETSP0198

Vaucheria_litorea CCMP2940	MMETSP0945

| **Reagents** |  |  |
| --- | --- | --- |
| 2-propanol(certified ACS)  Ethanol (pure) 200 proof | Sigma-aldrich | I9516-25ML |
| **Critical Commercial Assay Kits** |  |  |
| Nextera XT DNA Library Prep Kit | Illumina | FC-131-1096 |
| Nextera XT index kit v2 set A | Illumina | FC-131-2001 |
| Nextera XT index kit v2 set B | Illumina | FC-131-2002 |
| Nextera XT index kit v2 set C | Illumina | FC-131-2003 |
| Nextera XT index kit v2 set C | Illumina | FC-131-2003 |

| **Software and Algorithms** |  |  |
| --- | --- | --- |
| CD-HIT | [71] | <https://github.com/weizhongli/cdhit> |
| Trinity v2.4.0 | [72] | https://github.com/trinityrnaseq/trinityrnaseq/ |
| TransDecoder v 5.5.0 | [73] | https://transdecoder.github.io/ |
| TmHmm v 2.0 | [74] | http://www.cbs.dtu.dk/services/TMHMM/ |
| Bowtie2 v 2.3.4.3 | [75] | <https://sourceforge.net/projects/bowtie-bio/files/bowtie2/2.3.3.1> |
| Diamond v 0.9.25 | [76] | <https://github.com/bbuchfink/diamond> |
| OrthoMCL v 5.0 | [77] | http://orthomcl.org/orthomcl/ |
| Trimmomatic v 0.35 | [78] | https://github.com/timflutre/trimmomatic |
| Mafft-Linsi v 7 | [79] | https://mafft.cbrc.jp/alignment/server/ |
| Bmge v 1.12 | [80] | ftp://ftp.pasteur.fr/pub/GenSoft/projects/BMGE |
| Sequencer v 5.4.6. |  | http://www.genecodes.com/ |
| BLAST 2.2.30+ |  | https://blast.ncbi.nlm.nih.gov/ |
| IQtree v 1.5.5 | [81] | http://www.iqtree.org |
| Rsem | [82] | https://github.com/deweylab/RSEM |
| SignalIP 5.0 | [83] | http://www.cbs.dtu.dk/services/SignalP/ |
| Oyster River Protocol v 2.1.1 | [84] | https://oyster-river-protocol.readthedocs.io/en/latest/ |
| Meme-Suite v 5.0.4 | [85] | http://meme-suite.org/index.html |
| DeepLoc -1.0 | [86] | http://www.cbs.dtu.dk/services/DeepLoc/ |
| InterProScan 5.27-66.0 | [23] | https://www.ebi.ac.uk/interpro/search/sequence-search |
| ETE3 | [87] | <http://etetoolkit.org/documentation/ete-view/> |
| Pfam | [88] | <https://pfam.xfam.org/> |
| Mcl | [89] | <https://micans.org/mcl/> |

**Contact for Reagent and resources sharing**

***Sib (similar to integrin beta)* proteins found in dictyostelids** | **Sib Proteins comparison with ITB-like**

SibA, a cell adhesion molecule that binds to phagocytic particles similar to ITB, was discovered in *Dictyostelium discoideum* [45]. Sib proteins (A to E) are capable of binding to talin and SibC regulates cell adhesion [46]. However their protein structure is different from the canonical ITB in a lack of three cation binding motifs as well as a cysteine rich region at the C-terminus [26]. Although the varioseans, described above, are phylogenetically close to dictyostelids their type two ITB-like proteins with three cation motifs are distinct from Sib proteins, though with a great variation in amino acid sequences. Interestingly, we did not observe Sib proteins outside of the Dictyostelia clade, suggesting that they evolved recently in the dictyostelids. The protein architecture of SibA has a similar composition to metazoan ITB solely with a GXXXG motif in the transmembrane region, but it is observed in type II ITB (see supplemental table 2). In addition, LamG3, vWD domains in type II ITB have never been observed in Sib.

**Supplemental Figure 1:** Repertoire of IMAC including filamin and tensin in Amoebozoa: names of IMAC proteins are listed at the top and names of amoebozoan taxa are listed on side of the map. Arrows indicates amoebozoan species that have nearly complete set of IMAC proteins. Dark blue indicates the presence of a canonical IMAC protein, white indicates the absence of an IMAC protein and light blue indicates a truncated form of IMAC protein.

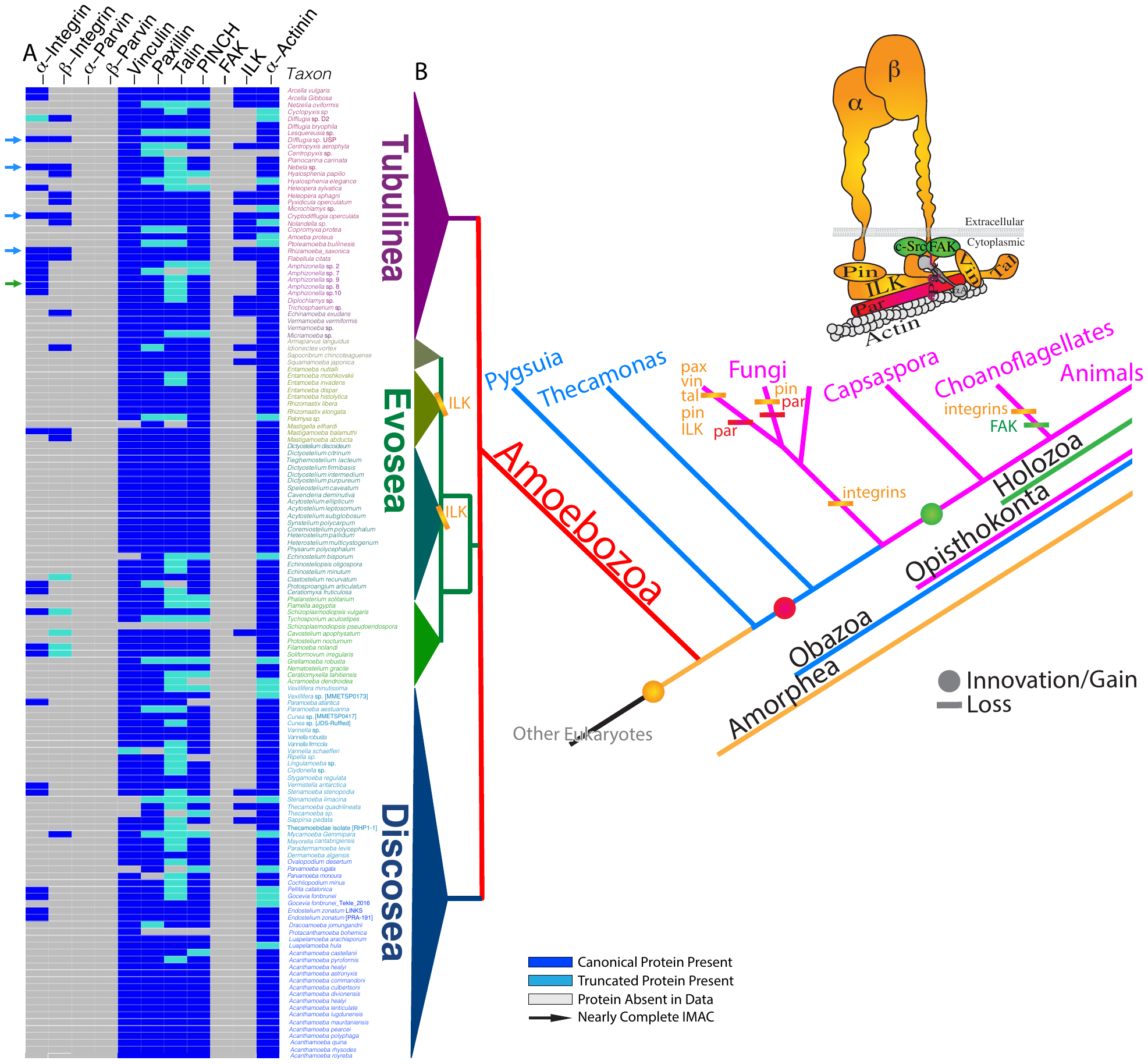
**Supplemental Figure 2:** Repertoire of IMAC in only amoebozoan species that have either integrin alpha or beta. Names of IMAC proteins are listed at the top and names of amoebozoan taxa are listed on side of the map. Arrows indicates amoebozoan species that have nearly complete set of IMAC proteins. Dark blue indicates the presence of a canonical IMAC protein, white indicates the absence of an IMAC protein and light blue indicates a truncated form of IMAC protein

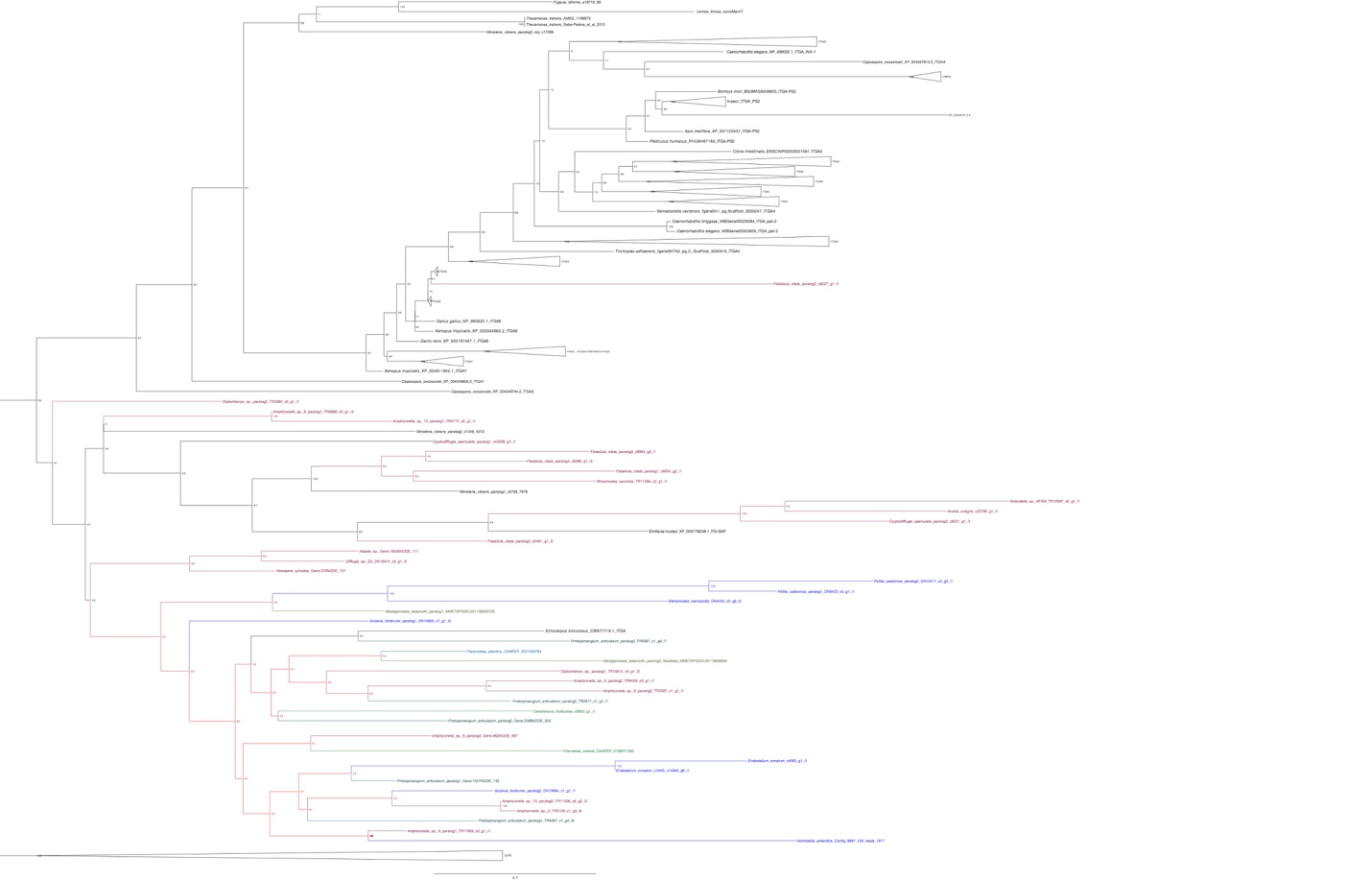
**Supplemental Figure 3:** Maximum likelihood tree of ITA homolog and other cell adhesion molecules that contain beta-propellers. Bootstrap values ≥50 are shown on the branch points. Phylogenetic tree was built with IQtree v1.5.5 under under the LG+C60+F+G model of protein evolution ML bootstrap (MLBS) (1000 ultrafast BS reps) values respectively. The protein sequences were aligned by Mafft with the parameter linsi, maxiterate 1000 and local pair. BMGE was used to mask the alignment with the parameter of gap penalty of 0.8. ITA homolog phylogenetic tree are rooted with Linkin protein. Amoebozoan ITA genes are mostly concentrated on a single clade (colored) except for the *Flabellula citata* paralog.

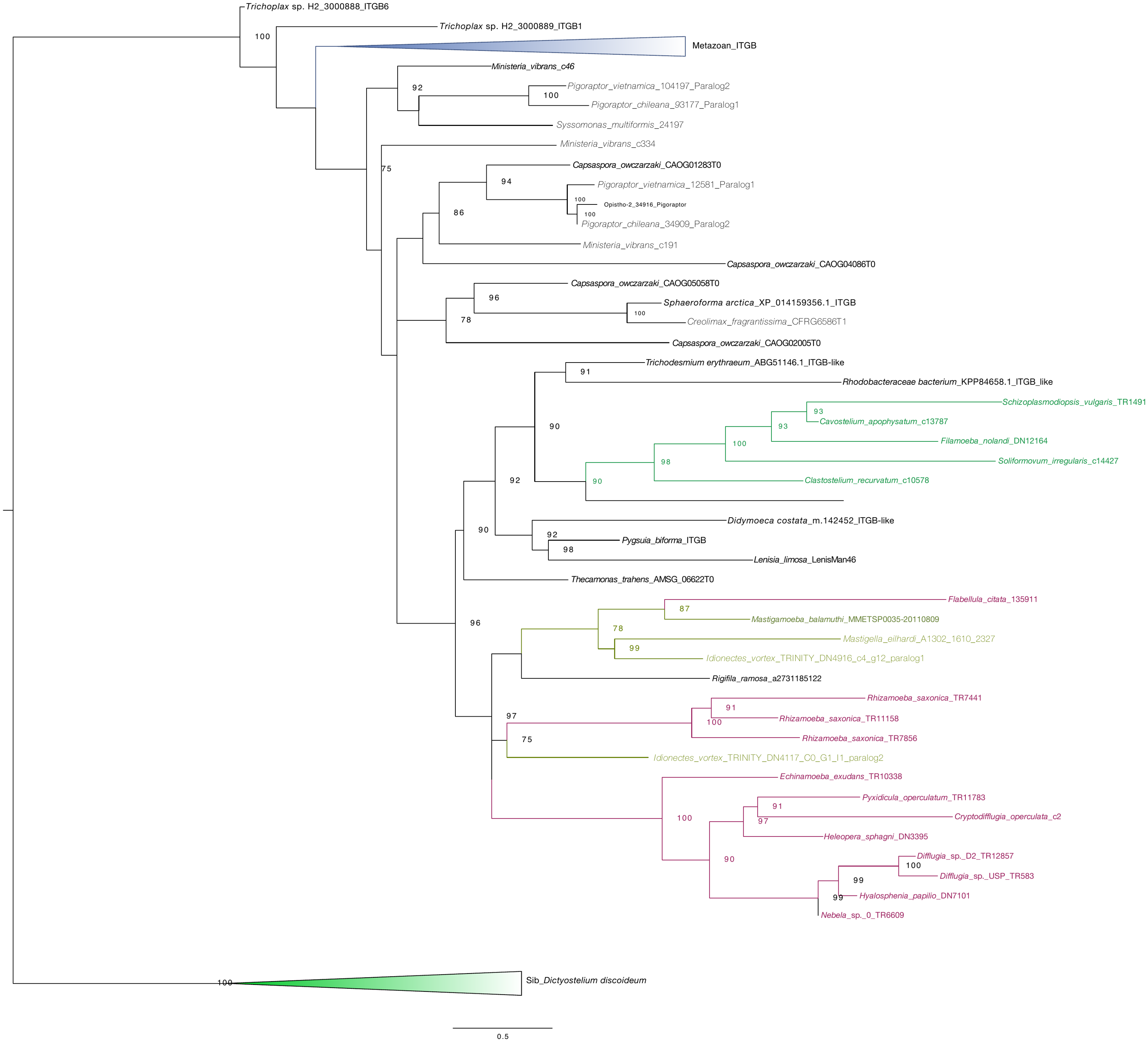
**Supplemental Figure 4:** Maximum likelihood tree of ITB homolog and Sib protein. Bootstrap values ≥50 are shown on the branch points. Phylogenetic tree was built with IQtree v1.5.0 under the LG+C60+F+G model of protein evolution ML bootstrap (MLBS) (1000 ultrafast BS reps) values respectively. The protein sequences were aligned by Mafft with the parameter linsi, maxiterate 1000 and local pair. BMGE was used to mask the alignment with the parameter of gap penalty of 0.6. ITB homolog phylogenetic tree are rooted with Sib protein. Amoebozoan ITB genes are mostly concentrated on a single clade (colored).

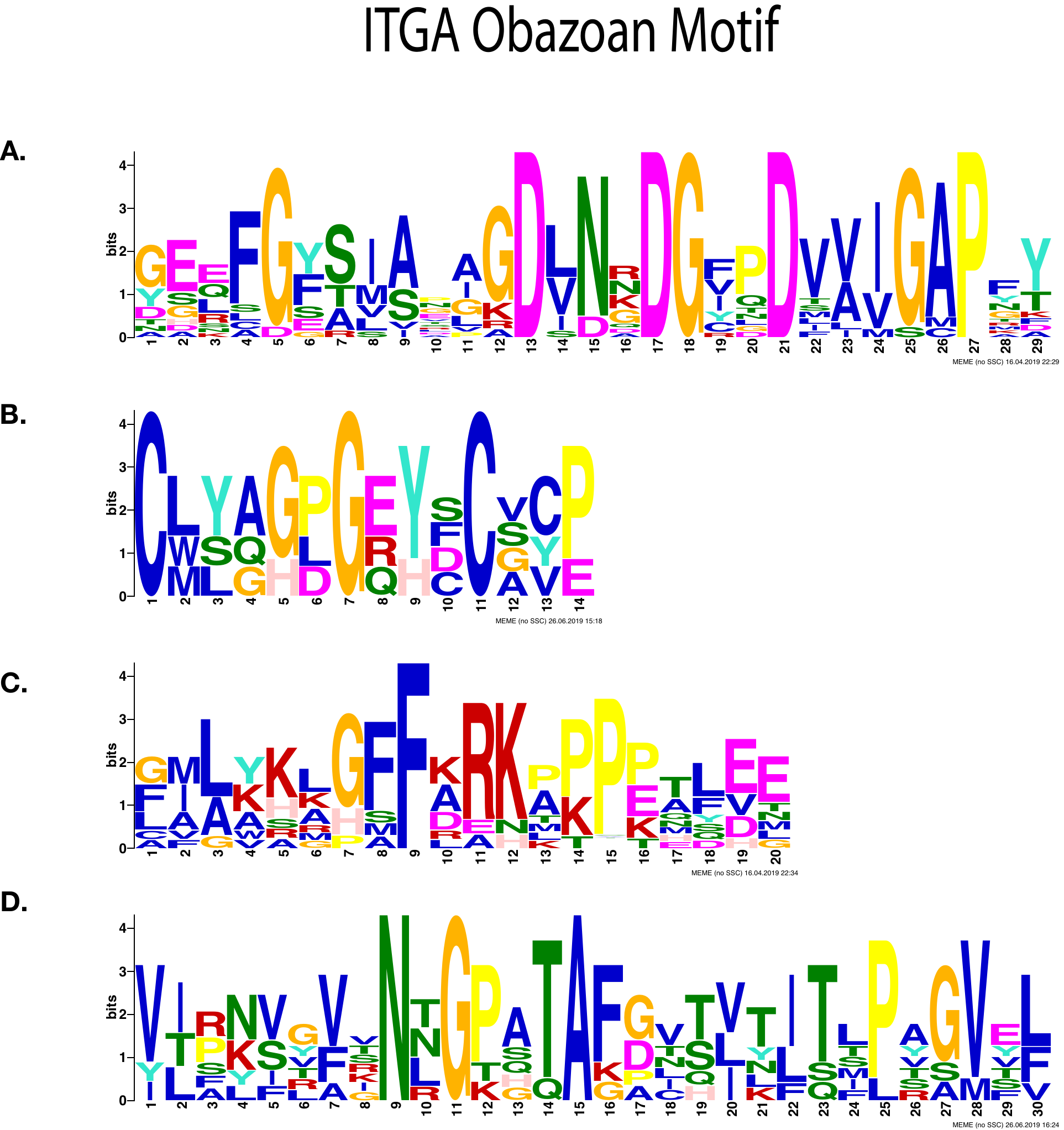

**Supplemental Figure 6:** Most significantly enriched motifs in Obazoan ITA, obtained by MEME-suite. (**A**) Consensus cation motif and FG-GAP obtained by RNA-seq for an Obazoan ITA, (**B**) EGF motif found exclusively in *Ministeria vibrans* (**C**) GFFKR motif found in Obazoa, (**D**) immunoglobulin like motif found in Obazoa.

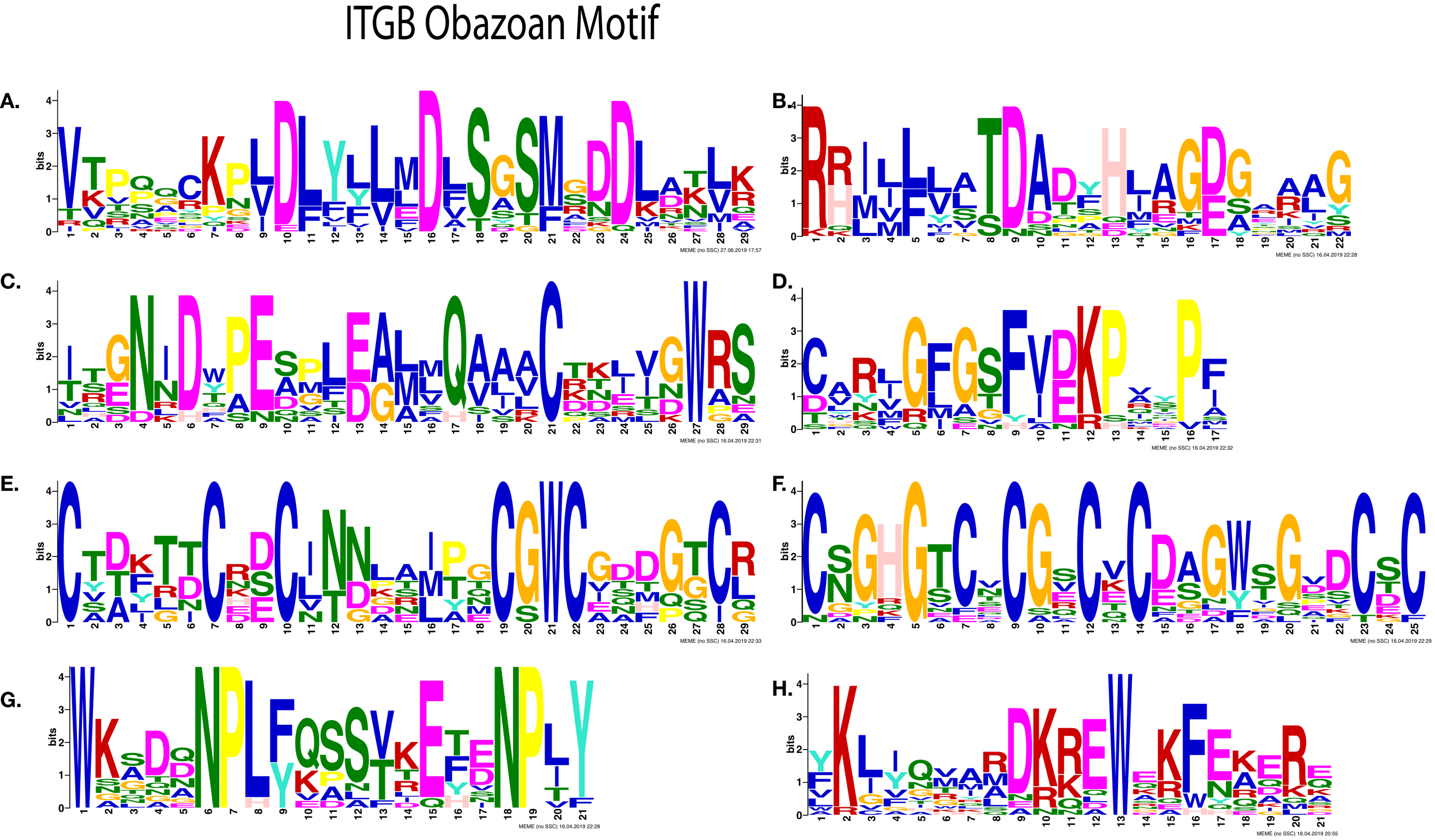
**Supplemental Figure 7:** Most significantly enriched motifs in Obazoan ITB, obtained by MEME. (**A**-**D**) Cation binding motifs (**A**) MIDAS binding motif DXSXS (**B**) Possible fifth position of MIDAS and ADMIDAS, (**C**) ITB SyMBS (green) and MIDAS (Yellow), (**D**) Possible E of SyMBS (first amino acid of SyMBS) (**E**-**F**) Cysteine rich motif, (**E**) PSI domain, (**F**) Cysteine-rich motif stalk, (**G**-**H**) C-terminal motifs (**G**) NPXY motif, (**H**) Possible KLXXXD motif

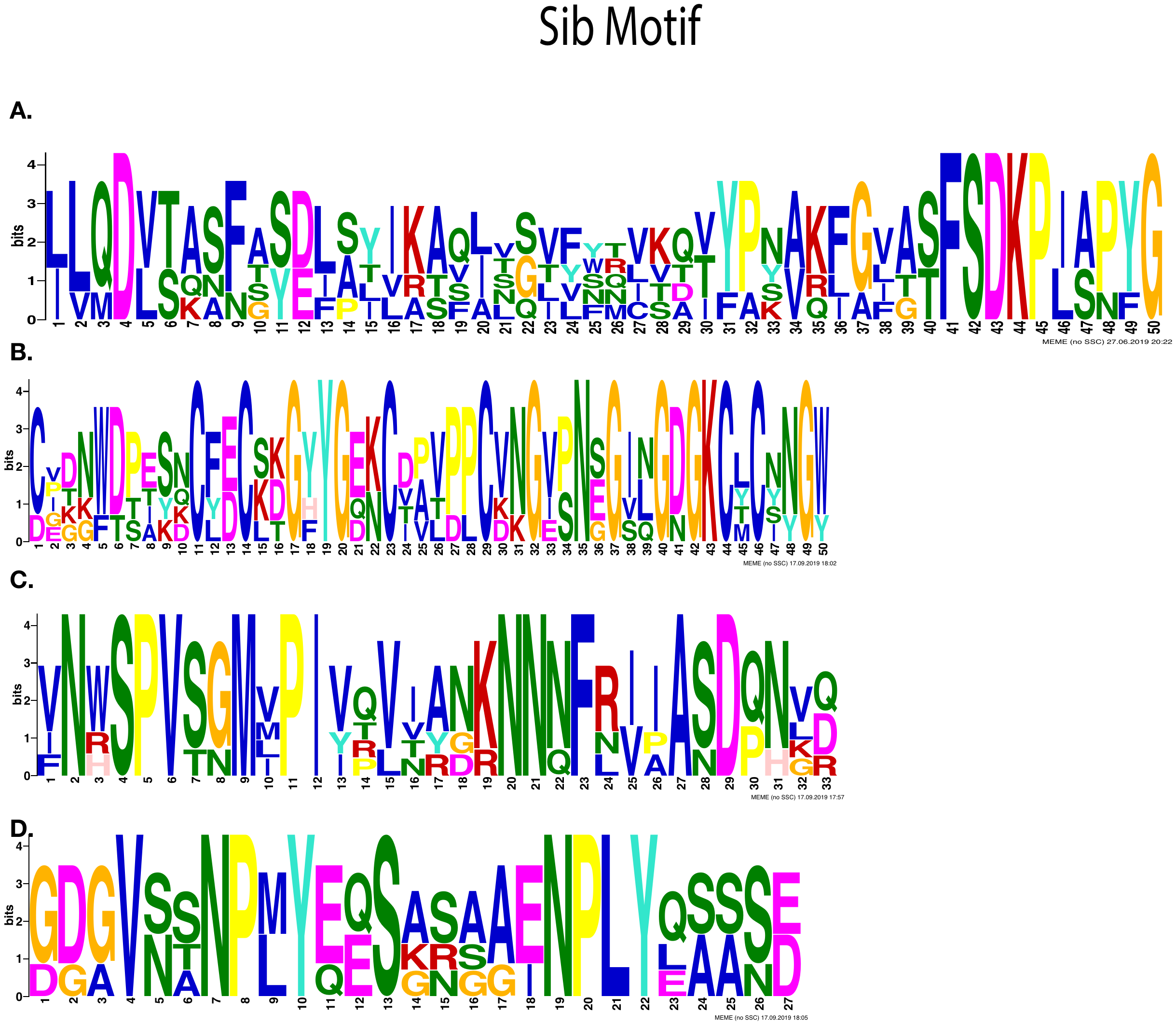
**Supplemental Figure 8:** Most significantly enriched motifs in Sib protein, obtained by MEME. (**A**)-(**D**) Cation binding motifs (A) Possible binding motif of MIDAS and ADMIDAS, (**B**) Possible fifth position of MIDAS and ADMIDAS (C) Possible binding motif of SyMBS (**D**) NPXY motif

**Supplemental Table 1:** The table shows the list of Amorphea ITGA proteins characteristic including: length, beta propeller motifs, subcellular location, any extra domains attached to ITGA, c-terminal motif signal peptide, transmembrane regions and type of ITGA. The order of taxons are listed in accordance to the Figure 2. The subcellular localization is displayed with likelihood value in parentheses. The location of extra domains attached to ITGA are shown as the C-terminal end (CTE) or at the start of N-terminal (NTB). The presence of ITGA interacting motif with ITGB is shown as GFFKR and GXXXG. The location of beta propeller motifs performed by MEME-suite is displayed. The presence of FG-GAP motifs are shown in asterixis. The location of beta motifs that corresponds to IPRSCAN output are highlighted by parentheses. The presence of consensus cation motifs are displayed in number 2 superscripts.

| Species | Size of AA | Subcellular localization  Predictability based on likelihood | Extra Domain  C-terminal end = CTE  N-terminal Beginning =NTB | GFFKR motif  GXXXG motif | Location of Beta propellor  *= predict to possess FG-GAP  ^2^ = Cation motif (D/E-h-D/N-x-D/N-G-h-x-D/E) | Signal peptide(SP) or transmembrane (TM) | Type I and II |
| --- | --- | --- | --- | --- | --- | --- | --- |
| *Filamoeba nolandi* (CAMPEP_0168571262) | 631 | Cell membrane protein Likelihood (0.971) | PRICHEXTENSN(CTE)  protein kinase (CTE) | GFFKR  GIAVG | 16, 47, [109], 171 | TM | II |
| *Ministeria vibrans* Paralog 2 (2753598) | 917 | Extracellular (0.9862)  Soluble (1)  Likelihood (0.997) | EGF 2 repeat (NT) |  | 195^2^, 255^2^, [319]^2^, [436]^2^*, [581]^2^, 772, 812^2^ |  | II |
| *Rhizamoeba saxonica* (TR11094_c0_g1_i1) | 449 | Cell membrane protein Likelihood (0.437) | None |  | [55], 122^2^, 184, 254^2^* | SP, TM | I |
| *Flabellula citata* paralog3 (c2461_g1_i2) | 673 | Cell membrane protein Likelihood (0.301) | None |  | 57*, 121, 184*, 251*, 313 | SP, TM | I |
| *Cryptodifflugia operculata* Paralog 2 (c8521_g1_i1) | 1200 | Cell membrane Protein Likelihood (0.257) | (NTB) Thrombospondin type -1 repeat, EGF 4 repeat  Intramolecular chaperone |  | 837^2^, 889^2^*, [1033]^2^, [1107]^2^ | SP | II |
| *Nolandella* sp. AFSM  (TR12097c0_g1_i1) | 975 | Cell membrane Protein Likelihood (0.672) | (NTB) Thrombospondin type -1 repeat, EGF 4 repeat |  | 613, [670]^2^*, [734]^2^*, [805]^2^*, 934 | SP | II |
| *Arcella intermedia* (c23786_g1_i1) | 1074 | Nucleus membrane Protein Likelihood (0.58) | (NTB) Thrombospondin type -1 repeat, EGF 4 repeat  Intramolecular chaperone |  | [899]^2^*, [965]^2^* | SP | II |
| *Flabellula citata* paralog1 (c9004_g2_i1) | 626 | Cell membrane Protein Likelihood (0.994) | EGF (CTE) |  | 123, [259], [323]^2^* | SP | II |
| *Flabellula citata* paralog2 (c8227_g1_i1) | 713 | Cell membrane Protein Likelihood (0.669) | EGF (CTE) |  | 88*, 150*, 255, 310 | TM | II |
| *Ministeria vibrans* Paralog 1 (2757223) | 860 | Peroxisome (0.3325), Soluble (0.9929)  Likelihood (0.386) | None |  | 58^2^, [125]^2^*, 185, 243, 310, 387^2^, 448, [578]^2^*, [647], 748 |  | I |
| *Flabellula citata* paralog5 (c8994_g2_i16) | 604 | Extracellular (0.5213)  Soluble (0.9842)  Likelihood (0.622) | None |  | 113, 172, [302]* |  | I |
| *Flabellula citata* paralog4 (c8386_g1_i3) | 646 | Cell membrane Protein Likelihood (0.833) | None |  | 53, [116]*, [178]* | SP, TM | I |
| *Cryptodifflugia operculata* Paralog 1 (c43306_g1_i1) | 581 | Cell membrane Protein Likelihood (0.445) | None |  | 58^2^, [120]*^2^, [188]*, 261^2^, 325^2^, 383*, [449]^2^ | SP, TM | I |
| *Amphizonella* sp 9 paralog2 (TR2587c1_g1_i1) | 565 | Extracellular (0.9902)  Soluble (0.9998)  Likelihood (0.997) | None |  | [47], 121, [198]^1^, 262, [344]^2^, [414]^2^, 482^2^ | SP | I |
| *Mastigamoeba balamuthi* paralog1  (MMETSP0035-201108093729) | 813 | Cytoplasm (0.3215)  Soluble (0.8872)  Likelihood (0.493) | Cadherin/[Ig-like fold](http://www.ebi.ac.uk/interpro/entry/IPR013783) (CTE) |  | 66, [200]^2^, 268, [345], 416, 468, [540] |  | I |
| *Amphizonella* sp 8 Paralog 2  (TR9459c0_g1_i1) | 569 | Vacuole (0.4562)  Soluble (0.8119)  Likelihood (0.362) | None |  | [71],138, 207, 274 [353], 419 | SP | I |
| *Heleopera sylvatica* (Gene.370_NODE_157) | 743 | Cell membrane Protein Likelihood (0.493) | ADP-ribose (NTB) | GIVAG | [56], 139, 215^2^, [280]^2^ | TM | II |
| *Nebela* sp. (Gene.16036_NODE_111) | 839 | Peroxisome (0.3381), Soluble (0.6007)  Likelihood (0.125) | Cadherin (CTE) |  | [36], [109], [179], [244], [320], [388], [455] | TM | I |
| *Difflugia* sp. D2 (DN18414_c0_g1_i5) | 266 | Organelle (0.3485) soluble (0.8094) Likelihood (0.287) | None |  | [55], [122], [190] |  | I |
| *Protosproangium articulatum* Paralog3  (TR6387\|m.15047) | 671 | Cell membrane Protein Likelihood (0.937) | None | GAVVG | [48], [124], [194] | TM | I |
| *Ceratiomyxa fruticulosa* (c6850_g1_i1) | 565 | Vacuole (0.4506)  membrane (0.7419)  Likelihood (0.215) | None |  | 74^2^, [127], 198, 259, 321, 393, 471 | SP, TM | I |
| *Diplochlamys* sp. Paralog2 (m.4379) | 666 | Cell membrane Protein Likelihood (0.989) | None | GAIAG | [120]^2^, 249, [311]^2^, | SP, TM |  |
| *Paramoeba atlantica*  (2184673) | 623 | Cell membrane Protein Likelihood (0.498) | none |  | 69, [125], 198, [337]^2^, 425^2^ | SP, TM | I |
| *Mastigamoeba balamuthi* paralog2  (MMETSP0035-20110809854) | 1042 | Cell membrane Protein Likelihood (0.674) | Cadherin/[Ig-like fold](http://www.ebi.ac.uk/interpro/entry/IPR013783) (CTE) |  | [133], 202, 265, 343,522 613, [685], 754 | TM | I |
| *Amphizonell*a sp 9 Paralog3 (Gene.962_NODE_597) | 685 | Cell membrane Protein Likelihood (0.87) | PRICHEXTENSN(CTE) |  | 68, [133]^2^, [195], 255, [287]^2^, [380]^2^*, 455, 520^2^, 586 | SP, TM | II |
| *Amphizonella* sp 9 paralog1 (TR17939c0_g1_i1) | 677 | extracellular (0.8519) soluble (0.8141) Likelihood (0.936) | None |  | [62], [124], [199]^2^, [268]*, [337], [409]^2^, [488]^2^ | SP | I |
| *Diplochlamys* sp. Paralog1 (2112184) | 540 | Vacuole (0.559)  Soluble (0.5526)  Likelihood (0.639) | None |  | [68], 143, 397, [468] | SP | I |
| *Vermistella antarctica* (156452) | 570 | Extracellular (0.7931)  Soluble (0.9906)  Likelihood (0.905) | None |  | 83, 224, 299^2^, 368, 442^2^, 516 | SP | I |
| *Protosproangium articulatum* Paralog4  (TR6387_c1_g4_i7) | 631 | Vacuole (0.4093)  Soluble (0.8173)  Likelihood (0.459) | None |  | [108], [180], 248, 329, [390]*, [487], [567]* | SP | I |
| *Amphizonella* sp 8 Paralog 1  (TR6668_c0_g1_i4) | 656 | Extracellular (0.8352)  Soluble (0.992)  Likelihood (0.964) | PRICHEXTENSN(CTE) |  | [53], [125], [200]^2^, 266^2^, 332^2^, [403]^2^, 473 | SP | II |
| *Amphizonella* sp. 10  Paralog1 (TR4717c0_g1_i1) | 563 | Cytoplasm (0.625)  Soluble (0.8464)  Likelihood (0.758) | PRICHEXTENSN(CTE) |  | 16, 87, [161], 237, 299^2^, 370, [438] |  | II |
| *Amphizonella* sp 2  (TR2129_c1_g3_i9) | 544 | Endoplasmic reticulum (0.2477)  Soluble (0.5541)  Likelihood (0.224) | none |  | 111, [189]^2^, [257], [334]^2^, [403]^2^, 474, [539]^2^ | SP | I |
| *Endostelium zonatum* LINKS (789702) | 683 | Cell membrane Protein Likelihood (0.451) | PRICHEXTENSN(CTE) |  | 232^2^, [295], 432, [497]^2^, [572]^2^, [647] | SP | II |
| *Endostelium zonatum*  (m.5907) | 681 | Cell membrane Protein Likelihood (0.479) | PRICHEXTENSN(CTE) |  | 231^2^, [293], 430, [495]^2^, [570]^2^, [645] | SP | II |
| *Protosproangium articulatum* Paralog5 (TR2944c0_g1_i1) | 735 | Extracellular (0.4086)  Soluble (0.4867)  Likelihood (0.572) | none |  | [70], [137], [200], [268], [344], 407, 480 | SP | I |
| *Protosproangium articulatum* Paralog1 (TR2074c0_g1_i2) | 851 | Cell membrane Protein Likelihood (0.448) | protein kinase | GIVFG | 57^2^, [131]^2^, [201], [275]^2^, [344], 418, 485 | SP, 2TM | II |
| *Protosproangium articulatum* Paralog2 (TR2817c1_g1_i1_6465) | 685 | Cell membrane Protein Likelihood (0.997) | None | GVGLG | [105], 176, [240], 303, [378], 450, 512 | TM | I |
| *Stenamoeba stenopodia* (2782329) | 662 | Cell membrane Protein Likelihood (0.93) | None |  | [168]^2^, 310^2^, 444^2^, 514^2^ | SP, TM | I |
| *Gocevia fonbrunei*  Paralog1 (DN14950_c7_g1_i2) | 869 | Cell membrane Protein Likelihood (0.999) | Cadherin/[Immunoglobulin-like fold](http://www.ebi.ac.uk/interpro/entry/IPR013783) (CTE) | GAIIG | [66]^2^, [207], 281, [348], 417, [488]^2^, | TM | II |
| *Gocevia fonbrunei*  Paralog2 (DN13664_c1_g1_i1) | 665 | Extracellular (0.5821)  Soluble (0.9361)  Likelihood (0.656) | None |  | 52, [121], 194, 257, [318]^2^, [386]^2^*, 463 | SP | I |
| *Amphizonella* sp. 10  Paralog2 (TR11506c6_g2_i3) | 523 | Extracellular (0.9162)  Soluble (0.9984)  Likelihood (0.929) | None |  | [56], [129]^2^, [201], [273]^2^, 340^2^, 397^2^, [468] | SP | I |
| *Ectocarpus siliculosus* | 1105 | Cell membrane Protein Likelihood (0.758) | PRICHEXTENSN(CTE) |  | [56], [127], 195, 260, 326, [396], 464* | SP, TM | II |
| *Ministeria vibrans* Paralog 3 (m.23418) | 634 | Cell membrane Protein Likelihood (0.214) | None | GFFKR | [18]^2^*, 78 | TM | I |
| *Pygsuia biforma* (ITGA) | 3312 | Extracellular (0.9162)  Soluble (0.9984)  Likelihood (0.93) | DUF11 4 repeats (CTE) | GFFKR  GLAIG | 267, [685], [807] | TM | I |
| *Lenisia limosa* (LenisMan47) | 3346 | Cytoplasm (0.5704)  Soluble (0.9615)  Likelihood (0.632) | Ig fold like 11 repeats (CTE) | GFFKR | 310^2^, [701], [826]^2^ |  | I |
| *Thecamonas trahens* | 2114 | Cell membrane Protein Likelihood (0.975) | Ig fold like 2 repeats (CTE)  Laminin_g3 (CTE) | GFFKR | 349*, [894]^2^*, [956]^2^* | SP, TM | I |

**Supplemental Table 2:** The table shows ITGB motifs, subcellular localizations, signal peptide, transmembrane and extra domains attached to ITGB protein. The order of taxons are listed in accordance to the Figure 3. For ITGB motifs of MIDAS, AMIDAS and SyMBS, any non-canonical amino acids were highlighted in red colour, cysteine rich motifs are either shown as EGF or PSI domains. The presence of C-terminal motifs are shown in KLXXXD or NPXY/F and GXXXG. The parentheses indicate number of motifs present in ITGB.

| Species | MIDAS | AMIDAS | SyMBS | CRM  Cysteine rich motifs | Size  aa | Signal Peptide and Transmembrane | B-tail motifs | Type | Subcellular localization (Deeploc) |
| --- | --- | --- | --- | --- | --- | --- | --- | --- | --- |
| *Clastostelium recurvatum* | 25D  27T  29S  123E  139R | 29S  32S  33D  202A  160D | 81E  120R  121P  122D  123E | None | 1405aa | TM | GAVVG | II | Extracellular  Likelihood (0.3436)  soluble likelihood (0.731) |
| *Filamoeba nolandi* | 893D  895Y  897S  956E  991D | 897S  899K  900A  1066A  991D | 906D  949D  950E  951P  952N | EGF, PSI (NTB) | 2714aa | SP, TM | GAFIG | II | Cell membrane Protein likelihood (0.5463) |
| *Soliformovum irregularis* | 872D  874S  876N  958S  975N | 874N  879Y  880N  1071A  975N | 924D  1012D  1013P  1014N  1015E | PSI | 2218aa | SP | NPXY Present | II | Extracellular  likelihood (0.8972) soluble likelihood (0.9172) |
| *Schizoplasmodiopsis vulgaris* | 839D,  841T  845S  928E  944D | 845S  846Y  857E  1045A  944D | 889D  923N  925Q  927L  928E | PSI | 3641aa | SP, TM | NPXY present  GAAG | II | Extracellular  likelihood (0.5465) soluble likelihood (0.6244) |
| *Cavostelium apophysatum* | 507D  509T  511S  592N  642D | 511S  514S  515D  708A  642D | 546D  656N  658D  659E  660D | PSI | 3296aa | TM | NPXY present  GAASG | II | Cell membrane protein likelihood (0.7274) |
| *Mycamoeba*  *gemmipara* | 277D  279T  281S  375E  416E | 281S  285N  286D  530A  390D | 328E  403N  411D  414P  416E | Unique cysteine rich region pattern downstream from 500AA-860AA | 927aa | TM | NPXY present  GVIAG | I | Cell membrane protein likelihood (0.998) |
| *Mastigamoeba balamuthi* | 159D, 161T, 163S 242E  271D | 163S,  170P,  171N  371S,  271D | 198E,  238S,  239D,  241P,  242E | PSI, EGF-like present 5 times | 1315 aa | SP, TM | NPXA present  GAASG | I | Cell membrane Protein likelihood (0.561) |
| *Mastigella eilhardi* | 127D, 129S, 131S 212E  230D | 131S  134D  135D  339A  230D | 198E,  207N,  209D,  211P,  212E | PSI, EGF-like present 5 times | 1292 aa | SP, TM | NPXA present  GAASG | I | Cell membrane Protein likelihood (0.7939) |
| *Rigifila limosa* | 124D  126S  128S  206E  237D | 128S  131D  133D  391A  237D | 164D  201N  203D  205P  206E | PSI, EGF-like present 13 times | 932 aa | SP, TM | NPXY present  GTATG | I | Extracellular soluble likelihood (0.681) |
| *Echinamoeba exudans* | 123D, 125S, 127S, 209E,  239D | 127S,  140G  141E  338A  239 D | 174E  204N,  206E,  208P,  209E | PSI, EGF-like present 3 times | 593 aa | SP, TM | NPXY present (2)  KMAAAD  GAVIG | I | Cell membrane protein likelihood (0.5596) |
| *Difflugia* sp. D2 | 115D, 117S, 119S,  213E,  249D | 119S  136D  137E  295I  249D | 177E,  208N,  210E,  212P,  213E | PSI, EGF present 3 times | 605 aa | TM | NPXY present (2) | I | Cell membrane protein likelihood (0.8598) |
| *Nebula* sp. | 131D,  133S, 135S,  225E,  264D | 135S  138P  139Y  361A,  264D | 192E,  223N,  225E,  227P,  228E | PSI, EGF present 3 times | 630 aa | SP, TM | NPXY present (2) | I | Cell membrane protein likelihood (0.7413) |
| *Heleopera sphagni* | 127D  129S  131S  221E  266D | 131S  134P  135H  352A  266D | 174E  216N  218E  220P  221E | PSI, EGF present 3 times | 622 aa | SP, TM | NPXY present (2)  KADX(4)E | I | Cell membrane protein likelihood (0.9332) |
| *Difflugia* sp. USP | 146D,  148S,  150S,  245E  280D | 150S  153P  154Y  370A  280D | 197E  239N  241E  243P  245E | PSI, EGF present 3 times | 630 aa | SP, TM | NPXY present (2) | I | Cell membrane protein  likelihood (0.9847) |
| *Hyalosphenia papilio* | 35D  37S  39S  132E  172D | 39S  42P  43Y  270D  172D | 85D  127N  129E  131P  132E | EGF present 3 times | 531 aa | TM | NPXY present (2)  KADX(4)D | I | Cell membrane protein likelihood (0.8186) |
| *Rhizamoeba saxonica* (paralog 1) TR11158c0_g1_i1 | 143D  145S  147S  228E  259D | 147S  150Q  151E  383A  259D | 182E  223N,  225D,  227P,  228E | PSI, EGF present 7 times | 951 aa | SP, TM | NPXY present (2) | I | Cell membrane protein likelihood (0.7754) |
| *Cryptodifflugia operculum* | 144D  146S  148S  241E  278D | 148S  151P  152L  370G  278D | 194E  236N  238E  240P  241E | PSI, EGF present 3 times | 633aa | SP, TM | NPXY present (2) | I | Cell membrane protein likelihood (0.9426) |
| *Rhizamoeba Saonxica* (Paralog2) TR7856c0_g1_i1 | 142D,  144S  146S  227E,  258D | 145S  148D  149D  329A  258D | 181E,  222N,  224D,  226C,  227E | PSI, EGF present 3 times | 932aa | TM | NPXY present (2) | I | Cell membrane protein likelihood (0.8204) |
| *Flabellula citata* | 163D,  165S  167S  234E,  282D | 167S  172M  173E  364A  282D | 185D,  229G  231D,  233P  234E | PSI, EGF present 5 times | 1141 aa | SP | GAAVG | I | Cell membrane protein likelihood (0.7827) |
| *Rhizamoeba saxonica*  (Paralog 3) TR7441_c0_g1_i1 | 832D  834S  836S  918E  949D | 836S  839P  840F  1063A  949D | 871E,  913N  915D  917C  918E | EGF-like present 2 times | 1456 aa | SP, TM | NPXY present (2) | I | Extracellular (05263) soluble likelihood (0.731) |
| *Pyxidicula operculata* | 133D, 135S, 137S,  229E,  264D | 137S,  140P  141Y,  360A,  264D | 192E,  224N,  226E,  228P,  229E | PSI, EGF present 3 times | 621 aa | SP, TM | NPXY present  KADX(4)D  GECGG | I | Extracellular (0.9867) soluble likelihood (0.9995) |
| *Pygsuia biforma* | 110D  112S  114S  204E  231D | 114S  117D  118D  331A  231D | 161D  200Q  201D  203P  204E | PSI, EGF present 34 times | 1793 aa | TM | NPXY present KAGX(4)E present | I | Cell membrane Protein likelihood (0.9792) |
| *Lenisia limosa* | 130D,  132S 134S  218S  246D | 134S  137N  138D  357N  246D | 182N  215N  216E  217P  218S | PSI, EGF present 12 times, EGF like 4 times |  | SP, TM | NPXY present  RFNAAMKE present | I | Cell membrane Protein likelihood (0.9934) |
| *Ministeria vibrans (paralog 1)* c191\|m.702 | 85D,  87S  89S  179E  210D | 89S  92D  93D  310S  210D | 124E  174N  176D  178P  179E | EGF-like present 5 times |  | SP, TM | NPXY present  KAWQQWEE | I | Cell membrane protein likelihood (0.9827) |
| *Ministeria vibrans (paralog 2) c46_198* | 73D,  75S  77S  194E  225D | 77S  80N  81D  339A  225D | 122D  189N  191D  193P  194E | EGF-like present 14 times |  | TM | NPXF present (2)  KLAQVAAD | I | Cell membrane Protein likelihood (0.9941) |
| *Ministeria vibrans (paralog 3)*  c334\|m.1235 | 594D,  596S 598S  682E  714D | 598S  601N  602D  824A  714D | 633E  677N  679H  681S  682E | EGF-like present at N-terminal and C-terminal |  | SP, TM | NPXY present (2)  KLITGAAD | I | Cell membrane Protein likelihood (0.9985) |
| *Didymoeca costata* | 444D  446S,  448S  530E, 562D | 446S  449D  451D  652A  562D | 524S  527D  529P  530E | None | 1673aa | SP, TM | None | I | Cell membrane protein likelihood (0.7458) |
| *Sphaeroforma arctica* | 119D  121S  123S  231E  260D | 123S  125D  127D  296A  260D | 195E  227N  229D  230P  231E | EGF-like present  7 times | 1141aa | TM | KGIQMVMD | I | Cell membrane protein likelihood (0.638) |

**Supplemental Table 3:** This table shows Rsem value of protists that have either a complete set of IMAC, integrins with extra domains or unexpected discovery of an integrin protein. The values are shown in transcripts per million (TPM).

| Species | ITA | ITB | Talin | PINCH | ILK | Paxillin | Actinin | Actin | eF1α | Vinculin |
| --- | --- | --- | --- | --- | --- | --- | --- | --- | --- | --- |
| *Filamoeba nolandi* | 10.23 | 91.07(?) | 45.54 | 30.62 | 11.71 | 0.00 | 138.81 | 2417.45 | 5932.12 | 36.07 |
| *Difflugia* sp. D2 | 0.00 | 0.00 | 0.00 | 6.01 | N/A | 0.00 | 63.16 | 12.24 | 0.00 | 0.00 |
| *Flabellula citata* | 16.31(Paralog1)  9.81(Paralog2)  14.56(paralog3)  4.90 (Paralog4)  0.67(paralog 5) | 421.32 | 18.56 | 34.78 | 5.54 | 25.09 | 109.90 | 2708.65 | 6180.65 | 7.28 |
| *Nebela* sp. | 3.73 | 11.34 | 3.95 | 85.57 | N/A | 25.24 | 444.60 | 653.00 | 2388.17 | 6.42 |
| *Rhizamoeba saxonica* | 174.00 | 52.55(Paralog3)  82.61(Paralog2)  121.63(paralog1) | 25.63 | 29.96 | 3.23 | 13.13 | 106.07 | 12079.17 | 5322.57 | 20.45 |
| *Cryptodifflugia operculata* | 4.93 (paralog1)  1.80 (paralog2) | 7.95 | 50.58 | 2.40 | 2.47 | 15.50 | 5.55 | 272.10 | 1857.80  (c7512_g1_i1) | 9.65 |
| *Nolandella* sp. AFSM | 2.77 | N/A | 9.84 | 41.65 | N/A | 11.66 | 3.44 | 2929.71 | 3621.35 | 12.38 |
| *Arcella intermedia* | 2.98 | N/A | 4.91 | 1.19 | N/A | 11.19 | 43.20 | 110.29  (c17090_g2_i1) | 2001.59 | 4.71 |
| *Rigifila ramosa* | N/A | 810.35 | 348.93 | 59.65 | N/A | 141.87 | 127.94 | 18519.18 | 2284.60 | 7.37 |
| *Pygsuia biforma* | 2.14 | 1.45 | 41.95 | 111.50 | N/A | 13.47 | 103.13 | 17397.16 | 7071.38 | N/A |
| *Ministeria vibrans* | 1.71 (Paralog 1)  2.90 (Paralog 2)  20.02 (Paralog 3) | 0.00(Paralog 1)  5.17(Paralog 2)  10.56 (Paralog3) | 3.88 | 13.43 | 3.38 | 13.43 | 8.72 | 277.70 | 310.59 | 1.15 |

**Supplemental Table 4:** A). Summary of amoebozoan ITA type I table listing: number of paralogs, Beta propeller, FG-GAP and canonical motifs. The table also shows presence of signal peptide and transmembrane region, c-terminal interacting motif, GXXXG motif and Ig-fold like/Cadherin domain. B). Summary of amoebozoan ITA type II table listing: number of paralogs, Beta propeller, FG-GAP and canonical motifs. The table also shows presence of signal peptide and transmembrane region, c-terminal interacting motif, GXXXG motif and extra domain attached to ITA.
